# Bayesian decision theoretic design of two-founder experimental crosses given diallel data

**DOI:** 10.1101/489682

**Authors:** Gregory R. Keele, Paul L. Maurizio, Daniel Oreper, William Valdar

## Abstract

In designing experimental crosses of inbred strains of model organisms, researchers must make a number of decisions. These include the selection of the appropriate strains, the cross design (*eg*. F2 intercross), and the number of progeny to collect (sample size). These decisions strongly influence the potential for a successful quantitative trait locus (QTL) mapping experiment; good design decisions will lead to efficient and effective science. Thus experimental design deserves careful consideration and planning. Experimental outcomes can be quantified through utility functions using a Bayesian decision theoretic approaches. For QTL mapping experiments, the power to map a QTL is an appealing utility function to maximize. Using any utility function to aid in experimental design will be dependent on assumptions, such as the QTL effect size in the case of power. Rather than arbitrarily selecting QTL effect size values, they can be estimated from pilot data using a Bayesian hierarchical model. The information in the pilot data can be propagated to the utility function, using Markov Chain Monte Carlo (MCMC) to sample from the posterior distribution. Key features of this approach include: 1) distributional summaries of utility, which are preferable to point estimates, and 2) a comprehensive search of the experimental space of crosses of inbred lines for well-designed experiments. We evaluate this Bayesian theoretic approach using diallel crosses as the pilot data. We present results from simulations as well as present examples from both Mendelian and complex traits in the founder strains of the mouse Collaborative Cross. All analyses were performed using our R package, DIDACT (Diallel-Informed Decision theoretic Approach for Crosses Tool), developed to perform Bayesian cross selection based on diallel pilot data.

## Introduction

Geneticists commonly conduct experiments with the goal of identifying quantitative trait loci (QTL) using crosses of inbred lines of model organisms. These experiments can be costly in terms of resources, due to the organisms, their care, genotyping or sequencing’ as well as the time and energy required for the experiment itself. In the face of these constraints, procedures that explore the potential set of experimental cross designs and allow researchers to select experiments with greater potential to be successful are beneficial to the field of complex traits.

Although the goals for a given experiment will be nuanced and unique to each study, the mapping portion is successful if a QTL is detected with a statistically significant signal, using established methodologies (Lander and Botstein 1989; Haley and Knott 1992; Dupuis and Siegmund 1999; Broman 2001). This outcome is not guaranteed simply due to the presence of segregating QTL in the mapping population: the experimental design may not be sufficiently powered to identify them. The power of an experiment, the probability that a non-zero effect will be recognized given that it is present, is influenced by a number of biological factors, some of which can be more easily manipulated and optimized through experimental design choices. These factors include genetic architecture, mode of action, and the variation in the population due to noise. If the genetic architecture of the trait is highly polygenic with many loci of small effect, power will be reduced compared to tests for QTL of larger effect. Similarly, mode of action (e.g. additive, dominant), for a QTL will also influence power because certain experimental designs will have differential ability to detect a given effect type. For example, a backcross (BC) cannot identify a QTL with a fully recessive effect when the homozygote of the recessive allele is never observed. Finally, an increase in variation due to noise will decrease power because the noise drowns out the true signal. Ideally, investigators would select the experiment that can best handle these factors in the given setting.

In the context of crosses of inbred organisms, one major component of the experimental design is the founder or parental lines. The selection of parental lines allows the investigator to control the genetic background of the experimental population, which can greatly influence the previously mentioned biological factors, and ultimately influence the potential for mapping success. For example, a trait could be highly polygenic and have loci with complex modes of action within natural populations, but much of the genetic and phenotypic variation becomes fixed within two closely related inbred lines. The reduced genetic variability can impact all of the biological factors: the complexity or polygenic nature of the genetic architecture by fixing many of the loci, the mode of action by limiting the potential for epistatic effects through less segregating variants, and the variance attributable to noise through the reduction in phenotypic variability.

The ability to strongly influence the sources of variation in the population is important to consider. If the QTL explains a large proportion of the variance in the population, a simple cross will be well-powered to identify the QTL, even if its effect is small. The balance between the variance attributable to the QTL versus how generalizable the experiment is to natural populations is important to consider when making decisions about experimental design. Ultimately a finding that is characteristic of only a very unnatural experimental population and does not generalize well to more natural ones, will greatly reduce the impact of such an experiment and even undermine the purpose of experiments with model organism in general. The ideal experiment will be well-powered to identify QTL, but also generalizable to natural populations.

The power of an experiment cannot be directly assessed because it requires knowledge of the true effect, which is unknown. Instead power calculations are performed for a range of plausible parameters, usually over varying effect sizes or sample sizes, given some type I error level and error variance, which can then be represented as power curves. Analytical solutions to power calculations have been specifically developed and refined for simple cross designs such as F2 intercross, BC, and recombinant inbred lines (RIL) panels, using an information perspective approach, which posits that the complete information is composed of the unobserved information and the missing information (Sen *et al*. 2005). These power estimates are still dependent on assumed parameters, in this case QTL effect sizes and error variances. As a result, meaningful and useful power calculations still depend on the consideration of an appropriate set of values for these unknown quantities, otherwise the power estimates could be uninformative or even misleading.

Pilot data can provide information about the underlying genetic signals present in potential experiments. One source of pilot data is the inbred founder lines themselves as well as their hybrid crosses (F1). Comparisons of F1 individuals to the inbred strains can provide estimates of various genetic effects for given lines, aggregated from causal variants across the entire genome. These effects can include additive, inbred, and epistatic. An additive effect for a given strain can be estimated from averages of F1 that do not have the strain as a parent (0 copies), to averages of F1 that do have the strain as a parent (1 copy), and finally to the inbred line itself (2 copies). An inbred effect is estimated from these same sets of crosses, but represent the average departures observed from the expectation of the hybrid according to the additive effect to its actual observed value. An epistatic effect represents departures from expectation for a specific cross of two strains, thus it is an interaction of the two strains.

Additional information is contained in the reciprocal crosses that compose the F1 hybrids, and can be characterized as parent-of-origin effects (POE). Reciprocal F1 crosses have the same parental lines, but the dam-sire identities are switched. Average differences between reciprocal crosses can be used to estimate POE, allowing QTL that underly these POE to be mapped using a unique BC design that we will refer to as reciprocal parent-of-origin (RBC^PO^) (Gonzalo *et al*. 2007). Traditional reciprocal BC refer to related BC in which F1 are back-crossed to alternative parental lines, whereas RBC^PO^ have the same F1 and back-crossed parent, but the dam and sire strains are reversed between reciprocal pairs; thus the parent-of-origin for each allele is known at heterozygous sites, and differences in the trait that correlate to genotype and parent-of-origin can be detected. The estimation of POE through reciprocal crosses allows researchers to add RBC to their collection of potential experiments. Though RBC^PO^ are not as frequently used as F2 and BC, interest in POE has increased (Lawson *et al*. 2013; Bérénos *et al*. 2014; Connolly and Heron 2015; Harper *et al*. 2014; Zou *et al*. 2014). Pilot data that distinguish between reciprocal F1s allow for an even larger number of experiments to be explored and considered. These potential bi-parental mapping populations, F2, BC, and RBC^PO^, are depicted in Figure 1.

**Figure 1.**
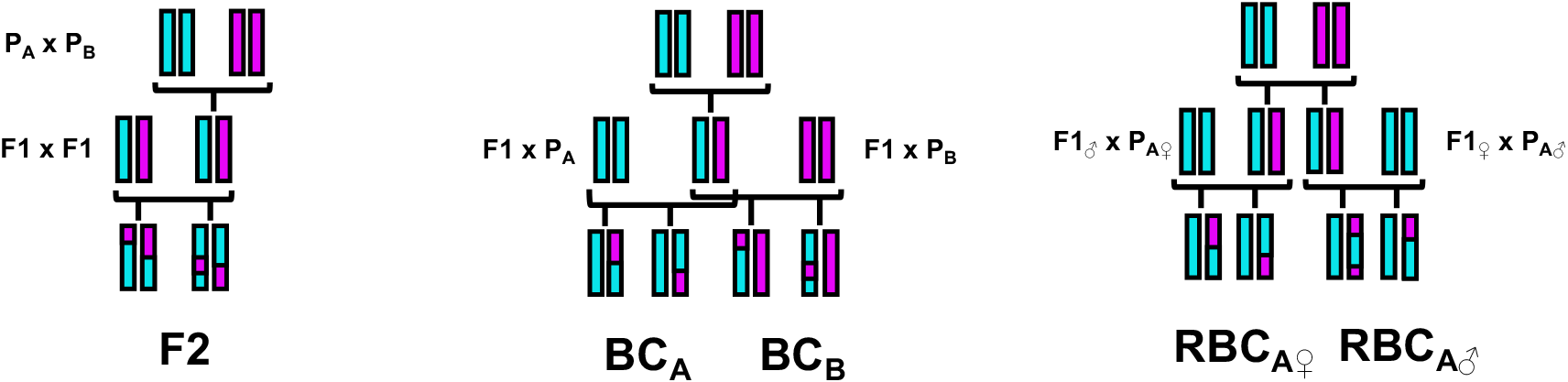
The potential bi-parental crosses considered within DIDACT: F2 [left], BC [middle], and RBC^PO^ [right]. A single inbred genome is represented as a colored chromosome. The parental (P) and F1 generations are replicable, whereas the mapping populations are not. All three genotypes at a locus (A/A, A/B, and B/B) can be observed in the F2 population, allowing for the estimation of additive and dominance effects for a putative QTL. With traditional BC, only the homozygote of the backcrossed parental allele is observed. By jointly analyzing RBC^PO^, it is possible to detect effects from heterozygous sites in which the parent-of-origin differs for the backcrossed parental allele.

These experiments can best be explored with the full set of potential founder lines and their F1 hybrids, which represent a classic genetic experiment, the diallel cross. Diallels have been used to study numerous traits in a diverse range of organisms, including: mating speed, female receptivity, and temperature preference in flies (Parsons 1964; Casares *et al*. 1992; Yamamoto 1994); immune function, polyandry, and genetic-environment interactions in crickets (Rantala and Roff 2006; Ivy 2007; Nystrand *et al*. 2011); and heterosis and reciprocal effects in poultry (Fairfull *et al*. 1983). Additionally, the diallel has a long history in plant breeding (Gilbert 1958) and numerous recent applications (Bahari *et al*. 2012; Ghareeb Zeinab and Helal 2014; Dos Santos *et al*. 2016).

Since being described in the early 20^th^ century, statistical methodology for the diallel has seen steady advancements, from estimating the general combining ability of related F2 populations (Griffing 1956) to random effect (Zhu and Weir 1996; Tsaih *et al*. 2005) and Bayesian hierarchical modeling of sparse diallel (Greenberg *et al*. 2010). Recently, Lenarcic *et al*. (2012) used Bayesian hierarchical modeling of diallel data to allow for stable estimation of a large number of strain-level genetic effects (such as additive, inbred, epistatic, and maternal), and this method has been used to analyze a number of phenotypes and organisms, such as cranial shape (Gonzalez *et al*. 2016), response to treatment and infection (Crowley *et al*. 2014; Maurizio *et al*. 2018), and litter size (Shorter *et al*. 2018) in mice, and shoot growth in carrots (Turner *et al*. 2018). Even incomplete or sparse diallel data can be used for the characterization of some of the underlying strain-level genetic signals, which can then be used to evaluate the potential space of experiments, and potentially allow for the selection of a favorable one. A simplified representation of a diallel, in the founders of the Collaborative Cross (CC), a multiparental recombinant inbred panel in laboratory mouse, is depicted in Figure 2.

**Figure 2.**
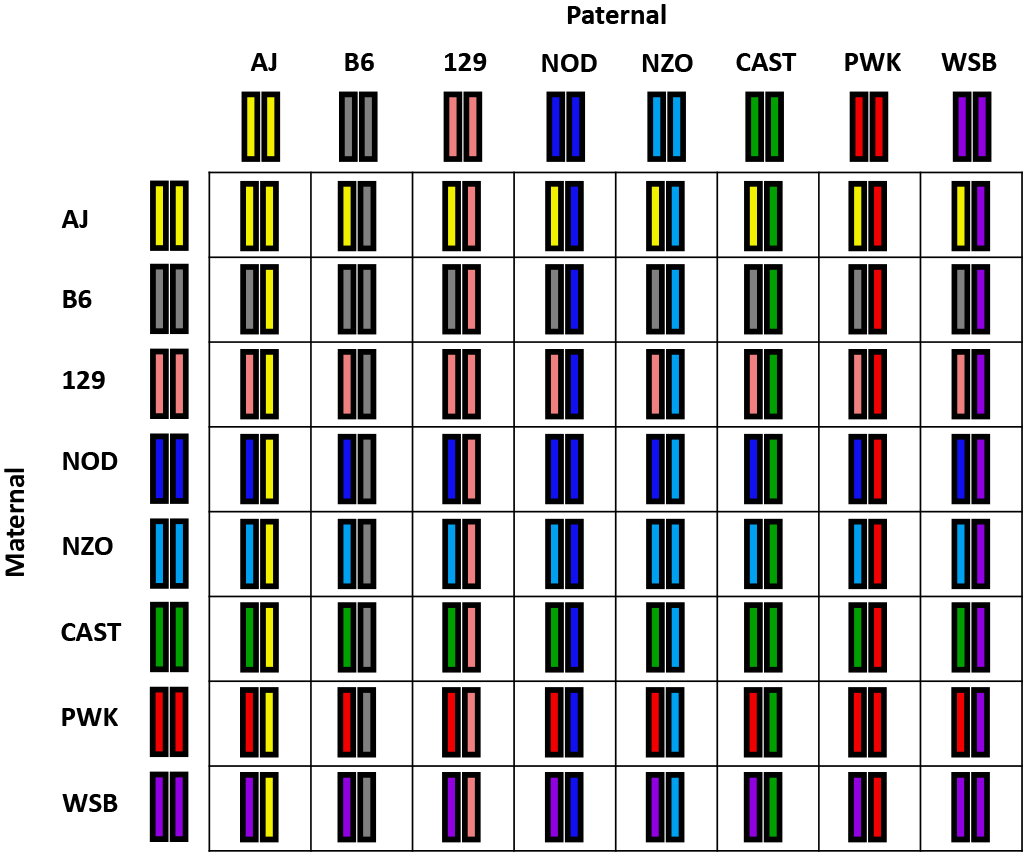
A diallel of the CC founders. Each unique strain genome is represented as a colored chromosome. Genomes along the diagonal represent the inbred founders. Off-diagonal genomes represent the F1 hybrids of a pair of founders. Reciprocal F1 genomes possess the same genome, ignoring mitochondria and sex chromosomes for males, but parent-of-origin for each chromosome will be reversed. All the genomes of an inbred diallel cross are replicable, and thus replicate observations can be measured per genome. Some cells of the diallel may not be observed, reducing the ability to accurately estimate certain strain-level effects.

A number QTL mapping approaches have been developed within the context of jointly modeling related populations, including daughter populations of a diallel cross (Rebaї and Goffinet 1993, 2000; Liu and Zeng 2000; Li *et al*. 2013), as well as QTL allele effect estimation for these related populations (Jannink and Wu 2003). More recently, Verhoeven *et al*. (2006) investigated jointly modeling diallel data with the related downstream F2 populations, and found that it allowed for the simultaneous dissection of the trait across all the populations, or characterization of strain-level effects, as well as generalization of the QTL findings from the mapping populations in terms of the multiparental diallel population. We focus on the situation in which none of the F2 populations, or any such downstream cross populations, are observed, and attempt to evaluate the utility of potential crosses in terms of QTL mapping. Herein we bring together three lines of research:

1. The estimation of the power to map putative QTL of given effect sizes.
2. The characterization of strain-level genetic effects from diallel pilot data.
3. The selection of optimal experiments through a decision theoretic approach.

We use the BayesDiallel model (Lenarcic *et al*. 2012), a Bayesian hierarchical model for characterizing the genetic information contained in diallel data as aggregate strain effects. Bayesian approaches can stably estimate a large number of genetic effects through the sharing of information across strains, as well as assess the uncertainty around these effects. These strain effects are next propagated to utility functions, including power to map a putative QTL underlying the strain effects, for an array of potential experimental crosses. Our approach will allow researchers to select better experiments with greater potential based on pilot data over ineffective or inefficient options. These opportunities include not only favorable experiments for mapping additive traits, which have commonly been studied, but also for mapping the QTL responsible for less well-understood effects such as POE.

## Statistical Models and Methods

Our approach builds on three separate areas of research. Firstly we consider the calculation of power to map QTL given that the QTL effect *θ* is known. This will require the review of general concepts in quantitative genetics and statistics in the context of crosses of two inbred lines. Because in reality *θ* is never actually observed, we next consider the characterization of *θ* from pilot data. Finally we discuss the selection of optimal experimental crosses through the maximization of a chosen utility function.

### Power to map QTL

#### Single QTL model

Here we review the general concepts in quantitative genetics and statistics that support the method used by Sen *et al*. (2005) for power calculations of traditional crosses such as the F2 and BC. Consider this model:

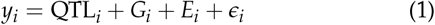

where *y_i_* is the phenotype of individual *i*, QTL_*i*_ is the effect of the QTL for individual *i, G*_*i*_ is the effect of other genetic elements for individual *i, E*_*i*_ is the effect of environmental factors for individual *i*, and *ϵ*_*i*_ is the random noise for individual *i. G*_*i*_ and *E*_*i*_ are un-modeled, and can thus be collapsed with *ϵ*_*i*_ into a single error term *ε*_*i*_.

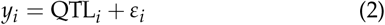

where *ε*_*i*_ ~ N(0, *σ*^2^) with *σ*^2^ representing the error variance in the data. The QTL effect is a vector, traditionally parameterized as additive and dominant effects (Lynch and Walsh 1998). This can be formulated in a traditional regression framework:

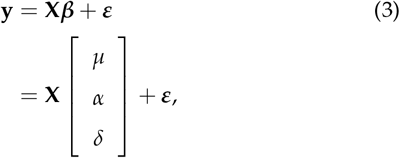

where **y** is the phenotype vector, **Χ** is the design matrix that we will define further, ***β*** is the vector of effects composed of *μ*, the overall phenotypic mean, *α*, the additive effect of the QTL, and *δ*, the dominance effect for the QTL, and ***ε*** is the vector of errors. Consider an F2 or BC of strains *A* and *B*, with the genotype of an individual represented in terms of strain identity, denoted in the subscript. *α* is the midpoint of the difference between the homozygotes:

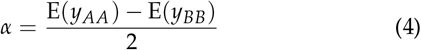

*δ* is the deviation of the heterozygote from the average of the homozygotes:

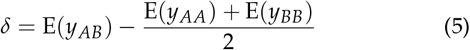

**Table S1** lists Eq 3 parameterized in terms of these QTL effects. This parameterization maintains the identifiability of all the effects, though it may not be as intuitive to researchers accustomed to more traditional regression models used commonly in genome-wide association studies.

Returning to the formulation of the model in Eq 6, the variance of the model can be characterized as follows with the assumption that there is no covariance between the QTL effect and the error,

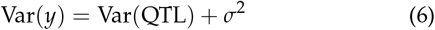

The background genetic and environmental variation are captured in *σ*^2^; here we focus on the variability due to the QTL. E(*y*) will vary depending on the genotype, which will vary probabilistically according to the type of cross, as described in **Table S1**. As example, for an F2 cross, the 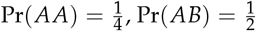, and 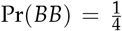. The variance of a random variable *X* is defined as Var(*X*) = E(*X* – E(*X*))^2^. The variable *X* in this setting is QTL, which is the categorical genetic state at the QTL. The expectation of *X* is 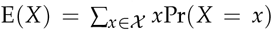. Based on the genotype probability for a given cross, the variances due to the QTL in terms of the QTL effects are presented in **Table S2**.

The mode of action of the locus impacts the variability in phenotype due to QTL within a cross type, as seen in **Table S2**. This is particularly noticeable in the BC experiments, where certain modes of action produce no variance. If the locus is recessive (or conversely dominant), the genotype with differing phenotype will not be observed, and nor will variation due to QTL. Finally, cross type also impacts the QTL variance, which is also clear in **Table S2**. Increasing the variance attributable to the QTL will increase power to map the QTL; in contrast, increasing the overall variance that is attributable to noise (un-modeled background genetic factors or environmental factors) will reduce the significance of statistical tests, and thus decrease the power.

#### Power calculations

Analytical power calculations are the probability mass above some threshold for the distribution of a statistic of interest under the alternative hypothesis. This requires that the alternative distribution be reasonably characterized. Consider *θ*, some function of the QTL effects *α* and *δ*, as the parameter of interest. We wish to calculate the probability of mapping the QTL that results in *θ*. In terms of the association modeling, a natural null hypothesis is H_0_: *θ* = *θ*_0_ with *θ*_0_ = 0, that there is no QTL effect. The alternative hypothesis is *H*_*A*_: *θ* ≠ *θ*_0_. By specifying a model for the data, or more precisely the distribution of the error term of the model, the likelihood 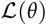 can be evaluated. The likelihood ratio test (LRT) statistic, 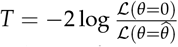, where 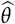 is a proposed estimate of *θ*, can be used to perform power calculations.

To use the LRT statistic for power calculations, a significance threshold and corresponding statistic distribution for *T* are necessary. The traditional scale of significance used in the linkage and QTL fields is the log_10_ likelihood ratio or LOD (logarithm of odds) score. Historically a LOD score of 3 (2log(10) × 3 on the likelihood ratio scale) has been used as a significance threshold, meaning approximately that the data support the alternative model over the null model 1000 to 1. A more stringent significance threshold than 3 can be used to further reduce the risk of false positives or possibly account for a multitude of tests (though it is worth noting these tests will not be fully independent). Given some significance threshold *C* is chosen to determine genome-wide significance; if *T* ≥ *C* for some locus, the null hypothesis is rejected. The threshold *C* will affect the the true positive and false positive rates, and more important to our topic, the power.

Statistically, power is the probability that the null hypothesis is rejected given that alternative hypothesis is true. The LRT T is the statistic upon which the power calculations are drawn, thus the power will be Pr(*T* ≥ *t*|*θ* ≠ *θ*_0_) where *t* is the observed statistic produced by the data. With the LRT statistic, when the models are nested and the maximum likelihood estimate (MLE) is used (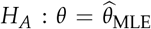), as they are in this case, and the null model is true, *T* is asymptotically 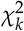 distributed, where *k* is the degrees of freedom, the difference in number of parameters between the models. A power calculation from this distribution would not be useful because it would represent the probability that the null hypothesis is rejected when there is no genetic effect, or the false positive probability. The power is rather based on the alternative hypothesis being true, *θ* ≠ *θ*_0_, and thus 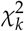 distribution is inappropriate. When the alternative hypothesis is true rather than the null, that 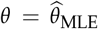, *T* is proportional to the noncentral *χ*^2^ distribution with noncentrality parameter 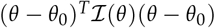 where 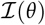 is the expected Fisher information matrix. We model the data with a Gaussian mixture distribution with a shared residual variance, which naturally extends from the two line cross statistical model. A key feature of this model is that the LRT reduces to the variance attributable to the QTL as a function of effects that we presented in table 1. This variance parameter is scaled by *σ*^2^, which sets the variance of each Gaussian component to 1. Thus the power calculations are intuitively a function of the effect size, the proportion of the variance explained by the QTL (effect size combined with residual error variance), and the sample size.

**Table 1.**
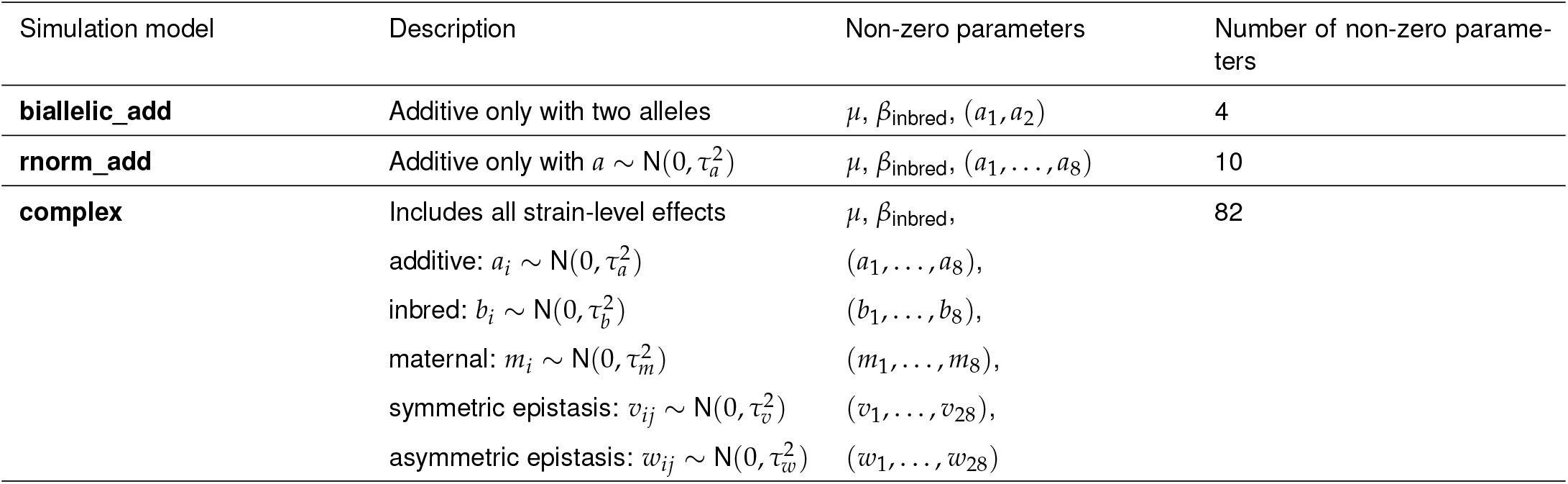
Summary of data-generating models for simulations.

| Simulation model | Description | Non-zero parameters | Number of non-zero parameters |
| --- | --- | --- | --- |
| **biallelic_add** | Additive only with two alleles | $\mu, \beta_{\text{inbred}}, (a_1, a_2)$ | 4 |
| **rnorm_add** | Additive only with $a \sim N(0, \tau_a^2)$ | $\mu, \beta_{\text{inbred}}, (a_1, \dots, a_8)$ | 10 |
| **complex** | Includes all strain-level effects | $\mu, \beta_{\text{inbred}},$ | 82 |
| | additive: $a_i \sim N(0, \tau_a^2)$ | $(a_1, \dots, a_8),$ | |
| | inbred: $b_i \sim N(0, \tau_b^2)$ | $(b_1, \dots, b_8),$ | |
| | maternal: $m_i \sim N(0, \tau_m^2)$ | $(m_1, \dots, m_8),$ | |
| | symmetric epistasis: $v_{ij} \sim N(0, \tau_v^2)$ | $(v_1, \dots, v_{28}),$ | |
| | asymmetric epistasis: $w_{ij} \sim N(0, \tau_w^2)$ | $(w_1, \dots, w_{28})$ | |

It is important to note that the actual 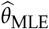 cannot be calculated because no actual cross data for QTL mapping is observed, but the underlying theory of the method assumes that the alternative *θ* is the MLE estimator. *σ*^2^ is also never actually known, but we estimate it from the information present in the pilot data. The final interpretation of this power calculation is the probability that a significant result is found (*T* ≥ *t*) given that there is some QTL effect specified in the proposed MLE estimator 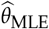 with an error variance of *σ*^2^.

Sen *et al*. (2005) develop the theory further to account for the fact that the information is generally never complete in QTL studies. The true QTL variant is most likely not observed (genotyped), but rather loci in linkage disequilibrium are, and thus contain some of the information from the QTL. They develop the theory to take into account the missing information from sparse markers (as previously described), as well as selective genotyping (genotyping study individuals on the tails of the phenotype distribution). As a result of this, power can be reduced by not only greater error variance, but also missing information. The advancement in genotyping technology is generally leading to denser markers in QTL studies, leading us to make the assumption of complete information. We directly incorporate the R package qtlDesign (Sen *et al*. 2007) into our method, so missing information can be specified in the power calculations. See Sen *et al*. (2005) for a description of the missing information theory used.

### Strain-level genetic effects

The power calculations described above are dependent on known QTL effects *θ*, but in reality, *θ* is not observed. However, information about *θ* is contained in pilot data, which can be exploited to characterize plausible distributions for *θ*.

#### Bayesian modeling of diallel data

One potential convenient source of pilot data are the parental lines and some subset of their F1 hybrids. Direct estimation of *θ* is not possible because no recombinations occur between the parental haplotypes within F1 individuals, but rather strain effects that represent the accumulated effect of the segregating variants within each inbred strain can be estimated. Denote these strain effects, the vector of effects that will be defined in Eq 7, as ***ϕ*** to distinguish them from *θ*, the effect of a single QTL.

The strain-level vector ***ϕ*** can encompass effects of different modes of actions based on the strain identities of the dam and sire of an individual. These strain-level effects include additive, inbred, epistatic, and maternal. The additive effects characterize the average effect of a strain constrained to a dosage-like model. Such a simple model is not always sufficient to accurately model data, such as the situation that an F1 hybrid is not approximately the midpoint between the parental strain phenotypes. We account for this potential deviation from additivity with an inbred effect, which is in contrast to the more traditional view of nonadditivity as dominance. This parameterization of the model is appropriate for our pilot data because, considering *J* parental strains, there will be *J*(*J* – 1) possible F1 hybrids, and only *J* inbreds. When *J* is greater than 2, which is likely, the number of possible hybrid F1 will outnumber the *J* lines. Thus modeling the state of being outbred as the default state more intuitively matches the structure of our data.

Epistatic and maternal effects represent other potential sources of deviation from strict additivity. Epistatic effects are essentially an interaction between strains, thus allowing a specific F1 hybrid to deviate from its additive expectations. Maternal effects can capture strain-specific POEs where there is an average difference between reciprocal F1. As demonstrated in Lenarcic *et al*. (2012), consider pilot data that are some subset of the *J* inbred strains and their F1. The strain identities of dam, sire, and dam-sire pair for individual *i* are indexed as *j*[*i*], *k*[*i*], and (*j, k*)[*i*], respectively. We model the pilot data as

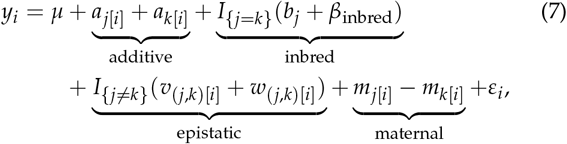

where *y* is the continuous phenotype value, *μ* is the intercept, *a* is a strain-specific additive or dose effect, *β*_inbred_ and *b* are respectively a general inbred effect and a strain-specific inbred effect that are included only if individual *i* is inbred, *v* and *w* are respectively symmetric and asymmetric strain-by-strain interaction effects that we will refer to as epistatic effects and are only included if individual *i* is a hybrid, *m* is a strain-specific maternal effect, and *ε*_*i*_ is the individual-specific noise (deviation from the model expectation) and is distributed: *ε*_*i*_ ~ N(0, *σ*^2^). The model can also include important covariates, such as sex, that need to be adjusted for as fixed effects. The complete set of founder strains and all their reciprocal F1 hybrids represent what is called a diallel, which would allow for the estimation of the full set of strain effects described. Although an incomplete diallel cannot estimate all the strain effects, it still provides information that can be used to estimate ***ϕ***. See Appendix A for descriptions of prior specifications used.

#### Connecting strain-level effects to QTL effects

Transitioning from strain-level genetic effects ***ϕ*** to the effect of a single QTL *θ* requires some strong assumptions. Pilot data consisting solely of F1 individuals cannot provide information about specific loci or the number of loci contributing to a strain effect; there are an infinite number of genetic architectures that can explain a given strain effect. It is possible that conducting a small set of F2 crosses and investigating the variability in phenotype for the resulting population could provide information about the trait genetic architecture, such as distinguishing between highly polygenic and oligogenic traits, but here we focus on using only the F1 generation. We make the assumption that the strain effects represent the effect of a single QTL, which is biologically unlikely but provides an informed approach to connect information in the pilot data to the power calculations. Eq 7 provides expected phenotype values for a given cross of two strains, assuming the trait is controlled by a single QTL. Consider comparing strains *A* and *B*. Eq 7 can be used to estimate E(*y*_*AA*_), E(*y*_*AB*_), and E(*y*_*BB*_). From these predicted values, traditional single QTL additive (*α*) and dominant effects *δ* can be estimated from Eq 4 and Eq 5. These estimates along with estimates of *σ*^2^ can then be used with the power calculation machinery described before. Different QTL effects will be estimated from the model in Eq 7 for different potential crosses of inbred lines.

#### Generalizing single QTL effect for complex traits

The QTL effects defined in Eq 4 and Eq 5 make the assumption of a single underlying QTL; however, from the perspective of the strain-level model in Eq 7 these parameters can be interpreted as strain-level predicted phenotype contrasts. The corresponding variance expressions in **Table S2** then become simplifying functions of contrasts. Generally speaking, all of these quantities can be viewed as point summaries of predicted contrasts from a strain-level model, and thus have a meaningful and potentially useful interpretation outside of the single QTL Mendelian context.

### Decision theoretic approach

Different inbred lines will possess differing segregating variants to potentially identify. We use our model of pilot data to make predictions for some set of possible experiments, which can be viewed in the context of a decision theoretic space (Raiffa and Schlaifer 2000). Define 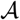 as the set potential experimental crosses. Considering *n* inbred lines, 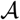 could contain all of or some subset of the (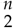) potential F2 crosses and 3*n*(*n* – 1) potential BC.

#### Power as utility function

Let an element of 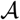, 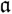 represent a specific action, in this setting, a cross experiment that has corresponding single QTL effect composed of *α* and *δ*. If we define *Q* to be a binary variable that the QTL that causes *θ* (*α* and *δ*) is successfully mapped:

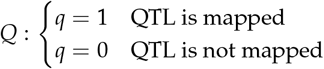

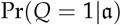 represents the power that the QTL is successfully mapped, and can be calculated using the noncentral *χ*^2^ distribution described previously. We next define 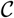 to be the consequence or experiment outcome space for a QTL mapping experiment, where *c* = {*q*_1_,… *q*_*p*_} is the specific joint mapping outcome of the *p* QTL that underly the strain effects. Along with using posterior contrasts as the utility function, this approach allows for the assumption of a single QTL causing the strain-level effects to be reduced.

A utility function is an important concept in decision theory. It provides a common scale to compare potential experimental outcomes, and select optimal experiments. Alternative utility functions can be devised and easily swapped to place value on differing aspects that investigator want to prioritize. We define a utility function, *u*(.), to map from 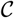 to the reduced utility space, 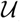, which we pose as a function of power, a natural quantity to prioritize. Consider the probability of a specific consequence, which will be a product of a function of the individual power for each 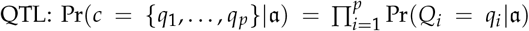. We define *u*(.) to be the count of *p* QTL that were successfully mapped: 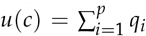. The probability of a utility *v* can be calculated from subsets of 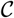:

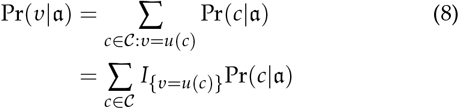

Strictly speaking, the probability of a utility is also dependent on QTL effect 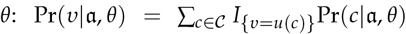. *θ* can be marginalized out through integration: 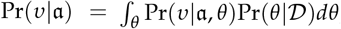, where 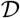 represents the pilot data. The probability of this utility function provides an evaluation of the uncertainty of mapping QTL of a given effect size, but does not take into account the uncertainty of *α, δ*, and *σ*^2^, which are produced from the Bayesian model. Through Gibbs sampling or some other Markov Chain Monte Carlo (MCMC) method, a Bayesian model can produce *S* draws from the posterior distribution of these parameters. Monte Carlo (MC) averaging allows us to take into account this extra source of variability, resulting in the posterior expected utility for cross 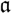:

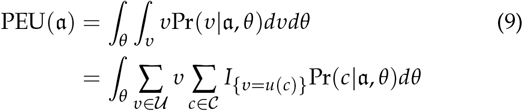

where *θ* is the vector function of *α, δ*, and *σ*^2^. The quantity 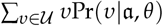 within the 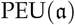 is the expected utility for a single draw s from the Bayesian model. This quantity is then averaged over the QTL effect space of the posterior distribution, traversed through the MC samples. This can be summarized as a point estimate such as the posterior mean or median, or the posterior distribution of expected utilities can be plotted for a given cross 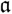. Interpretations of the 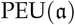 will vary amongst utility functions, but we will focus our discussions on power as the utility being maximized.

If it is assumed that all *p* QTL have the same effect size, the utility function *u*(*c*), the number of *p* QTL that were successfully mapped, follows a binomial distribution. Consider simple case of a single QTL (*p* = 1), in which the binomial reduces to the Bernoulli distribution. In this setting, the 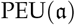 reduces to the posterior probability of mapping the QTL. When *p* is greater than one, as with a binomial variable, 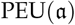 now represents the expected number of QTL to be mapped. Our approach should be flexible to any reasonable utility function investigators can define, but we emphasize power because its 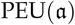 are easy to interpret, as well as posterior contrasts, which have require less assumptions and better extend to highly complex phenotypes.

### Simulation of diallel data

Diallel data were simulated from Eq 7 based on a number of different strain-level effect settings to assess how well DIDACT performs in a variety of genetic architectures according to various quantities, such as the estimation of the strain-level effects, and more importantly, utility. Specifically 100 realizations were simulated of a full and balanced diallel of eight strains with five individuals per diallel cell, resulting in a total of 320 observations, for each specification of strain-level effects. The strain-level effects were scaled so that the proportion of total variance was controlled such that:

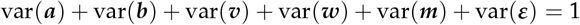

The proportion of variance from strain-level effects was set to 50 evenly spaced values between 0.01 to 0.99. The following strain-level effect specifications were used:

1. **biallelic_add:** The inbred, maternal, and both epistatic effects are all set to 0. Two alleles are simulated for the additive effects and distributed evenly amongst the parental strains.
2. **rnorm_add:** The inbred, maternal, and both epistatic effects are all set to 0. The eight additive alleles are simulated such that *a*_*j*_ ~ N(0,1).
3. **complex:** The additive, inbred, maternal, and both epistatic effects are all simulated such that *a*_*j*_, *b*_*j*_, *m*_*j*_, *v*_*jk*_, *w*_*jk*_ ~ N(0,1). The proportional of total variance is evenly split between the five strain-level effect types.

Table 1 describes these three diallel data-generating models, including the type and number of non-zero effects.

### Availability of data and software

All analyses were conducted in the statistical programming language R (**?**). The R package DIDACT (Diallel Informed Decision theoretic Approach for Crosses Tool), available on GitHub at https://github.com/gkeele/DIDACT, can estimate strain-level effects from diallel data using a Bayesian hierarchical model, and then perform the posterior utility analysis. The R package BayesDiallel can alternatively be used to estimate the strain-level effects, and potentially used as inputs to DIDACT.

DIDACT includes three diallel data sets, each with a number of phenotypes, of the CC founders (Churchill *et al*. 2004; Collaborative Cross Consortium 2012; Srivastava *et al.* 2017), described in great detail in Lenarcic *et al.* (2012). Specifically, results shown here are from a hemoglobin trait measured in 626 mice. The CC founders represent the following inbred strains of mouse (abbreviated names in parentheses): A/J (AJ), C57BL/6J (B6), 129S1/SvImJ (129), NOD/ShiLtJ (NOD), NZO/H1LtJ (NZO), CAST/EiJ (CAST), PWK/PhJ (PWK), and WSB/EiJ (WSB).

An additional diallel data set of the CC founders of response to Influenza A virus (IAV) infection phenotypes is used here and included in the DIDACT package, with the original data available at https://github.com/mauriziopaul/flu-diallel. In previous work (Maurizio *et al*. 2018), we investigated strain-level effects in day four post-infection (D4 p.i.) body weight loss percentage in a diallel of the CC founders. The phenotype of interest is a response to infection, in which three infected animals were compared to a single mock-infected animals. Occasionally three infected animals were not observed at later time points, which was accounted for through a multiple imputation procedure that imputed unobserved animals from the posterior predictive distributions of the BayesDiallel model (Lenarcic *et al*. 2012). Here only a singly imputed data set of 131 outcomes is used, as this example is a proof of principle for DIDACT, and not a rigorous investigation of strain-level effects.

## Results

We provide results from simulations and example analyses from diallel data of the CC founders to demonstrate the decision theoretic procedure used in the DIDACT approach. The use of QTL mapping power as a utility function depends on assumptions about the effect of a single putative QTL in a bi-parental cross (described in **Table S1**) given strain-level effects estimated from diallel data based on the parameterization described in Eq 7. This assumption is most straightforward in the case of a largely Mendelian phenotype, in which a single locus modulates the variation observed in a relatively deterministic manner, and as such, the QTL effect *θ* can draw from the strain-level effect ***ϕ*** wholly. When the genetic architecture of the phenotype is complex, as is likely with many traits, the use of predicted phenotype contrasts requires less assumptions, as the nominal interpretation of power is no longer valid.

### Simulations

100 simulations per 50 levels of strain-level effect sizes (evenly spaced between 0.01 and 0.99) for three different model types were used to evaluate strain-level effect estimation accuracy, and more importantly, DIDACT’s utility estimation. For effect estimation accuracy, the averaged mean squared error (MSE) of the additive strain-level effects (***a***) was used, calculated as:

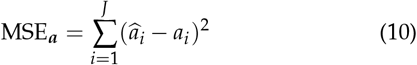

where *J* is the number of founders (8 in the CC) and 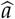 is the posterior mean of *a*. For utility estimation accuracy, the rank correlation of posterior contrasts was used.

DIDACT can effectively estimate strain-level effects, but more importantly, is consistently accurate at ranking posterior utilities (Figure 3). The simulations show that DIDACT can both accurately estimate the additive strain-level effects when those are the only non-zero effects in the data (Figures 3 A and B). When many non-zero and non-additive strain-level effects are present, the accuracy of the effect estimation is reduced (Figure 3 C). Despite this reduction in effect estimation accuracy for the BayesDiallel model in a complex setting, DIDACT still estimates and ranks utilities effectively (Figure 3 F) in comparison to the simpler additive settings (Figures 3 D and E).

**Figure 3.**
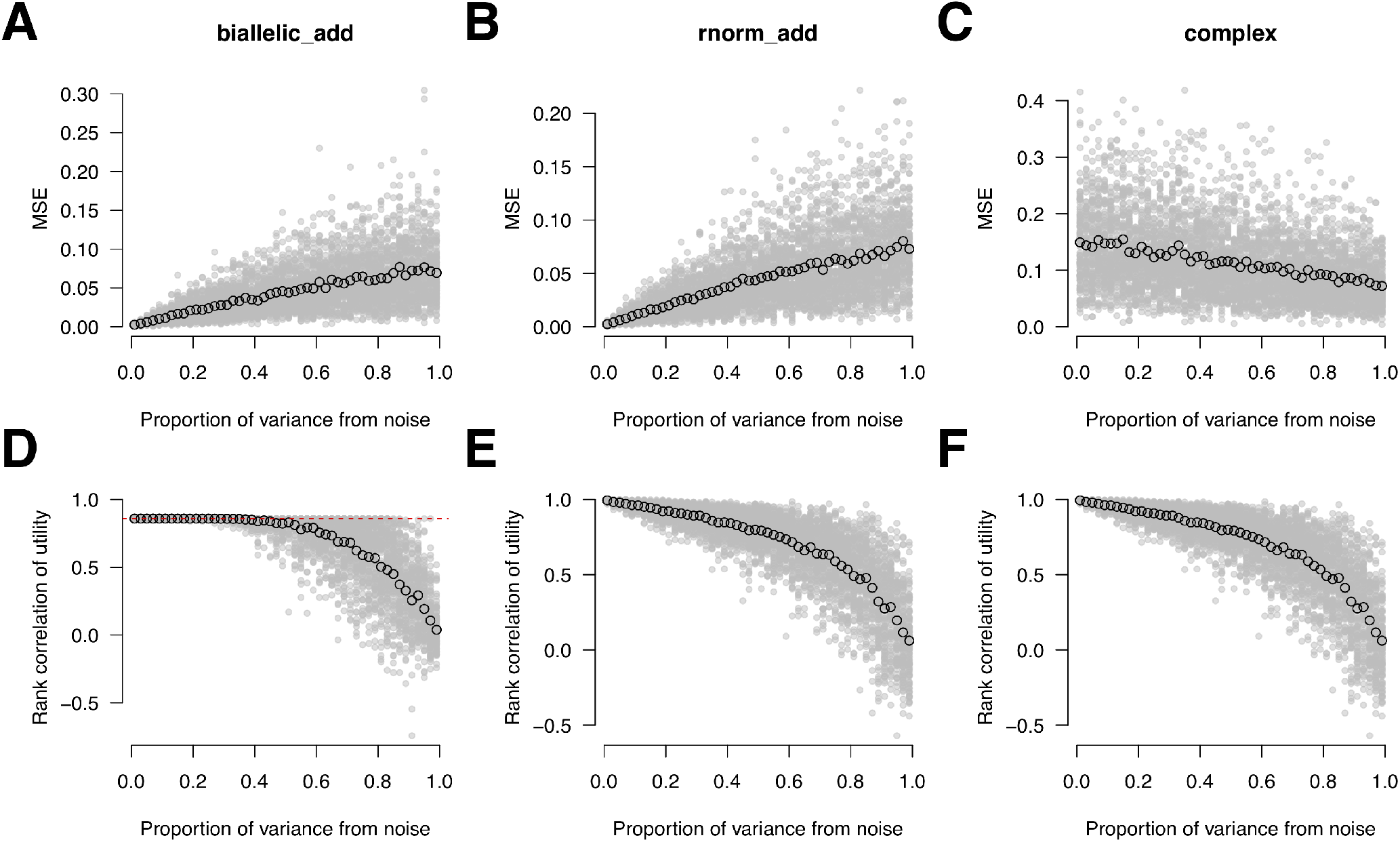
The BayesDiallel model’s estimates of the additive strain-level effects improve with decreasing proportion noise variance (A-B). When the underlying strain-level effects are complex, the BayesDiallel model struggles to accurately estimate the additive effects, even with decreasing variation due to noise (C). DIDACT’s ranking of posterior contrasts for simulated diallel crosses of varying complexity improves as proportion noise variation decreases (D-F), suggesting that even though effect estimation is challenging in the presence of complex strain effects, DIDACT’s posterior utility is robust to this complexity. 100 diallel crosses were simulated per level of variance explained (50 values evenly spaced from 0 to 1) per model type. **biallelic_add** is a Mendelian QTL with two alleles distributed evenly amongst the founder strains. **rnorm_add** is also an additive-only model but with *a* ~ N(0, *τ*^2^), where *τ*^2^ is one minus the proportion of variance determined by noise. **complex** has a complex set of non-zero strain-level effects for additive, inbred, maternal, and both symmetric and asymmetric epistatis, each contributing equally to the overall variance. Gray dots represent single simulations, and black circles the mean over the 100 simulations. The red dashed line in D marks the max rank correlation (< 1) for **biallelic_add** due to the true utilities at a Mendelian locus containing ties whereas the estimated ones will be continuous.

### Mendelian phenotype

To demonstrate a straightforward application of DIDACT to a phenotype largely driven by a single locus, we use resistance to IAV infection and the *Mx1* gene. As described in greater detail in the **Methods**, the phenotype of interest is day four postinfection (D4 p.i.) body weight loss percentage in a diallel of the CC founders.

#### Mx1 is a critical host-resistance factor in mice

It has previously been shown that *Mx1* largely drives IAV-resistance in the CC founders, and has three major functional classes corresponding to the three subspecies of *Mus musculus: domesticus* (hereafter *dom*; CC founders with *dom* allele are AJ, B6,129, NOD, and WSB), *castaneus* (*cast*; CAST), and *musculus* (*mus*; PWK and NZO) (Ferris *et al*. 2013). The *dom* allele of *Mx1* was found to be functionally null and those individuals susceptible to IAV infection, whereas *mus* and *cast* confer degrees of resistance.

Though IAV-resistance is largely driven by *Mx1*, the genetic variation at the gene in the diallel of CC founders is more complicated than a bi-allelic locus. Instead, *Mx1* has multiple functional alleles, *mus* and *cast* contrasting the null allele *dom. mus* has a dominant mode of action, conferring approximately the same level of resistance to IAV in *dom/mus* individuals as in *mus/mus*, whereas *cast* is additive with *cast/dom* being intermediate between *dom/dom* (low resistance) and either *mus* carriers or *cast/cast* (highest resistance).

The increased IAV-resistance of *mus* and *cast* is apparent in the raw data as darker horizontal and vertical bands for NZO and PWK, and to a lesser extent for CAST (Figure 4A). The raw data also suggests that WSB may posses additional modifiers of resistance with some hybrids displaying high resistance for individuals with homozygous *dom*. The resistance conferred by variation at *Mx1* is further confirmed in the strain-level effects estimated through BayesDiallel, highlighted in Figure 4B.

**Figure 4.**
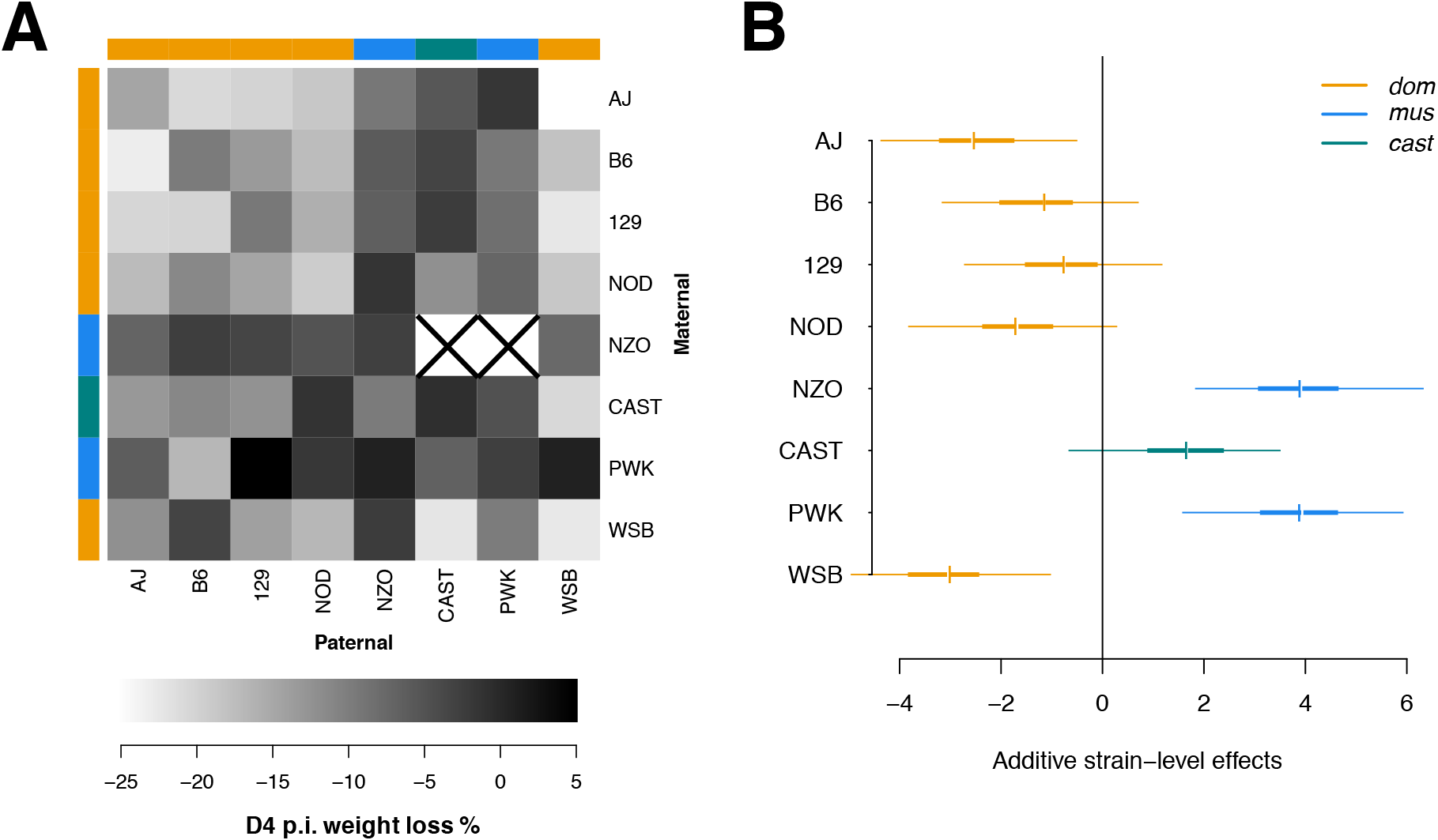
*Mx1* is as a driver of IAV-resistance. This is manifested in the raw data of D4 p.i. body weight loss % in a diallel of the CC founders and their hybrids (381 mice) as relatively dark vertical and horizontal bands at the NZO, CAST, and PWK founders, representing low percent weight loss. (A). Squares with a black “X” represent crosses that produce no viable offspring (NZO_♀_ × PWK_♂_ and NZO_♀_ × CAST_♂_). The subspecies allele at *Mx1* for each CC founder strain is indicated by colored bars below and to the right of the phenotype grid, with orange representing *dom*, blue *mus*, and green *cast*. The *mus* and *cast* alleles are functional, whereas the *dom* is functionally null. Moreover, the *mus* allele is dominant in comparison to *cast*. The strain-level additive effects estimated from the BayesDiallel model correspond to the subspecies lineages at the *Mx1* locus, with *mus* (NZO and PWK) conferring more resistance than *cast* (CAST) (B). The highest posterior density (HPD) intervals from the BayesDiallel strain-level additive effects are shown with 95% HPD as thin lines and 50% HPD as thick lines. The posterior means and medians are indicated as colored and white ticks, respectively. The effects closely match those estimated in Maurizio *et al*. (2018), which used a more complex BayesDiallel model, and summarized over multiply imputed data sets.

#### DIDACT favors crosses with segregating Mx1 variants

For a Mendelian phenotype, DIDACT should prefer crosses that maintain segregating alleles at the locus, in this case *Mx1*. In particular, it should prioritize crosses that match *dom* with *mus* or *cast*. Crosses with no segregating functional variants of *Mx1* will have greatly reduced phenotypic variation, and ultimately not be able to detect the largely Mendelian locus.

As expected, DIDACT largely estimates high QTL mapping power for crosses that possess multiple segregating *Mx1* alleles, either *dom* with *mus* or *cast*. Mean posterior power from DIDACT for F2 experiments are shown in Figure 5A and for BC experiments in Figure 5B. For F2, PWK × WSB and NZO × WSB are ranked 1^st^ and 2^nd^ in posterior power, respectively. CAST × PWK is ranked 3^rd^, which matches the expectations that *mus* × *dom* are preferable to *cast* × *dom*. WSB is likely preferred as the *dom* carrier because of its relatively unique phenotype. In general, other crosses that maintain multiple segregating alleles of *Mx1* are preferred in comparison to crosses that fix the variability at the locus.

**Figure 5.**
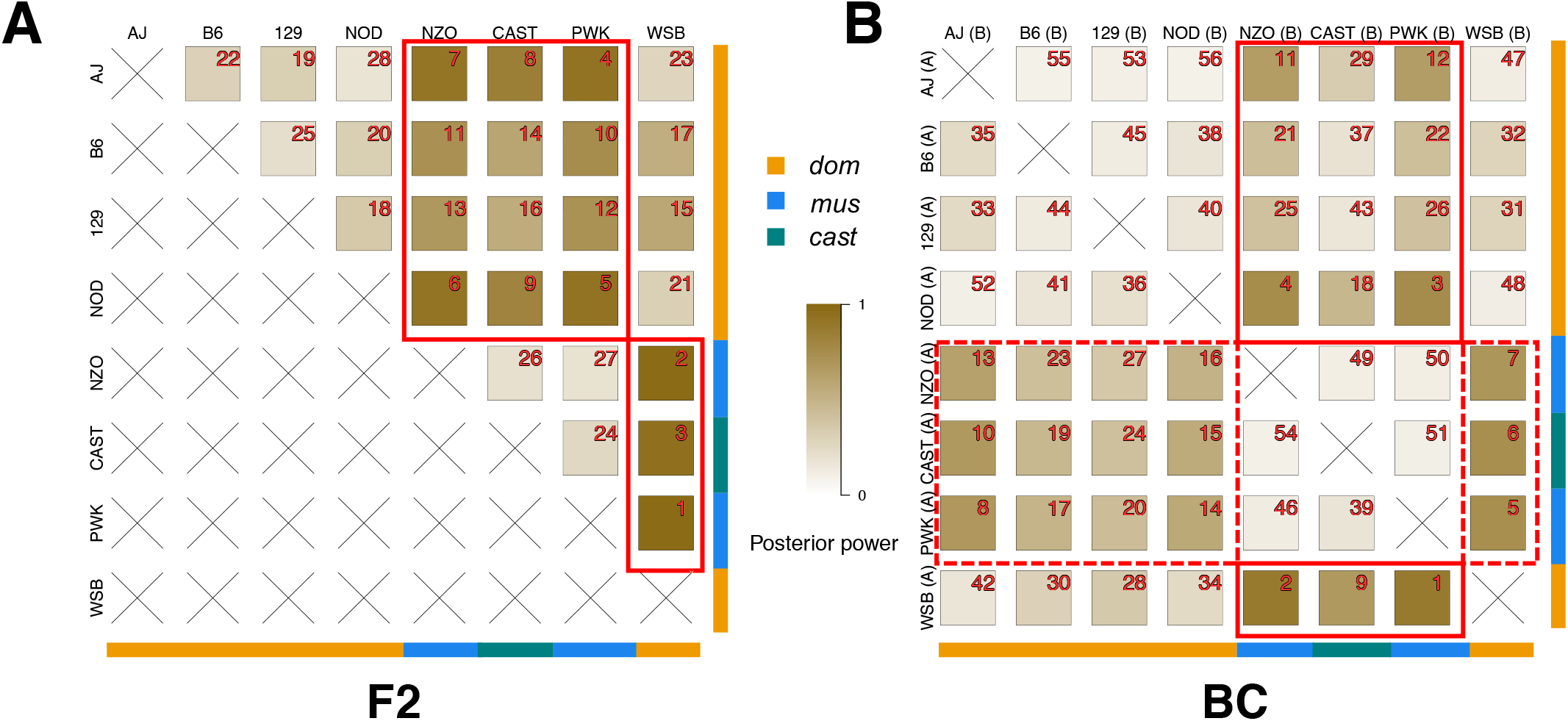
For IAV resistance measured as D4 p.i. weight loss %, DIDACT ranks F2 and BC experiments that maintain genetic variability at *Mx1* higher. Posterior mean utility as power to map a single QTL in 60 mice, for the 28 possible F2 crosses (A) and the 56 possible BC (B) of the CC founder strains. The red number in the top right corner of a cross square represents the posterior power rank for a given cross type (F2 or BC). Subspecies allele at *Mx1* is indicated by colored bars below and to the right of the grids, with orange representing *dom*, blue *mus*, and green *cast*. For F2, the solid red squares enclose crosses that pair functional alleles (*mus* and *cast*) of *Mx1* with *dom*. For BC, the solid red squares enclose crosses that pair strains with functional alleles with strains with *dom* and the backcrossed parent has *dom*. The red dashed squares also pair functional with nonfunctional alleles of *Mx1*, but the backcrossed parent has the functional allele instead. Backcrossing strains with *mus* are expected to have reduced power due to the dominance of the allele.

For BC, similar patterns are observed. Though subtle, DIDACT generally favors BC for crosses of *dom* and *mus* in which the backcrossed parent strain (A) has *dom*, thus reflecting the dominance of *mus*. In the case of a completely dominant QTL, there would be no power to map the QTL in a cross of A × B, given that A is the backcrossed parent and possesses the dominant allele. For example with *Mx1*, WSB (A) × PWK (B) is ranked 1^st^ whereas PWK (A) × WSB (B) is 5^th^. Though not perfect, reflecting the complexities of real data and a phenotype with potentially additional genetic modifiers, DIDACT largely favors crosses of *dom* (A) × *mus* (B), given a specific pairing of strains.

As with the F2, DIDACT prefers crosses of WSB (*dom*) with strains that possess *mus* or *cast*, likely reflecting strain-level WSB effects that are independent of *Mx1* (Maurizio *et al*. 2018). It is also important to reiterate that this analysis of *Mx1* represents a single imputation of multiply imputed data, representing an additional source of noise.

### Complex trait: calculated hemoglobin

Many phenotypes are the not Mendelian, but rather the product of potentially many loci across the genome and complicated genetic architectures. Such traits violate the assumption of attributing the strain-level effects to a single QTL. DIDACT can still be used to analyzed these complex traits. Though the power utility function is still meaningful in selecting crosses of phenotypically distinct strains, the single QTL assumption can be relaxed by using the phenotypic contrasts directly as the utility function. Here, contrasts are used for calculated hemoglobin (cHGB), which is likely not Mendelian.

As reported in Lenarcic *et al*. (2012), blood phenotypes were measured on 626 mice from a diallel of the CC founders, which included cHGB (g/dl), an estimate of the quantity of hemoglobin in the blood (Figure 6A). The raw data do not suggest obvious additive strain-level effects as seen with D4 p.i. weight loss % (Figure 6A), though great phenotypic variability is observed, and patterns that reflect complex strain-level effects. CAST individuals have a notably lower trait-value. Additionally, PWK hybrids with AJ, B6, and 129 have unusually high trait-values.

**Figure 6.**
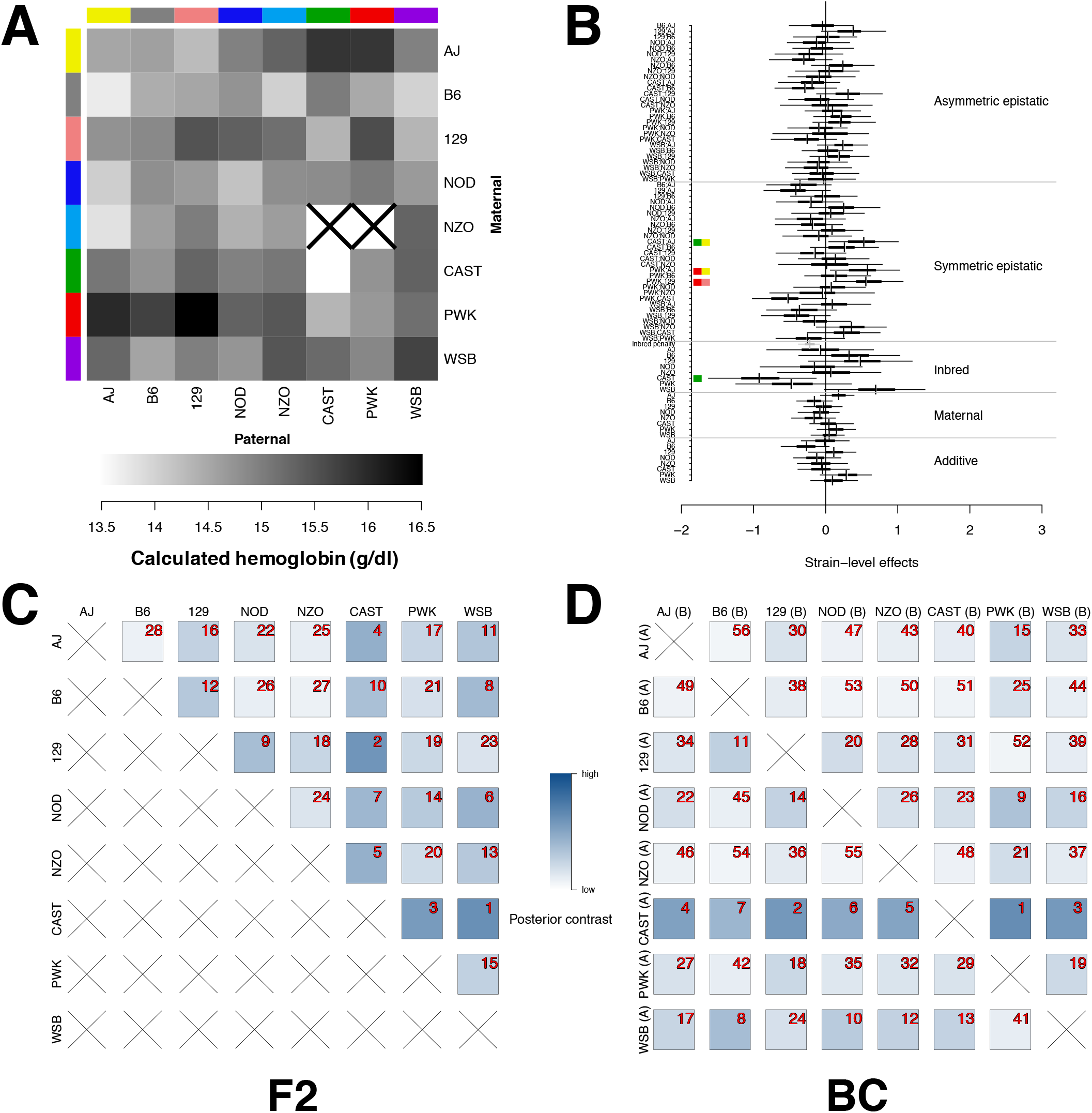
DIDACT can analyze complext traits, such as calculated hemoglobin (cHGB) in a diallel of the CC founder strains of 626 mice. The mean raw data for cHGB suggests inbred effect for CAST (A). A black “X” represents crosses that did not produce viable offspring. BayeDiallel estimates a number of non-zero strain-level effects (B). Effects are represented as HPD intervals, with 95% and 50% as thick and thin lines, respectively, and colored and white ticks representing posterior means and medians, respectively. The gray bar represents the the overall inbred parameter. A Green square along the y-axis highlights a non-zero inbred CAST effect. Non-zero symmetric epistatic effects are similarly emphasized for PWK × AJ, PWK × 129, and CAST × AJ. DIDACT posterior contrasts for F2 (C) and BC (D) experiments rank crosses with CAST highly.

These patterns are reflected in the BayesDiallel estimates of non-zero strain-level effects (Figure 6B). CAST has a nonzero negative inbred effect. For symmetric epistatis, PWK has positive non-zero effects with AJ and 129, and a suggestive positive effect with B6. CAST also has a non-zero positive effect with AJ, which is also consistent with the raw data. Stable effects are based on observations across multiple crosses and replicates of each pairing, thus likely representing heritable factors.

The posterior contrasts for possible F2 and BC mapping crosses are shown in Figures 6C and 6D, respectively. DIDACT’s ranking of contrasts largely prefers crosses with CAST, corresponding to its non-zero strain-level effects. Notably For BC, the strongly negative CAST inbred effect is manifested in DIDACT estimating higher posterior power for BC in which CAST is the backcrossed parent (A). If only CAST hybrids are observed, the mapping cross will not produce as much phenotypic variation, which stems from the CAST homozygotes in this trait. DIDACT can dynamically take into account complex strain-level effects, and rank crosses accordingly.

### Additional DIDACT summaries

DIDACT can provide more detailed descriptions of the predicted bi-parental crosses than shown in Figures 5 and 6. At its core, DIDACT is an extension of the BayesDiallel model, in which the strain-level effects from the Bayesian hierarchical model are propagated to predetermined utility functions, and as such, posterior intervals can be produced in addition to the point estimates for any function of the model parameters.

Three potential F2 crosses were selected from the full panel from cHGB (Figure 6C), and are presented in **Figure S2**. Additional summaries include a histogram of the posterior distribution of the utility function, in this case contrasts, the median utility represented as a vertical dashed line on the histogram, predicted phenotypes for each QTL genotype (based on a single QTL assumption) as bar plots, and the phenotypic variation attributable to the strain-level effects as a pie chart, all overlayed on the square with posterior mean determining the background color. CAST × WSB has higher mean posterior contrast compared to NZO × WSB and AJ × B6. This is reflected in phenotype predictions that vary more greatly (bar plots) and an estimated higher proportion of phenotypic variance explained by the strain-level effects (63%).

### Parent-of-origin effects and RBC^PO^

There is not currently a satisfactory approach and solution for parameterizing QTL effects that contain a POE mode of action, such as exists for additivity and dominance as described in Eq 4 and 5 as well as in **Tables S1** and **S2**, which ultimately limits the ability of DIDACT to make power calculations for RBC^PO^ as described in Figure 1C. However, it is possible for DIDACT to characterize the utility in terms of predicted BC, but with the maternal and paternal identities fixed as in the RBC^PO^. Though the power calculation will not correspond to the design specified in Figure 1C, in which three genetic states are observed in comparison to two for BC, differences in QTL mapping power for BC that are equivalent except for the maternal and paternal statuses of the backcrossed parental strain and F1 are potentially interesting, shown in **Figure S3** for cHGB.

RBC^PO^ that have markedly different posterior contrasts can identify pairings of strains with interesting combinations strain-level maternal effects in Figure 6B. The BayesDiallel model estimates several suggestive strain-level maternal effects, primarily AJ and B6, suggesting that different RBC^PO^ may have varying posterior contrasts. For example B6 (A_♀_) × PWK (B_♂_) is ranked 50^th^ compared to 13^th^ for B6 (A_♂_) × PWK (B_♀_) (**Figure S3**). More work is needed to formally extend analytical expressions of QTL mapping power to RBC^PO^, but posterior utility can still be used to highlight interesting crosses based on the diallel data.

## Discussion

DIDACT represents an approach to selecting experiments using the information contained within diallel crosses. In this approach, the diallel cross is used as pilot data to characterize strain-level genetic effects within a Bayesian hierarchical model, which are then applied to utility functions for the purposes of identifying promising bi-parental crosses for mapping QTL. Herein, we define utility to be QTL mapping power, though other functions could be used, such as phenotype contrasts, so long as the strain-level effects are their inputs.

### Violation of assumptions connecting strain-level effect to QTL effect

DIDACT requires assumptions to connect the strain-level diallel effects to experimental design-relevant utility functions. How easily these spaces are connected will vary with the complexity of the phenotype. DIDACT performs well in a mostly Mendelian phenotype in which the strain-level effects can be correctly attributed to a single putative QTL. However, phenotypes that are highly heritable can be modulated through many loci, often with few to none exhibiting large effects, height in humans being a clear example (Wood *et al*. 2014). In the vast majority of complex traits, the assumption of a single QTL absorbing all or most of the strain-level effects is wildly optimistic. However, we posit that though the assumption is unlikely, its use as a utility function can still produce a useful analysis of potential bi-parental crosses.

The power utility function used in DIDACT favors QTL that explain a large proportion of the variability in the phenotype. In fact, the power function corresponds closely to the variability explained by the putative QTL, which will relate to the variability explained by strain identity in the diallel in the context of complex but heritable phenotypes. Alternatively, the phenotype contrasts could be used as the utility function instead. Though the interpretation of the posterior utility as an accurate power is unrealistic, it will still select pairings that are phenotypically distinct, which is a common criterion for selecting crosses. And, it will do so in a highly principled approach that intuitively accounts for uncertainty.

### Genetic similarity of strains

DIDACT, in its current form, does not make use of any information regarding the similarity of the parental strains included in the diallel cross, which could further inform how appealing an experimental cross is for fine-mapping detected QTL. The reduced complexity cross (RCC) is a developing approach in systems genetics (Williams and Williams 2017) in which strains that are phenotypically divergent but genetically similar are crossed, such as C57BL/6J and C57BL/6N sub-strains (Khisti *et al*. 2006; Mulligan *et al*. 2008; Kumar *et al*. 2013; Simon *et al*. 2013; Kirkpatrick and Bryant 2014). RCC provide a powerful tool for fine-mapping causal variants because the genetic variability between strains is greatly reduced.

There are a number of ways that DIDACT could be modified to incorporate genetic similarity information, probably most simply through the utility function. The utility function could be expanded to flexibly weight potential experimental crosses by the genetic similarity, resulting in posterior utilities that are informed by both phenotype and genetic similarity. We believe this highlights the potential of DIDACT, and its underlying concept in general, to be flexible to the context of the experimental system, at the hierarchical model, but particularly at the utility function.

### Extension to multiparental populations

The DIDACT analyses presented here are from diallel crosses of the CC founders, which naturally poses the question of designing experiments of the CC strains based on their related diallel data. The CC population represents a multiparental population (MPP), in which each individual is descended from all of the founder strains. Extending DIDACT to experiments of an MPP RI panel is challenging because the analytical power calculations (Sen *et al*. 2005, 2007) are based on bi-parental populations with two founder alleles. Though these could be generalized to populations with multiple alleles, the CC strains also differ from traditional mapping crosses in that the recombination events that randomize segments of the genome to allow for QTL mapping have already occurred in the breeding scheme that derived the inbred strains.

Maybe a more natural approach to extending DIDACT to an MPP RI panel would be to consider the MPP RI panel as a large sparse diallel, with off-diagonal cells representing the F1 hybrids, in the case of the CC, these would be CC-RIX (Bogue *et al*. 2015). DIDACT could then be adapted to select potentially interesting but unobserved CC-RIX based on the CC-specific strain-level effects. Effectively adapting DIDACT for design of MPP experiments is an area of interest for future research.

### Summary

DIDACT is a novel approach to using prior collected diallel data from a panel of inbred strains to inform the selection of potential downstream experiments according to a user-specified utility function, in our case, power to map QTL in bi-parental cross experiments, consisting of F2, BC, and RBC^PO^. The core of this approach is to propagate the uncertainty present in the Bayesian hierarchical model through to the utility functions, which can be customized to the needs and constraints of the system at hand.

As proof of principle, we evaluated DIDACT in a phenotype known to be Mendelian: resistance to IAV-infection, which is largely modulated by the *Mx1* gene with a null (susceptible) and two non-null (resistant) alleles. DIDACT largely evaluated bi-parental crosses of null with non-null *Mx1* strains as having higher posterior power to map the QTL. For the non-Mendelian calculated hemoglobin, DIDACT favors crosses that pair strains with contrasting phenotypes. Though the posterior power as utility, in the sense of its nominal interpretation as power, is highly optimistic, still provides a reasonable metric for comparing potential experiments, given the available pilot data. This approach has many potential applications, in terms of both the utility functions that are being evaluated and the model organism systems, many which have sparse diallel data available in the form of strain surveys. DIDACT represents a philosophical advancement in terms of good experimental design and efficient use of available resources.

## Supporting information

## Acknowledgments

We thank Amy Herring, now at Duke University, for her thoughts and suggestions on this paper, which began as a group project in her Bayesian statistics course at UNC. We also thank Alan Lenarcic for his expertise on Bayesian modeling of diallel crosses.

This work was primarily supported by the National Institute of General Medical Sciences under awards R01-GM104125 and R35-GM127000 (to W.V.).

## Appendix A Prior specification

Following the lead of (Lenarcic et al. 2012), conjugate priors were used for the parameters in the BayesDiallel model. For example, the strain-level additive effects are distributed following 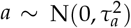. For fixed effect terms, such as *β*_inbred_, *τ*^2^ is set to 10^3^. For the variance parameters, consider *σ*^2^ which is distributed following *σ*^2^ ~ IG(*v*/2, *ψ*/2). For the hyper parameters *v* and *ψ* 0.02 and 2 were used, respectively. These represent diffuse priors, with the intention of allowing the information in the data to inform the estimates. The hyper parameter values can be adjusted within DIDACT R package.

