## Supplementary material for "Bayesian decision theoretic design of two-founder experimental crosses given diallel data"

```
##############################################################################
##############################################################################
########
########  Title: File S1
########  Purpose: Generate figures for DIDACT manuscript
########  Authors: Greg Keele
########
##############################################################################
##############################################################################

##############################################################################
########  
########  Necessary Packages
########
##############################################################################

library(DIDACT) 
## - Purpose: Rank potential bi-parental crosses using diallel pilot data
## - Available at https://github.com/gkeele/DIDACT
## - If you use the R package devtools, simply use:
##   - install_github("gkeele/DIDACT")

##############################################################################
########  
########  Figure 3: Simulations results from BayesDiallel and DIDACT
########
##############################################################################
## See sim_biallelic_add.R: Figures 3A,D
## See sim_rnorm_add.R: Figures 3B, E
## See sim_complex.R: Figures 3C, F

##############################################################################
########  
########  Figure 4: IAV-induced weight loss diallel data and additive effects
########
##############################################################################
## Subfigure A: Mean Day 4 pi weight loss
diallel.phenotype.map(data = flu.dat, 
                      mother.str.var = "dam", 
                      father.str.var = "sire", 
                      phenotype = "pct_D4", 
                      phenotype.title = "Day 4 post-infection weight loss", 
                      do.reorder = FALSE,
                      strains.reorder = NULL, 
                      strain.names = c("AJ", "B6", "129", "NOD", "NZO", "CAST", "PWK", "WSB"),
                      strain.colors = rev(c("orange2", "dodgerblue2", "#008080", "dodgerblue2", 
                                          rep("orange2", 4))), 
                      strain.cex = 1.5)
# Phenotype scale
diallel.phenotype.scale(data = flu.dat, 
                        mother.str.var = "dam", 
                        father.str.var = "sire", 
                        phenotype = "pct_D4")

## Subfigure B: Additive effects
# Run Gibbs sampler to estimate strain-level effects
set.seed(10)
flu.par <- diallel.gibbs(phenotype = flu.dat$pct_D4, 
                         sex = as.numeric(flu.dat$Sex=="F"),
                         mother.str = flu.dat$dam, 
                         father.str = flu.dat$sire, 
                         n.iter = 1000, 
                         burn.in = 10000,
                         use.constraint = TRUE,
                         strains.reorder = LETTERS[1:8])
caterpillar.plot(gibbs.object = flu.par, 
                 include.effect.types = "add", 
                 override.col = c("orange2", "dodgerblue2", "#008080", "dodgerblue2", rep("orange2", 4)), 
                 override.title = "D4 p.i. weight loss %", 
                 override.xlab = "Additive strain-level effects",
                 include.grid = FALSE, 
                 include.prefix = FALSE,
                 zero.lty = 1,
                 zero.color = "black",
                 effect.label.cex = 1)
legend("topright", 
       legend = c(expression(italic("dom")),
                  expression(italic("mus")), 
                  expression(italic("cast"))), 
       col = c("orange2", "dodgerblue2", "#008080"), 
       lwd = rep(2, 3),
       lty = rep(1, 3), 
       bty = "n")

##############################################################################
########  
########  Figure 5: DIDACT F2 and BC for Day 4 pi weight loss
########
##############################################################################
flu.par.exp <- evaluate.experiments(flu.par, 
                                    n = 60, 
                                    qtl.num = 1)
## Subfigure A: F2
# Grid
diallelPlotter(results = flu.par.exp, 
               cross.type = "f2", 
               pheno.name = "Day 4 p.i. weight loss %", 
               strains.relabel = c("AJ", "B6", "129", "NOD", "NZO", "CAST", "PWK", "WSB"),
               include.info.plot = FALSE, 
               include.off.x = TRUE,
               include.density = FALSE, 
               include.bar.plots = FALSE, 
               include.biparent.labels = FALSE,
               include.rank = TRUE, 
               border.col = "black",
               rank.col = "firebrick1", 
               median.line.col = "black", 
               col.range = c("white", "goldenrod4"),
               label.cex = 2,
               label.padj = -0.5)
## Utility gradient
make.gradient.scale(mode = "vertical", 
                    spectrum = make.spectrum(c("white", "goldenrod4"), n = 100))
## Subfigure B: BC
diallelPlotter(results = flu.par.exp, 
               cross.type = "bc", 
               pheno.name = "Day 4 p.i. weight loss %", 
               strains.relabel = c("AJ", "B6", "129", "NOD", "NZO", "CAST", "PWK", "WSB"), 
               include.info.plot = FALSE, 
               include.off.x = TRUE,
               include.bar.plots = FALSE, 
               include.density = FALSE, 
               include.rank = TRUE, 
               border.col = "black", 
               median.line.col = "black", 
               rank.col = "firebrick1", 
               col.range = c("white", "goldenrod4"),
               label.cex = 2,
               label.padj = -0.5)

##############################################################################
########  
########  Figure 6: cHGB data, effects, and DIDACT F2 and BC
########
##############################################################################
## Subfigure A: cHGB phenotype
diallel.phenotype.map(data = advia.dat, 
                      mother.str.var = "mother.strain.name", 
                      father.str.var = "father.strain.name", 
                      phenotype = "cHGB", 
                      phenotype.title = "calculated hemoglobin", 
                      strain.names = c("AJ", "B6", "129", "NOD", "NZO", "CAST", "PWK", "WSB"),
                      strain.cex = 1.5)
## Gradient scale
diallel.phenotype.scale(data = advia.dat, 
                        mother.str.var = "mother.strain.name", 
                        father.str.var = "father.strain.name", 
                        phenotype = "cHGB",
                        include.decimal = TRUE)

# Subfigure B: cHGB strain-level effects
set.seed(10)
chgb.par <- diallel.gibbs(phenotype = advia.dat$cHGB, 
                          sex = as.numeric(advia.dat$is.female=="F"),
                          mother.str = advia.dat[,1], 
                          father.str = advia.dat[,2], 
                          n.iter = 1000, 
                          burn.in = 10000,
                          use.constraint = TRUE)
caterpillar.plot(chgb.par, 
                 include.effect.types = c("add", "mat", "inbred", "epi_sym", "epi_asym"),
                 override.title = "cHGB", 
                 override.xlab = "Strain-level effects",
                 override.col = c(rep("black", 8), 
                                  rep("black", 8),
                                  rep("black", 8),
                                  "gray",
                                  rep("black", 28),
                                  rep("black", 28)),
                 include.grid = FALSE,
                 manual.limits = c(-2, 3),
                 include.prefix = FALSE,
                 zero.lty = 1,
                 zero.color = "black")
abline(h = 8.5, col = "black", lty = 1)
abline(h = 16.5, col = "black", lty = 1)
abline(h = 25.5, col = "black", lty = 1)
abline(h = 53.5, col = "black", lty = 1)
text(labels = "Additive", x = 1.5, y = 4.5, adj = 0, cex = 0.9)
text(labels = "Maternal", x = 1.5, y = 12.5, adj = 0, cex = 0.9)
text(labels = "Inbred", x = 1.5, y = 20.5, adj = 0, cex = 0.9)
text(labels = "Symmetric epistatic", x = 1.5, y = 38.5, adj = 0, cex = 0.9)
text(labels = "Asymmetric epistatic", x = 1.5, y = 67.5, adj = 0, cex = 0.9)
## Subfigure C: cHGB DIDACT F2
# Run DIDACT for cHGB
chgb.par.exp <- evaluate.experiments(chgb.par, 
                                     n = 50, 
                                     qtl.num = 1)
diallelPlotter(results = chgb.par.exp, 
               cross.type = "f2", 
               pheno.name = "cHGB", 
               col.range = c("white", "dodgerblue4"), 
               utility.type = "contrasts",
               strains.relabel = NULL, 
               include.bar.plots = FALSE, 
               include.density = FALSE, 
               include.rank = TRUE, 
               include.off.x = FALSE, 
               include.info.plot = FALSE,
               include.biparent.labels = FALSE,
               label.cex = 2,
               label.padj = -0.5)

## Subfigure D: cHGB DIDACT BC
diallelPlotter(results = chgb.par.exp, 
               cross.type = "bc", 
               pheno.name = "cHGB", 
               col.range = c("white", "dodgerblue4"), 
               utility.type = "contrasts",
               include.bar.plots = FALSE, 
               include.density = FALSE, 
               include.rank = TRUE, 
               include.off.x = FALSE, 
               include.info.plot = FALSE,
               label.cex = 2,
               label.padj = -0.5)
## Utility gradient
make.gradient.scale(spectrum = make.spectrum(c("white", "dodgerblue4"), n = 100),
                    mode = "vertical", 
                    utility.type = "contrasts")

##############################################################################
########  
########  Figure S3: DIDACT cHGB detailed
########
##############################################################################
make.single.cross.plot(cross = "AJxB6",
                       cross.type = "f2",
                       utility.type = "contrasts",
                       didact.object = chgb.par.exp, 
                       col.range = c("white", "dodgerblue4"), 
                       border.col = "black", 
                       include.var.pie = TRUE,
                       median.line.col = "red", 
                       density.col = "white")
title("AJ (A) x B6 (B)")

make.single.cross.plot(cross = "NZOxWSB",
                       cross.type = "f2",
                       utility.type = "contrasts",
                       didact.object = chgb.par.exp, 
                       col.range = c("white", "dodgerblue4"), 
                       border.col = "black", 
                       include.var.pie = TRUE,
                       median.line.col = "red", 
                       density.col = "white")
title("NZO (A) x WSB (B)")

make.single.cross.plot(cross = "CASTxWSB",
                       cross.type = "f2",
                       utility.type = "contrasts",
                       didact.object = chgb.par.exp, 
                       col.range = c("white", "dodgerblue4"), 
                       border.col = "black", 
                       include.var.pie = TRUE,
                       median.line.col = "red", 
                       density.col = "white")
title("CAST (A) x WSB (B)")

make.gradient.scale(spectrum = make.spectrum(c("white", "dodgerblue4"), n = 100),
                    mode = "vertical", 
                    utility.type = "contrasts")

##############################################################################
########  
########  Figure S3: DIDACT POE BC
########
##############################################################################
diallelPlotter(results = chgb.par.exp, 
               cross.type = "rbc1", 
               pheno.name = "chgb", 
               utility.type = "contrasts", 
               include.bar.plots = FALSE, 
               include.density = FALSE, 
               include.rank = TRUE,
               include.info.plot = FALSE, 
               col.range = c("white", "dodgerblue4"),
               label.cex = 2,
               label.padj = -0.5)
diallelPlotter(results = chgb.par.exp, 
               cross.type = "rbc2", 
               pheno.name = "chgb", 
               utility.type = "contrasts", 
               include.bar.plots = FALSE, 
               include.density = FALSE, 
               include.rank = TRUE,
               include.info.plot = FALSE, 
               col.range = c("white", "dodgerblue4"),
               label.cex = 2,
               label.padj = -0.5)
make.gradient.scale(spectrum = make.spectrum(c("white", "dodgerblue4"), n = 100),
                    mode = "vertical", 
                    utility.type = "contrasts")
```
