## Supplementary material for "Bayesian decision theoretic design of two-founder experimental crosses given diallel data"

```
##############################################################################
##############################################################################
########
########  Title: sim_biallelic_add.R
########  Purpose: Simulate biallelic additive diallel samples and DIDACT analysis
########  Authors: Greg Keele
########
##############################################################################
##############################################################################

library(DIDACT)

sim <- 1
## Run for 1:100

run.didact.sims <- function(add.effect,
                            mu = 5, 
                            add.size, 
                            inbred.mu = 0, 
                            inbred.size = 0, 
                            epi.sym.size = 0,
                            epi.asym.size = 0,
                            maternal.size = 0) {
  counter <- 0
  results <- list()
  for (i in add.size) {
    this.sim <- simulate.diallel(add.effect = add.effect,
                                 mu = mu, 
                                 add.size = i, 
                                 inbred.mu = inbred.mu, 
                                 inbred.size = inbred.size, 
                                 epi.sym.size = epi.sym.size, 
                                 epi.asym.size = epi.asym.size,
                                 maternal.size = maternal.size, 
                                 num.sim = 1)
    this.bayesdiallel <- diallel.gibbs(phenotype = this.sim$diallel.data$y.sim.1, 
                                       sex = this.sim$diallel.data$is.female, 
                                       mother.str = this.sim$diallel.data$dam.id, 
                                       father.str = this.sim$diallel.data$sire.id, 
                                       strains.reorder = LETTERS[1:8], 
                                       strains.rename = LETTERS[1:8],
                                       n.iter = 500, 
                                       burn.in = 10000, 
                                       use.constraint = TRUE)
    this.didact <- evaluate.experiments(gibbs.object = this.bayesdiallel, 
                                        n = 50, 
                                        qtl.num = 1,
                                        use.progress.bar = FALSE)
    this.oracle.didact <- evaluate.experiments(gibbs.object = this.sim$didact.input, 
                                               n = 50, 
                                               qtl.num = 1,
                                               use.progress.bar = FALSE)
    
    counter <- counter + 1
    results[[counter]] <- list(sim = this.sim,
                               bayesdiallele = this.bayesdiallel,
                               didact = this.didact,
                               oracle.didact = this.oracle.didact,
                               effect = i)
  }
  return(results)
}

biallelic.results <- run.didact.sims(add.effect = c(rep(-1, 4), rep(1, 4)),
                                     add.size = seq(0.01, 0.99, length.out = 50))

dir.create("results/biallelic_add", recursive = TRUE, showWarnings = TRUE)
saveRDS(biallelic.results, paste0("./results/biallelic_add/sim", sim, ".RDS"))

####### Plot results from 100 simulations
add.mse.matrix <- matrix(NA, nrow = 50, ncol = 100)
rank.cor.matrix <- rank.cor.power.matrix <- matrix(NA, nrow = 50, ncol = 100)
for (i in 1:100) {
  this.sim <- readRDS(paste0("./results/biallelic_add/sim", i, ".RDS"))
  for (j in 1:50) {
    add.mse.matrix[j, i] <- sum((colMeans(this.sim[[j]]$bayesdiallele$mcmc[, paste("add", LETTERS[1:8], sep = ":")]) - this.sim[[j]]$sim$effects$add)^2)

    rank.cor.matrix[j, i] <- cor(rank(unlist(this.sim[[j]]$oracle.didact$var[[cross.type]]), ties.method="min"),
                                 rank(unlist(lapply(this.sim[[j]]$didact$var[[cross.type]], function(x) mean(x))), ties.method="min"))
  }
}
rank.cor.power.matrix[is.na(rank.cor.power.matrix)] <- 1
effect.size <- seq(0.01, 0.99, length.out=50)

## MSE
## Figure 3A: MSE for biallelic_add
plot(1-effect.size, rowMeans(add.mse.matrix), xlab="Proportion of variance from noise", ylab="MSE",
     main=sim.model, ylim=c(0, max(add.mse.matrix)), las=1, frame.plot=FALSE)
for (i in 2:100){
  points(1-effect.size, add.mse.matrix[,i], pch=20, col=scales::alpha("gray", 0.5))
}
points(1-effect.size, rowMeans(add.mse.matrix))

## Rank posterior contrasts
## Figure 3D: Rank utility for biallelic_add
plot(1-effect.size, rowMeans(rank.cor.matrix), xlab="Proportion of variance from noise", ylab="Rank correlation of utility",
     main=sim.model, ylim=c(min(rank.cor.matrix), 1), las=1, frame.plot=FALSE)
for (i in 2:100){
  points(1-effect.size, rank.cor.matrix[,i], pch=20, col=scales::alpha("gray", 0.5))
}
points(1-effect.size, rowMeans(rank.cor.matrix))
abline(h=0.85769, lty=2, col="red")
```
