## Supplementary material for "Bayesian decision theoretic design of two-founder experimental crosses given diallel data"

**Supplemental Tables and Figures**

**Table S1 Model of QTL effect on the mean for F2 and BC**

| Genotype | $\mathrm{E}(y_i)^b$ | $\mathbf{x}_i{}^c$ | Probability[a] | | |
|---|---|---|---|---|---|
| | | | F2 | $\mathrm{BC}_A$ | $\mathrm{BC}_B$ |
| AA | $\mu + \alpha - \frac{\delta}{2}$ | $\begin{bmatrix} 1 & 1 & -\frac{1}{2} \end{bmatrix}$ | $\frac{1}{4}$ | $\frac{1}{2}$ | $0$ |
| AB | $\mu + \frac{\delta}{2}$ | $\begin{bmatrix} 1 & 0 & \frac{1}{2} \end{bmatrix}$ | $\frac{1}{2}$ | $\frac{1}{2}$ | $\frac{1}{2}$ |
| BB | $\mu - \alpha - \frac{\delta}{2}$ | $\begin{bmatrix} 1 & -1 & -\frac{1}{2} \end{bmatrix}$ | $\frac{1}{4}$ | $0$ | $\frac{1}{2}$ |

[a] Mendelian inheritance probabilities based on independent assortment of alleles $A$ and $B$ for specified bi-parental cross.
[b] Parameters as defined in **Eq 3**, **Eq 4**, and **Eq 5**.
[c] $i^{\text{th}}$ row vector of the design matrix $\mathbf{X}$ in **Eq 3**.

**Table S2 Variance attributable to QTL effect for F2 and BC**

| Model | Parameter[a] | F2 | $\mathrm{BC}_A$ | $\mathrm{BC}_B$ |
|---|---|---|---|---|
| General | | $\frac{1}{4}\delta^2 + \frac{1}{2}\alpha^2$ | $\frac{1}{4}(\alpha + \delta)^2$ | $\frac{1}{4}(\alpha + \delta)^2$ |
| Fully additive | $\delta = 0$ | $\frac{1}{2}\alpha^2$ | $\frac{1}{4}\alpha^2$ | $\frac{1}{4}\alpha^2$ |
| A dominant | $\delta = \alpha$ | $\frac{3}{4}\alpha^2$ | $0$ | $\alpha^2$ |
| B dominant | $\delta = -\alpha$ | $\frac{3}{4}\alpha^2$ | $\alpha^2$ | $0$ |
| Fully dominant | $\alpha = 0$ | $\frac{1}{4}\delta^2$ | $\frac{1}{4}\delta^2$ | $\frac{1}{4}\delta^2$ |

[a] Parameters as defined in **Eq 3**, **Eq 4**, and **Eq 5**.

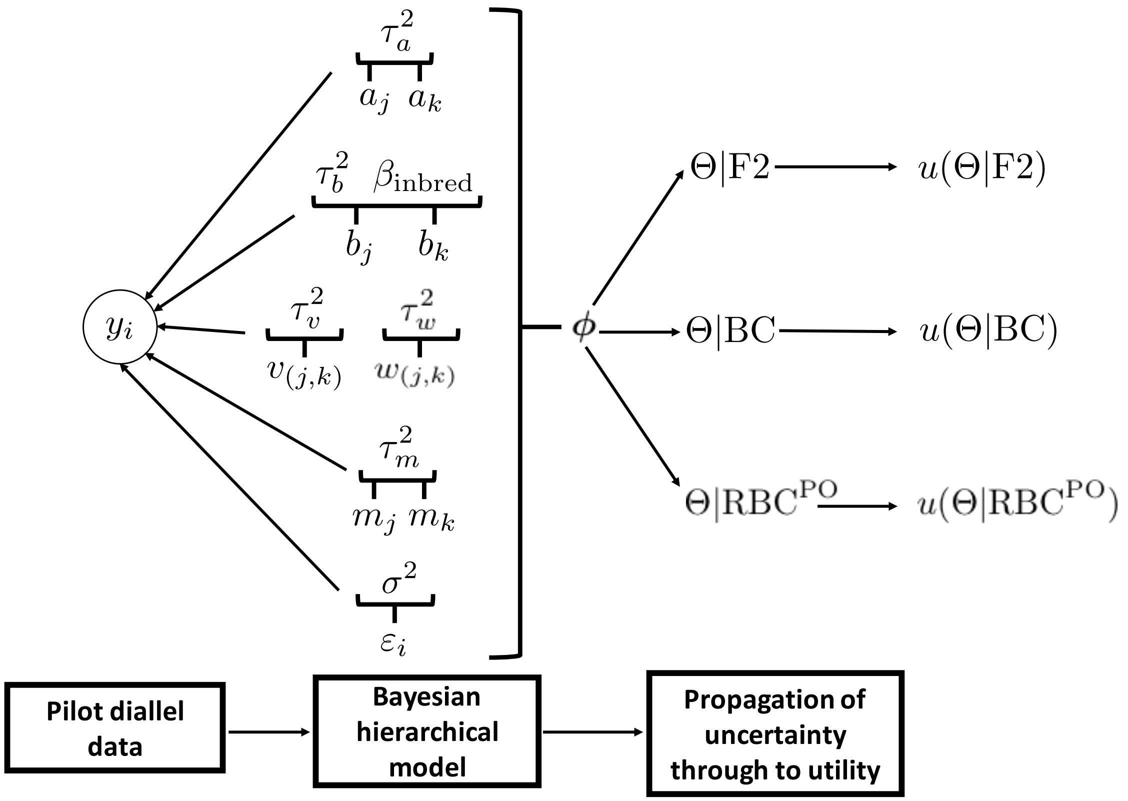


**Figure S1** The Bayesian hierarchical model underlying BayesDiallel that is fit within DIDACT. Strain-level effects are characterized and propagated to a utility function. The diagram represents a single sample from a Gibbs sampler, though the decision theoretic approach is compatible generally with other MCMC procedures. Strain-level effects are sampled from pilot diallel data, collectively defined as $\boldsymbol{\phi}$. Strain-level effects $\boldsymbol{\phi}$ are mapped to an effect from a single putative QTL $\theta|\mathfrak{a}$, using **Eq 4** and **Eq 5** as well as **Tables S1** and **S2**, where $\theta$ representing the effect of a single putative QTL in a bi-parental cross and $\mathfrak{a}$, the specific cross of two specific founder strains. $\Theta|\text{F2}$ collectively refers to all $\theta$ from the possible F2 crosses, with similar notation for BC and $\text{RBC}^{\text{PO}}$. Effectively $\Theta$ are functions of $\boldsymbol{\phi}$, which are then used as inputs into the utility function $u(.)$, in our case, a power estimate for a single putative QTL. This process is repeated for $s$ samples from the MCMC procedure, allowing for posterior estimates of utility.

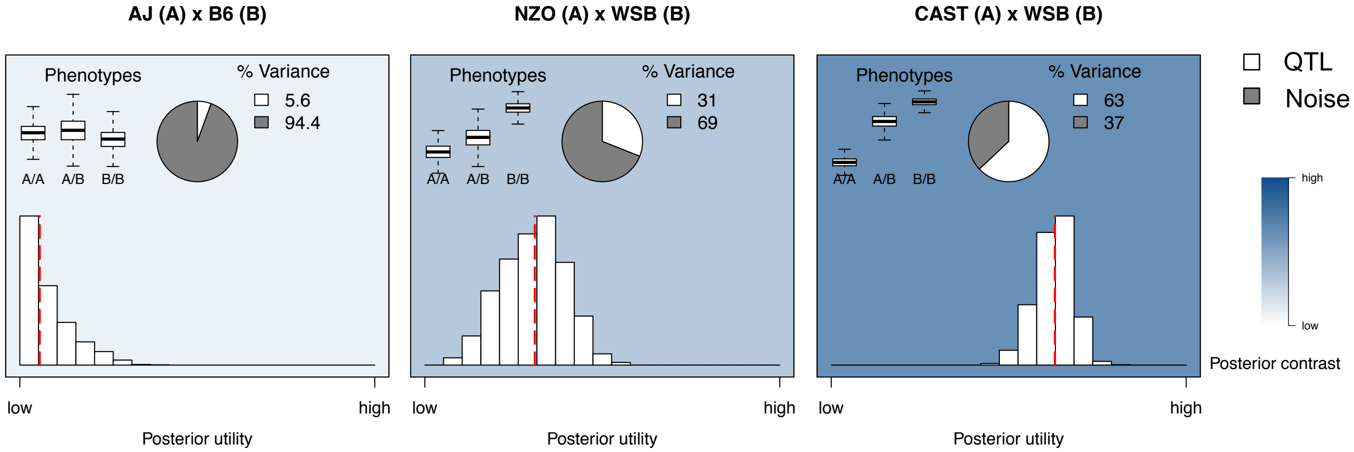


**Figure S2** Detailed summary plots of select F2 crosses from DIDACT for cHGB. The full panel of potential F2 crosses are presented in **Figure 6C**. DIDACT allows for additional information to be overlayed on the posterior mean of the utility, which is represented by the background color. These plots include the histogram of the posterior distribution of the utility function, in this case phenotype contrasts, the posterior median utility as a red dashed line, the posterior median variance explained by the strain-level effects (white) as a pie chart with point estimates, and posterior five point summaries of the predicted phenotypes per diallel cell. For example, the strain-level effects account for 63.2% of the phenotypic variance observed in the WSB, CAST, and their hybrid offspring. A CAST mouse is expected to have a low phenotype, a WSB mouse to have a high phenotype, and the hybrid intermediate with some dominance towards WSB. These estimates match the raw data in **Figure 6A**.

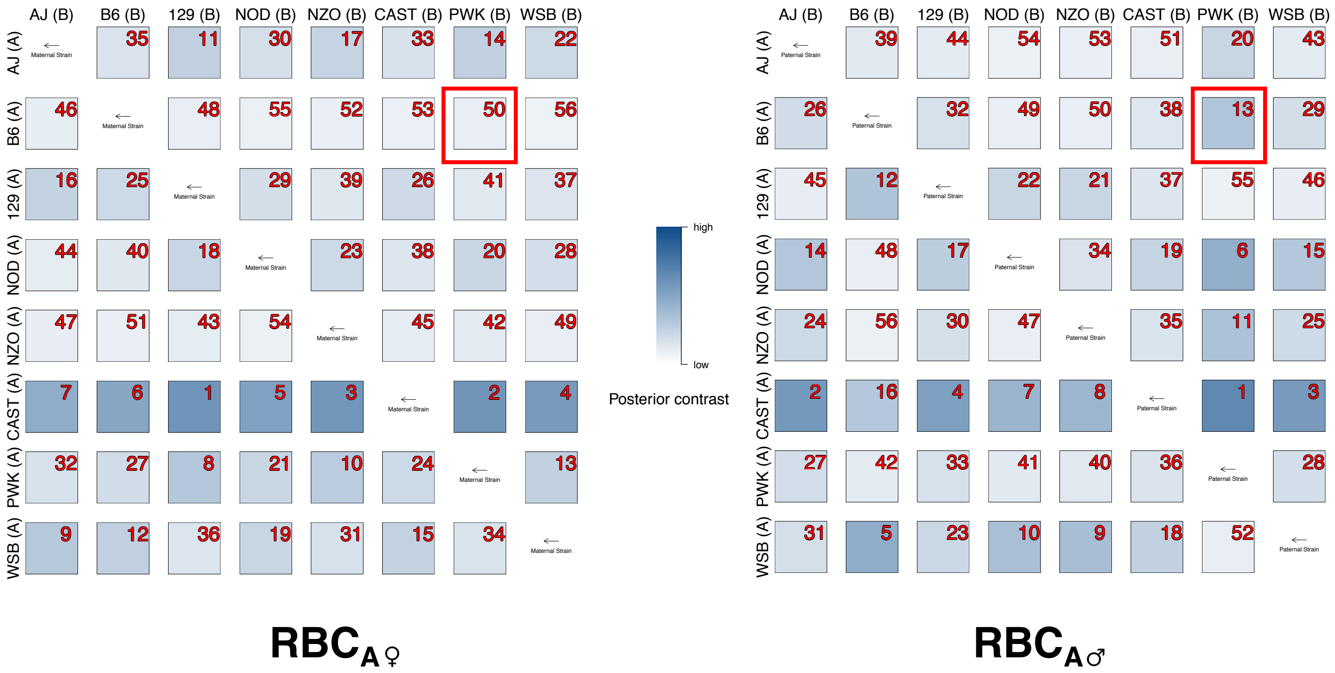


**Figure S3** A panel of DIDACT posterior contrasts for all possible BC in which the maternal and paternal strain identities of the F1 and backcrossed generation are fixed. Red numbers in the upper right corners represent the cross ranking of posterior mean contrasts within each panel. Though not a direct calculation of power for the RBC$^{PO}$ design, this approach highlights the potential that strain-level maternal effects can contribute to differences in predicted RBC$^{PO}$. The B6 $\times$ PWK BC are marked with red squares, for which DIDACT predicts the BC with the backcrossed B6 as the sire (lower - ranked 13$^{th}$) as having greater phenotype contrasts than when the backcrossed B6 is the dam (upper - ranked 50$^{th}$).
