## Supplementary material for "Bayesian decision theoretic design of two-founder experimental crosses given diallel data": DESCRIPTION

Package: DIDACT
Title: Diallel Informed Decision theoretic Approach for Crosses Tool
        (DIDACT)
Version: 1.1.1
Authors@R: c(person("Paul", "Maurizio", role = "aut"), 
             person("Daniel", "Oreper", role = "aut"),
	     person("William", "Valdar", role = "aut"),  
Description: DIDACT models a diallel experiment, the set of inbred founders and their F1 hybrids, with a Bayesian hierarchical model. The Monte Carlo (MC) sampling is extended to decision theoretic utility functions that researchers seek to maximize or minimize. One such utility function could be the power to map a QTL in a downstream F2 intercross or back cross (BC). In this framework, signal and uncertainty can be characterized in the training diallel data, and intuitively incorporated into the experimental design step of follow-up experiments.
Depends: R (>= 3.3.2)
License: GPL (>= 2)
Encoding: UTF-8
LazyData: true
RoxygenNote: 6.1.0
Imports: mnormt, coda, qtlDesign, colorRamps, mapplots
Suggests:
NeedsCompilation: no
Packaged: 2018-12-06 21:28:33 UTC; Greg
Author: Paul Maurizio [aut],
  Daniel Oreper [aut],
  William Valdar [aut],
  Gregory Keele [aut, cre]
Maintainer: Gregory Keele <>
