## Supplementary material for "Bayesian decision theoretic design of two-founder experimental crosses given diallel data": decision_functions.R

```
# Calculating QTL effects based on crosses
calc.f2.effects <- function(line.m, 
                            line.p, 
                            par.vec, 
                            qtl.num = 1,
                            strain.order = c("WSB", "PWK", "CAST", "NZO", "NOD", "129", "B6", "AJ")){
  m.a.effect <- par.vec[paste("add", line.m, sep=":")]
  p.a.effect <- par.vec[paste("add", line.p, sep=":")]
  i.effect <- par.vec["inbred penalty"]
  m.i.effect <- par.vec[paste("inbred", line.m, sep=":")]
  p.i.effect <- par.vec[paste("inbred", line.p, sep=":")]
  # Epistatic effects
  m.index <- which(line.m == strain.order)
  p.index <- which(line.p == strain.order)
  epi.indeces <- sort(c(m.index, p.index))
  if (m.index == p.index) {
    e.sym.effect <- par.vec[paste("epi_sym", paste(strain.order[epi.indeces[1]], strain.order[epi.indeces[2]], sep=";"), sep=":")]
    e.asym.effect <- par.vec[paste("epi_asym", paste(strain.order[epi.indeces[1]], strain.order[epi.indeces[2]], sep=";"), sep=":")]
  }
  else {
    e.sym.effect <- 0
    e.asym.effect <- 0
  }
  homo.m <- 2*m.a.effect + i.effect + m.i.effect
  homo.p <- 2*p.a.effect + i.effect + p.i.effect
  hetero <- m.a.effect + p.a.effect + e.sym.effect + e.asym.effect
  add <- (homo.m - homo.p)/2
  dom <- hetero - (homo.m + homo.p)/2
  effects <- c(add/qtl.num, dom/qtl.num)
  return(effects)
}

calc.bc.effects <- function(re.cross.line, 
                            other.line, 
                            par.vec, 
                            qtl.num = 1,
                            strain.order = c("WSB", "PWK", "CAST", "NZO", "NOD", "129", "B6", "AJ")){
  re.cross.a.effect <- par.vec[paste("add", re.cross.line, sep=":")]
  other.line.a.effect <- par.vec[paste("add", other.line, sep=":")]
  i.effect <- par.vec["inbred penalty"]
  re.cross.i.effect <- par.vec[paste("inbred", re.cross.line, sep=":")]
  # Epistatic effects
  re.cross.index <- which(re.cross.line == strain.order)
  other.index <- which(other.line == strain.order)
  epi.indeces <- sort(c(re.cross.index, other.index))
  if (re.cross.index == other.index) {
    e.sym.effect <- par.vec[paste("epi_sym", paste(strain.order[epi.indeces[1]], strain.order[epi.indeces[2]], sep=";"), sep=":")]
    e.asym.effect <- par.vec[paste("epi_asym", paste(strain.order[epi.indeces[1]], strain.order[epi.indeces[2]], sep=";"), sep=":")]
  }
  else {
    e.sym.effect <- 0
    e.asym.effect <- 0
  }
  homo.re <- 2*re.cross.a.effect + i.effect + re.cross.i.effect
  hetero <- re.cross.a.effect + other.line.a.effect + e.sym.effect + e.asym.effect
  add.dom <- hetero - homo.re
  effects <- add.dom/qtl.num
  return(effects)
}

calc.rbc.effects <- function(re.cross.line, 
                             other.line, 
                             mat.line, 
                             par.vec, 
                             qtl.num = 1,
                             strain.order = c("WSB", "PWK", "CAST", "NZO", "NOD", "129", "B6", "AJ")){
  # Determining paternal line
  if (mat.line == re.cross.line) {
    pat.line <- other.line
  }
  else {
    pat.line <- re.cross.line
  }
  re.cross.a.effect <- par.vec[paste("add", re.cross.line, sep=":")]
  other.line.a.effect <- par.vec[paste("add", other.line, sep=":")]
  i.effect <- par.vec["inbred penalty"]
  re.cross.i.effect <- par.vec[paste("inbred", re.cross.line, sep=":")]
  mat.effect <- par.vec[paste("mat", mat.line, sep=":")]
  pat.effect <- par.vec[paste("mat", pat.line, sep=":")]
  # Epistatic effects
  re.cross.index <- which(re.cross.line == strain.order)
  other.index <- which(other.line == strain.order)
  epi.indeces <- sort(c(re.cross.index, other.index))
  if (re.cross.index == other.index) {
    e.sym.effect <- par.vec[paste("epi_sym", paste(strain.order[epi.indeces[1]], strain.order[epi.indeces[2]], sep=";"), sep=":")]
    e.asym.effect <- par.vec[paste("epi_asym", paste(strain.order[epi.indeces[1]], strain.order[epi.indeces[2]], sep=";"), sep=":")]
  }
  else {
    e.sym.effect <- 0
    e.asym.effect <- 0
  }
  homo.re <- 2*re.cross.a.effect + i.effect + re.cross.i.effect
  hetero <- re.cross.a.effect + other.line.a.effect + mat.effect - pat.effect + e.sym.effect + e.asym.effect
  add.dom <- hetero - homo.re
  effects <- add.dom/qtl.num
  return(effects)
}

calc.phenotypes.f2 <- function(line.m, 
                               line.p, 
                               par.vec, 
                               qtl.num = 1,
                               strain.order = c("WSB", "PWK", "CAST", "NZO", "NOD", "129", "B6", "AJ")){
  m.a.effect <- par.vec[paste("add", line.m, sep=":")]
  p.a.effect <- par.vec[paste("add", line.p, sep=":")]
  m.i.effect <- par.vec[paste("inbred", line.m, sep=":")]
  p.i.effect <- par.vec[paste("inbred", line.p, sep=":")]
  # Epistatic effects
  m.index <- which(line.m == strain.order)
  p.index <- which(line.p == strain.order)
  epi.indeces <- sort(c(m.index, p.index))
  if (m.index == p.index) {
    e.sym.effect <- par.vec[paste("epi_sym", paste(strain.order[epi.indeces[1]], strain.order[epi.indeces[2]], sep=";"), sep=":")]
    e.asym.effect <- par.vec[paste("epi_asym", paste(strain.order[epi.indeces[1]], strain.order[epi.indeces[2]], sep=";"), sep=":")]
  }
  else {
    e.sym.effect <- 0
    e.asym.effect <- 0
  }
  homo.m <- par.vec["mu"] + 2*m.a.effect + par.vec["inbred penalty"] + m.i.effect
  homo.p <- par.vec["mu"] + 2*p.a.effect + par.vec["inbred penalty"] + p.i.effect
  hetero <- par.vec["mu"] + m.a.effect + p.a.effect + e.sym.effect + e.asym.effect
  phenos <- cbind(homo.m, hetero, homo.p)
  colnames(phenos) <- c("f2-hom1", "f2-het", "f2-hom2")
  return(phenos)
}

calc.phenotypes.bc <- function(re.cross.line, 
                               other.line, 
                               par.vec, 
                               qtl.num = 1, 
                               cross.type = "bc1",
                               strain.order = c("WSB", "PWK", "CAST", "NZO", "NOD", "129", "B6", "AJ")){
  re.cross.a.effect <- par.vec[paste("add", re.cross.line, sep=":")]
  other.a.effect <- par.vec[paste("add", other.line, sep=":")]
  re.cross.i.effect <- par.vec[paste("inbred", re.cross.line, sep=":")]
  # Epistatic effects
  re.cross.index <- which(re.cross.line == strain.order)
  other.index <- which(other.line == strain.order)
  epi.indeces <- sort(c(re.cross.index, other.index))
  if (re.cross.index == other.index) {
    e.sym.effect <- par.vec[paste("epi_sym", paste(strain.order[epi.indeces[1]], strain.order[epi.indeces[2]], sep=";"), sep=":")]
    e.asym.effect <- par.vec[paste("epi_asym", paste(strain.order[epi.indeces[1]], strain.order[epi.indeces[2]], sep=";"), sep=":")]
  }
  else {
    e.sym.effect <- 0
    e.asym.effect <- 0
  }
  homo.re <- par.vec["mu"] + 2*re.cross.a.effect + par.vec["inbred penalty"] + re.cross.i.effect
  hetero <- par.vec["mu"] + re.cross.a.effect + other.a.effect + e.sym.effect + e.asym.effect
  phenos <- cbind(homo.re, hetero)
  if (cross.type == "bc1") {
    colnames(phenos) <- c("bc1-hom", "bc1-het")
  }
  else {
    colnames(phenos) <- c("bc2-hom", "bc2-het")
  }
  return(phenos)
}  

calc.phenotypes.rbc <- function(re.cross.line, 
                                other.line, 
                                mat.line, 
                                par.vec, 
                                qtl.num = 1, 
                                cross.type = "rbc1_1",
                                strain.order = c("WSB", "PWK", "CAST", "NZO", "NOD", "129", "B6", "AJ")){
  # Determining paternal line
  if (mat.line == re.cross.line) {
    pat.line <- other.line
  }
  else {
    pat.line <- re.cross.line
  }
  re.cross.a.effect <- par.vec[paste("add", re.cross.line, sep=":")]
  other.a.effect <- par.vec[paste("add", other.line, sep=":")]
  re.cross.i.effect <- par.vec[paste("inbred", re.cross.line, sep=":")]
  # Epistatic effects
  re.cross.index <- which(re.cross.line == strain.order)
  other.index <- which(other.line == strain.order)
  epi.indeces <- sort(c(re.cross.index, other.index))
  if (re.cross.index == other.index) {
    e.sym.effect <- par.vec[paste("epi_sym", paste(strain.order[epi.indeces[1]], strain.order[epi.indeces[2]], sep=";"), sep=":")]
    e.asym.effect <- par.vec[paste("epi_asym", paste(strain.order[epi.indeces[1]], strain.order[epi.indeces[2]], sep=";"), sep=":")]
  }
  else {
    e.sym.effect <- 0
    e.asym.effect <- 0
  }
  mat.effect <- par.vec[paste("mat", mat.line, sep=":")]
  pat.effect <- par.vec[paste("mat", pat.line, sep=":")]
  homo.re <- par.vec["mu"] + 2*re.cross.a.effect + par.vec["inbred penalty"] + re.cross.i.effect
  hetero <- par.vec["mu"] + re.cross.a.effect + other.a.effect + mat.effect - pat.effect + e.sym.effect + e.asym.effect
  phenos <- cbind(homo.re, hetero)
  if (cross.type=="rbc1_1") {
    colnames(phenos) <- c("rbc1_1-hom", "rbc1_1-het")
  }
  else if (cross.type=="rbc1_2") {
    colnames(phenos) <- c("rbc1_2-hom", "rbc1_2-het")
  }
  else if (cross.type=="rbc2_1") {
    colnames(phenos) <- c("rbc2_1-hom", "rbc2_1-het")
  }
  else if (cross.type=="rbc2_2") {
    colnames(phenos) <- c("rbc2_2-hom", "rbc2_2-het")
  }
  return(phenos)
}

power.cruncher.general <- function(line.m, 
                                   line.p, 
                                   cross.type, 
                                   par.vec, 
                                   n, 
                                   re.cross = NULL, 
                                   mat.line = NULL, 
                                   qtl.num = 1,
                                   strains = c("AJ", "B6", "129", "NOD", "NZO", "CAST", "PWK", "WSB")){
  # Power calculation
  if(cross.type == "f2"){
    f2.effects <- calc.f2.effects(line.m=line.m, line.p=line.p, par.vec=par.vec, qtl.num=qtl.num, strain.order=rev(strains))
    power.val <- qtlDesign::powercalc(cross=cross.type, n=n, effect=f2.effects, sigma2=par.vec["sigma2"])[1]
  }
  else if (cross.type == "bc") {
    if(re.cross == line.m){
      other.line <- line.p
    }
    else{
      other.line <- line.m
    }
    bc.effects <- calc.bc.effects(re.cross.line=re.cross, other.line=other.line, par.vec=par.vec, qtl.num=qtl.num, strain.order=rev(strains))
    power.val <- qtlDesign::powercalc(cross=cross.type, n=n, effect=bc.effects, sigma2=par.vec["sigma2"])[1]
  }
  else if(cross.type == "rbc"){
    if(re.cross == line.m){
      other.line <- line.p
    }
    else{
      other.line <- line.m
    }
    rbc.effects <- calc.rbc.effects(re.cross.line=re.cross, 
                                    other.line=other.line, 
                                    mat.line=mat.line, 
                                    par.vec=par.vec, 
                                    qtl.num=qtl.num, 
                                    strain.order=rev(strains))
    power.val <- qtlDesign::powercalc(cross="bc", 
                                      n=n, 
                                      effect=rbc.effects, 
                                      sigma2=par.vec["sigma2"])[1]
  }
  return(power.val)
}

power.matcher.general <- function(par.vec, 
                                  n, 
                                  qtl.num = 1, 
                                  strains = c("AJ", "B6", "129", "NOD", "NZO", "CAST", "PWK", "WSB")){
  # Initialize data objects
  cross.names <- NULL
  powers <- NULL
  strain.matrix <- NULL
  mover <- 2 # Keeps inbred (NZOxNZO) and identical (NZOxB6 and B6xNZ) crosses from occurring
  for (i in 1:(length(strains)-1)) {
    for (j in mover:length(strains)) {
      # Cross types
      bc.cross1 <- power.cruncher.general(line.m=strains[i], 
                                          line.p=strains[j], 
                                          cross.type="bc", 
                                          par.vec=par.vec, 
                                          n=n, 
                                          re.cross=strains[i], 
                                          qtl.num=qtl.num, 
                                          strains=strains)
      rbc.cross1_1 <- power.cruncher.general(line.m=strains[i], 
                                             line.p=strains[j], 
                                             cross.type="rbc",
                                             par.vec=par.vec, 
                                             n=n, 
                                             re.cross=strains[i], 
                                             mat.line=strains[i], 
                                             qtl.num=qtl.num, 
                                             strains=strains)
      rbc.cross1_2 <- power.cruncher.general(line.m=strains[i], 
                                             line.p=strains[j], 
                                             cross.type="rbc", 
                                             par.vec=par.vec, 
                                             n=n, 
                                             re.cross=strains[i], 
                                             mat.line=strains[j], 
                                             qtl.num=qtl.num, 
                                             strains=strains)      
      bc.cross2 <- power.cruncher.general(line.m=strains[i], 
                                          line.p=strains[j], 
                                          cross.type="bc", 
                                          par.vec=par.vec, 
                                          n=n, 
                                          re.cross=strains[j], 
                                          qtl.num=qtl.num, 
                                          strains=strains)
      rbc.cross2_1 <- power.cruncher.general(line.m=strains[i], 
                                             line.p=strains[j], 
                                             cross.type="rbc", 
                                             par.vec=par.vec, 
                                             n=n, 
                                             re.cross=strains[j], 
                                             mat.line=strains[i], 
                                             qtl.num=qtl.num, 
                                             strains=strains)
      rbc.cross2_2 <- power.cruncher.general(line.m=strains[i], 
                                             line.p=strains[j], 
                                             cross.type="rbc", 
                                             par.vec=par.vec, 
                                             n=n, 
                                             re.cross=strains[j], 
                                             mat.line=strains[j], 
                                             qtl.num=qtl.num, 
                                             strains=strains)
      f2.cross <- power.cruncher.general(line.m=strains[i], 
                                         line.p=strains[j], 
                                         cross.type="f2", 
                                         par.vec=par.vec, 
                                         n=n, 
                                         qtl.num=qtl.num, 
                                         strains=strains)
      
      # Processing output
      cross.names <- paste(strains[i], "x", strains[j], sep="")
      powers <- cbind(bc.cross1, rbc.cross1_1, rbc.cross1_2, bc.cross2, rbc.cross2_1, rbc.cross2_2, f2.cross)
      colnames(powers) <- c("bc1", "rbc1_1", "rbc1_2", "bc2", "rbc2_1", "rbc2_2", "f2")
      rownames(powers) <- cross.names
      strain.matrix <- rbind(strain.matrix, powers)
    }
    mover <- mover + 1
  }
  return(strain.matrix)
}

## Calculate variance explained
calc.var.exp.general <- function(line.m, 
                                 line.p, 
                                 cross.type, 
                                 par.vec, 
                                 re.cross = NULL, 
                                 mat.line = NULL, 
                                 qtl.num = 1,
                                 strains = c("AJ", "B6", "129", "NOD", "NZO", "CAST", "PWK", "WSB")){
  if (cross.type == "f2") {
    effects <- calc.f2.effects(line.m=line.m, 
                               line.p=line.p, 
                               par.vec=par.vec, 
                               qtl.num=qtl.num, 
                               strain.order=rev(strains))
    qtl.var <- (1/2)*effects[1]^2 + (1/4)*effects[2]^2
    
    phenotypes <- calc.phenotypes.f2(line.m=line.m, 
                                     line.p=line.p, 
                                     par.vec=par.vec, 
                                     qtl.num=qtl.num, 
                                     strain.order=rev(strains))
  }
  else if (cross.type == "bc") {
    if (re.cross == line.m) {
      other.line <- line.p
      type <- "bc1"
    }
    else {
      other.line <- line.m
      type <- "bc2"
    }
    effects <- calc.bc.effects(re.cross.line=re.cross, 
                               other.line=other.line, 
                               par.vec=par.vec, 
                               qtl.num=qtl.num, 
                               strain.order=rev(strains))
    qtl.var <- (1/4)*(effects)^2
    
    phenotypes <- calc.phenotypes.bc(re.cross.line=re.cross, 
                                     other.line=other.line, 
                                     par.vec=par.vec, 
                                     qtl.num=qtl.num, 
                                     cross.type=type, 
                                     strain.order=rev(strains))
  }
  else if (cross.type == "rbc") {
    if (re.cross == line.m) {
      other.line <- line.p
      if (mat.line == line.m) {
        type <- "rbc1_1"
      }
      else {
        type <- "rbc1_2"
      }
    }
    else {
      other.line <- line.m
      if (mat.line == line.m) {
        type <- "rbc2_1"
      }
      else{
        type <- "rbc2_2"
      }
    } 
    effects <- calc.rbc.effects(re.cross.line=re.cross, 
                                other.line=other.line, 
                                mat.line=mat.line, 
                                par.vec=par.vec, 
                                qtl.num=qtl.num,
                                strain.order=rev(strains))
    qtl.var <- (1/4)*(effects)^2
    
    phenotypes <- calc.phenotypes.rbc(re.cross.line = re.cross, 
                                      other.line = other.line, 
                                      mat.line = mat.line, 
                                      par.vec = par.vec, 
                                      qtl.num = qtl.num, 
                                      cross.type = type,
                                      strain.order = rev(strains))
  }
  perc.var <- 100*qtl.var*qtl.num/(qtl.var*qtl.num + par.vec["sigma2"])
  return(list(phenotypes, as.numeric(perc.var)))
}

## Calculates variance explained by QTL for a given draw 
## from the Gibbs sampler for all possible crosses
var.matcher.general <- function(par.vec, 
                                qtl.num = 1, 
                                strains = c("AJ", "B6", "129", "NOD", "NZO", "CAST", "PWK", "WSB")){
  # 8 founder lines of CC are default
  # Initialize data objects
  cross.names <- NULL
  vars <- NULL
  var.matrix <- NULL
  phenotype.matrix <- NULL
  mover <- 2 # Keeps inbred (NZOxNZO) and identical (NZOxB6 and B6xNZ) crosses from occurring
  for (i in 1:(length(strains)-1)) {
    for (j in mover:length(strains)) {
      # Cross types
      bc.cross1 <- calc.var.exp.general(line.m = strains[i], 
                                        line.p = strains[j], 
                                        cross.type = "bc", 
                                        par.vec = par.vec, 
                                        re.cross = strains[i], 
                                        qtl.num = qtl.num, 
                                        strains = strains)
      rbc.cross1_1 <- calc.var.exp.general(line.m = strains[i], 
                                           line.p = strains[j], 
                                           cross.type = "rbc", 
                                           par.vec = par.vec, 
                                           re.cross = strains[i], 
                                           mat.line = strains[i], 
                                           qtl.num = qtl.num, 
                                           strains = strains)
      rbc.cross1_2 <- calc.var.exp.general(line.m = strains[i], 
                                           line.p = strains[j], 
                                           cross.type = "rbc", 
                                           par.vec = par.vec, 
                                           re.cross = strains[i], 
                                           mat.line = strains[j], 
                                           qtl.num = qtl.num, 
                                           strains = strains)

      bc.cross2 <- calc.var.exp.general(line.m = strains[i], 
                                        line.p = strains[j], 
                                        cross.type = "bc", 
                                        par.vec = par.vec, 
                                        re.cross = strains[j], 
                                        qtl.num = qtl.num, 
                                        strains = strains)
      rbc.cross2_1 <- calc.var.exp.general(line.m = strains[i], 
                                           line.p = strains[j], 
                                           cross.type = "rbc", 
                                           par.vec = par.vec, 
                                           re.cross = strains[j], 
                                           mat.line = strains[i], 
                                           qtl.num = qtl.num, 
                                           strains = strains)
      rbc.cross2_2 <- calc.var.exp.general(line.m = strains[i], 
                                           line.p = strains[j], 
                                           cross.type = "rbc", 
                                           par.vec = par.vec, 
                                           re.cross = strains[j], 
                                           mat.line = strains[j], 
                                           qtl.num = qtl.num, 
                                           strains = strains)
      
      f2.cross <- calc.var.exp.general(line.m = strains[i], 
                                       line.p = strains[j], 
                                       cross.type = "f2", 
                                       par.vec = par.vec, 
                                       qtl.num = qtl.num, 
                                       strains = strains)
      
      # Processing output
      cross.names <- paste(strains[i], "x", strains[j], sep="")
      cross.phenotypes <- cbind(bc.cross1[[1]], rbc.cross1_1[[1]], rbc.cross1_2[[1]], 
                                bc.cross2[[1]], rbc.cross2_1[[1]], rbc.cross2_2[[1]], f2.cross[[1]])
      rownames(cross.phenotypes) <- cross.names
      phenotype.matrix <- rbind(phenotype.matrix, cross.phenotypes)
     
      vars <- cbind(bc.cross1[[2]], rbc.cross1_1[[2]], rbc.cross1_2[[2]], 
                        bc.cross2[[2]], rbc.cross2_1[[2]], rbc.cross2_2[[2]], f2.cross[[2]])
      colnames(vars) <- c("bc1_perc", "rbc1_1_perc", "rbc1_2_perc", "bc2_perc", "rbc2_1_perc", "rbc2_2_perc", "f2_perc")
      rownames(vars) <- cross.names
      var.matrix <- rbind(var.matrix, vars)
    }
    mover <- mover + 1
  }
  return(list(phenotype.matrix, var.matrix))
}

## Runs power.matcher and var.matcher over the parameter space sampled from diallel.GS
## applies power.matcher over parameter draws from diallel.GS
par.cruncher.general <- function(par.mat, 
                                 n, 
                                 qtl.num = 1, 
                                 strains = c("AJ", "B6", "129", "NOD", "NZO", "CAST", "PWK", "WSB"),
                                 use.progress.bar = TRUE){
  power.list <- list()
  phenotype.list <- list()
  var.list <- list()
  if (use.progress.bar) {
    pb <- txtProgressBar(min=0, max=nrow(par.mat), style=3)
  }
  for (i in 1:nrow(par.mat)) {
    # Makes a list of cross powers - each element of list is a power matrix for a given set of parameters from diallel.GS
    power.list[[i]] <- power.matcher.general(par.vec = par.mat[i,], 
                                             n = n, 
                                             qtl.num = qtl.num, 
                                             strains = strains)
    eff.var.list <- var.matcher.general(par.vec = par.mat[i,], 
                                        qtl.num = qtl.num, 
                                        strains = strains)
    phenotype.list[[i]] <- eff.var.list[[1]]
    var.list[[i]] <- eff.var.list[[2]]
    if (use.progress.bar) {
      # Makes a progress bar
      setTxtProgressBar(pb, i)
    }
  }
  final <- list(power.list, phenotype.list, var.list)
  return(final)
}

## Moving from power (probabilities) to consequences
## Function to calculate the probability of the possible outcomes (consequences) of a QTL mapping experiment
utility.calculator.general <- function(power.mat, 
                                       col, 
                                       qtl.num = 1){
  c.matrix <- matrix(nrow = nrow(power.mat), ncol = qtl.num + 1)
  total <- qtl.num
  for (i in 0:total) {
    p <- log(choose(total, i)) + i*log(power.mat[,col] + 5e-324) + (total-i)*log(1 - power.mat[,col] + 5e-324)
    c.matrix[,i+1] <- exp(p)
    cross.names <- rownames(power.mat)
    rownames(c.matrix) <- cross.names
  }
  return(c.matrix)
}

## Runs all consequences for all 3 types of cross for each combination of lines
utility.runner.general <- function(power.mat, 
                                   qtl.num = 1){
  # Consequence probabilities
  c_bc1 <- utility.calculator.general(power.mat, 1, qtl.num)  
  c_rbc1_1 <- utility.calculator.general(power.mat, 2, qtl.num)
  c_rbc1_2 <- utility.calculator.general(power.mat, 3, qtl.num)
  c_bc2 <- utility.calculator.general(power.mat, 4, qtl.num)
  c_rbc2_1 <- utility.calculator.general(power.mat, 5, qtl.num)
  c_rbc2_2 <- utility.calculator.general(power.mat, 6, qtl.num)
  c_f2 <- utility.calculator.general(power.mat, 7, qtl.num)
  
  # Processing output
  c.list <- list(c_bc1, c_rbc1_1, c_rbc1_2, c_bc2, c_rbc2_1, c_rbc2_2, c_f2)
  return(c.list)
}

## Calculating expected utility given a parameter set
## Function calculates the expected utility (counts of QTL mapped: 2-0) for a given cross
eu.calculator <- function(c.mat){
  eu.vec <- 0
  for (i in 1:ncol(c.mat)) {
    eu.vec <- eu.vec + (i-1)*c.mat[,i]
  }
  return(eu.vec)
}

## Applies eu.calculator over all crosses for a given parameter set
eu.grinder.general <- function(c.list){
  # Expected utilities
  bc1 <- eu.calculator(c.list[[1]])
  rbc1_1 <- eu.calculator(c.list[[2]])
  rbc1_2 <- eu.calculator(c.list[[3]])
  bc2 <- eu.calculator(c.list[[4]])
  rbc2_1 <- eu.calculator(c.list[[5]])
  rbc2_2 <- eu.calculator(c.list[[6]])
  f2 <- eu.calculator(c.list[[7]])
  
  # Processing output
  eu.mat <- cbind(bc1, rbc1_1, rbc1_2, bc2, rbc2_1, rbc2_2, f2)
  colnames(eu.mat) <- c("bc1", "rbc1_1", "rbc1_2", "bc2", "rbc2_1", "rbc2_2", "f2")
  return(eu.mat)
}

## Selects specific element from matrix
picker <- function(mat, row, col){
  return(mat[row,col])
}

#' Propagate Gibbs Sampler (BayesDiallel model) to posterior utilities. 
#'
#' This function runs takes output from the BayesDiallel Gibbs Sampler and propagates these effects to the
#' utility space.
#' 
#' @param gibbs.object Posterior samples from the BayesDiallel model fit with diallel.gibbs().
#' @param n DEFAULT: 50. The number of individuals to be included in the proposed QTL mapping
#' experiment.
#' @param qtl.num DEFAULT: 1. The number of QTL underlying the estimated strain-level effects.
#' If greater than 1, the effects are reduced accordingly.
#' @param strains.relabel DEFAULT: NULL. Option to rename strains.
#' @param use.progress.bar DEFAULT: TRUE. A progress bar shows how calculating the utilities is progressing.
#' @export evaluate.experiments
#' @examples evaluate.experiments()
evaluate.experiments <- function(gibbs.object, 
                                 n = 50, 
                                 qtl.num = 1, 
                                 strains.relabel = NULL,
                                 use.progress.bar = TRUE){
  # Using previous functions
  par.mat <- gibbs.object$mcmc
  strains <- gibbs.object$strains
  num.strains <- length(strains)
  
  if (!is.null(strains.relabel)) {
    strains <- strains.relabel
  }
  
  par.bundle <- par.cruncher.general(par.mat = par.mat, 
                                     n = n, 
                                     qtl.num = qtl.num, 
                                     strains = strains,
                                     use.progress.bar = use.progress.bar)
  var.list <- par.bundle[[3]]
  pheno.list <- par.bundle[[2]]
  power.list <- par.bundle[[1]]
  u.list <- lapply(power.list, function(x) utility.runner.general(x, qtl.num = qtl.num))
  power.list <- lapply(u.list, function(x) eu.grinder.general(x))

  # Initializing output lists
  power.exp.list <- var.exp.list <- list(f2 = list(),
                                         bc1 = list(),
                                         bc2 = list(),
                                         rbc1_1 = list(),
                                         rbc1_2 = list(),
                                         rbc2_1 = list(),
                                         rbc2_2 = list())
  pheno.exp.list <- list(f2=list(hom1 = list(),
                                 hom2 = list(),
                                 het = list()),
                         bc1=list(hom = list(),
                                  het = list()),
                         bc2=list(hom = list(),
                                  het = list()),
                         rbc1_1=list(hom = list(),
                                     het = list()),
                         rbc1_2=list(hom = list(),
                                     het = list()),
                         rbc2_1=list(hom = list(),
                                     het = list()),
                         rbc2_2=list(hom = list(),
                                     het = list()))
  
  exp.type <- colnames(power.list[[1]])
  cross.type <- rownames(power.list[[1]])
  
  for (i in 1:length(exp.type)) {
    for (j in 1:choose(num.strains, 2)) {
      # utility
      power.experiment <- as.numeric(do.call(c, lapply(power.list, function(x) picker(x, j, i))))
      power.exp.list[[exp.type[i]]][[cross.type[j]]] <- power.experiment
      # variance
      var.experiment <- as.numeric(do.call(c, lapply(var.list, function(x) picker(x, j, i))))
      var.exp.list[[exp.type[i]]][[cross.type[j]]] <- var.experiment
    }
  }

  pheno.type <- colnames(pheno.list[[1]])
  for (i in 1:length(pheno.type)) {
    this.exp <- strsplit(pheno.type, split="-", fixed=TRUE)[[i]][1]
    this.pheno <- strsplit(pheno.type, split="-", fixed=TRUE)[[i]][2]
    for (j in 1:choose(num.strains, 2)) {
      # effect
      pheno.experiment <- as.numeric(do.call(c, lapply(pheno.list, function(x) picker(x, j, i))))
      pheno.exp.list[[this.exp]][[this.pheno]][[cross.type[j]]] <- pheno.experiment
    }
  }
  results <- list(power = power.exp.list, 
                  pheno = pheno.exp.list, 
                  var = var.exp.list, 
                  qtl.num = qtl.num, 
                  n = n,
                  strains = strains)
  return(results)
}
```
