## Supplementary material for "Bayesian decision theoretic design of two-founder experimental crosses given diallel data": diallel_GS.R

```
# Creates a group-identity matrix given a vector of factors
incidence.matrix <- function(fact) {
  m <- diag(nlevels(fact))[fact,]
  colnames(m) <- levels(fact)
  return(m)
}
# Creates constraint matrix
make.M <- function(X) {
  j <- ncol(X)
  k <- (-1 + sqrt(j))*(j - 1)^(-3/2)
  m <- 1/sqrt(j - 1)
  c <- (nrow(X) - 2)*k + m
  M <- diag(j - 1)
  M[M == 0] <- -k
  M <- rbind(M, rep(-m, j - 1))
  return(M)
}

# Update functions
update.beta <- function(X, 
                        y, 
                        res.var, 
                        prior.var) {
  post.var <- chol2inv(chol(t(X) %*% X + res.var*chol2inv(chol(prior.var))))*res.var
  post.mean <- (1/res.var)*post.var %*% t(X) %*% y
  beta.vec <- c(mnormt::rmnorm(1, mean=post.mean, varcov=post.var))
  return(beta.vec)
}
update.sigma.2 <- function(X, 
                           y, 
                           prior.alpha, 
                           prior.beta, 
                           par.vec) {
  post.alpha <- (nrow(X) + prior.alpha)/2
  post.beta <- (t(y - X %*% par.vec) %*% (y - X %*% par.vec) + prior.beta*prior.alpha)/2
  sigma.2 <- 1/rgamma(1, shape=post.alpha, rate=post.beta)
  return(sigma.2)
}
update.tau <- function(K, 
                       prior.alpha, 
                       prior.beta, 
                       par.vec) {
  post.alpha <- (ncol(K) + prior.alpha)/2
  post.beta <- (t(par.vec) %*% chol2inv(chol(K)) %*% par.vec + prior.beta)/2
  tau <- 1/rgamma(1, shape=post.alpha, rate=post.beta)
  return(tau)
}

#' Gibbs sampler that specifies a form of the BayesDiallel model. 
#'
#' This function runs the BayesDiallel Gibbs Sampler for diallel cross data.
#'
#' @param phenotype Vector of the phenotypes. Expects a quantitative phenotype.
#' @param sex Binary vector for females and males. By default, expects females to be coded as 1 and males as 0.
#' @param is.female DEFAULT: TRUE. If TRUE, then sex expects female=1 and male=0. If FALSE, then female=0 and male=1.
#' @param mother.str Vector of strain identities of mother/dam.
#' @param father.str Vector of strain identities of father/sire.
#' @param n.iter DEFAULT: 1000. The number of samples to be collected.
#' @param burn.in DEFAULT: 10000. The number of samples run as a burn-in, which are not stored.
#' @param multi.chain DEFAULT: 1. The number of MCMC chains to run. 
#' @param thin DEFAULT: 1. If 1, no thinning is used. If 2, then every other sample is stored. If 3, then every
#' third observation is stored. And so on.
#' @param sigma2.starter DEFAULT: 5. Starting value for the residual error variance parameter in the Gibbs Sampler.
#' @param tau_add.starter DEFAULT: 2. Starting value for the variance parameter of the additive effects in the Gibbs Sampler.
#' @param tau_inbred.starter DEFAULT: 2. Starting value for the variance parameter of the inbred effects in the Gibbs Sampler.
#' @param tau_mat.starter DEFAULT: 2. Starting value for the variance parameter of the maternal effects in the Gibbs Sampler.
#' @param tau_epi_sym.starter DEFAULT: 2. Starting value for the variance parameter of the symmetric epistatic effects in the Gibbs Sampler.
#' @param tau_epi_asym.starter DEFAULT: 2. Starting value for the variance parameter of the asymmetric epistatic effects in the Gibbs Sampler.
#' @param strains.reorder DEFAULT: c("AJ", "B6", "129", "NOD", "NZO", "CAST", "PWK", "WSB"). Orders the strains based on the standard
#' ordering of Collaborative Cross founders
#' @param strains.rename DEFAULT: c("AJ", "B6", "129", "NOD", "NZO", "CAST", "PWK", "WSB"). Renames the strains. By default, expects the
#' founders of the Collaborative Cross.
#' @param use.constraint DEFAULT: TRUE. Use a rotation matrix to reduce the dimension of strain matrices because the system is linearly dependent.
#' Can result in narrower posterior intervals.
#' @param use.progress.bar DEFAULT: TRUE. A progress bar shows how sampling is progressing.
#' @export diallel.gibbs
#' @examples diallel.gibbs()
diallel.gibbs <- function(phenotype, 
                          sex, 
                          is.female = TRUE, 
                          mother.str, 
                          father.str, 
                          n.iter = 1000, 
                          burn.in = 10000, 
                          multi.chain = 1, 
                          thin = 1,
                          sigma.2.starter = 5, 
                          tau_add.starter = 2, 
                          tau_inbred.starter = 2, 
                          tau_mat.starter = 2, 
                          tau_epi_sym.starter = 2, 
                          tau_epi_asym.starter = 2,
                          strains.reorder = c("AJ", "B6", "129", "NOD", "NZO", "CAST", "PWK", "WSB"),
                          strains.rename = c("AJ", "B6", "129", "NOD", "NZO", "CAST", "PWK", "WSB"),
                          use.constraint = TRUE,
                          use.progress.bar = TRUE) {
  if (!is.null(strains.reorder)){
    mother.str <- factor(mother.str, levels=strains.reorder)
    father.str <- factor(father.str, levels=strains.reorder)
  }
  else {
    mother.str <- factor(mother.str)
    father.str <- factor(father.str)
  }
  
  # Defining strain columns and incidence matrices
  if(!is.null(strains.rename)) {
    mother.str <- factor(mother.str, labels=strains.rename)
    father.str <- factor(father.str, labels=strains.rename)
  }
  
  strains <- unique(c(as.character(mother.str), as.character(father.str)))
  num.strains <- length(strains)
  
  mom.mat <- incidence.matrix(mother.str)
  pop.mat <- incidence.matrix(father.str)
  
  # For labeling
  strain.names <- colnames(mom.mat)
  
  # Labels
  add.names <- paste("add", strain.names, sep=":")
  dom.names <- paste("inbred", strain.names, sep=":")
  mat.names <- paste("mat", strain.names, sep=":")
  epi.names <- rep(NA, (length(strain.names)*(length(strain.names)-1))/2)
  counter <- 1
  for (m in 1:length(strain.names)) {
    for (n in 1:length(strain.names)) {
      if (m > n) {
        epi.names[counter] <- paste(strain.names[m], strain.names[n], sep=";")
        counter <- counter + 1
      }
    }
  }
  
  # Setting up the data
  y <- as.vector(phenotype)
  n <- length(y)
  
  # By default: female = 1, male = 0
  if(!is.female){
    sex <- ifelse(sex, 1, 0)
  } 
  
  # Make X
  X <- cbind(rep(1, n), sex, ifelse(mother.str == father.str, 1, 0))
  
  # Setting up Z
  # Additive
  add.part <- mom.mat + pop.mat
  # Inbred
  inbred.part <- 1*(add.part == 2)
  # Maternal
  mat.part <- mom.mat - pop.mat
  # Epistatic
  mom.index <- apply(mom.mat, 1, function(x) which(x == 1))
  pop.index <- apply(pop.mat, 1, function(x) which(x == 1))
  combined.index <- cbind(mom.index, pop.index)
  sorted.index <- t(apply(combined.index, 1, function(x) sort(x, decreasing=TRUE)))
  symmetric.pair <- paste(strain.names[sorted.index[,1]], strain.names[sorted.index[,2]], sep=";")
  epi_sym.part <- 1*(t(apply(matrix(symmetric.pair, nrow=length(symmetric.pair), ncol=1), 1, function(x) x == epi.names)))
  epi_asym.part <- epi_sym.part
  reciprocals <- apply(sorted.index, 1, function(x) paste(x, collapse=" ")) != apply(combined.index, 1, function(x) paste(x, collapse=" "))
  epi_asym.part[reciprocals,] <- -1*epi_sym.part[reciprocals,]
  
  # Making constraint matrix
  if(use.constraint){
    M.add <- make.M(X=add.part)
    M.inbred <- make.M(X=inbred.part)
    M.mat <- make.M(X=mat.part)
    M.epi_sym <- make.M(X=epi_sym.part)
    M.epi_asym <- make.M(X=epi_asym.part)
    
    # Transforming design matrices
    add.part <- add.part %*% M.add
    inbred.part <- inbred.part %*% M.inbred
    mat.part <- mat.part %*% M.mat
    epi_sym.part <- epi_sym.part %*% M.epi_sym
    epi_asym.part <- epi_asym.part %*% M.epi_asym
  }
  
  # Overall design matrix
  X.all <- cbind(X, add.part, inbred.part, mat.part, epi_sym.part, epi_asym.part)
  if (burn.in > 0) {
    cat("Burn-in:\n")
    # Progress bar
    if (use.progress.bar) {
      burnin.pb <- txtProgressBar(min=0, max=burn.in, style=3)
    }
  }
  # For multiple chains
  if (multi.chain > 1) {
    chain.list <- list()
  } 
  for (j in 1:multi.chain) {
    # Allocate memory
    n.col <- 3 + 3*num.strains + 2*choose(num.strains, 2) + 6
    p.mat <- matrix(0, n.iter, n.col) # 3 fixed effects, 8 additive, 8 inbred, 8 maternal, 28 epistatic symmetric, 28 epistatic asymmetric
    
    # Set hyperparameters
    ga.alpha <- 0.002
    ga.beta <- 2
    
    # Initialization
    fix.vec <- c(mean(phenotype[sex==0]), mean(phenotype[sex==1]), 0)
    rand.vec <- rnorm(ncol(X.all) - 3, 0, 20)
    beta.vec <- c(fix.vec, rand.vec)
    
    sigma.2 <- sigma.2.starter
    tau_add <- tau_add.starter
    tau_inbred <- tau_inbred.starter
    tau_mat <- tau_mat.starter
    tau_epi_sym <- tau_epi_sym.starter
    tau_epi_asym <- tau_epi_sym.starter
    
    # Keep track of matrix rows
    counter <- burnin.counter <- 1
    
    # Updating parameters
    for(i in 1:((n.iter*thin)+burn.in)){
      # prior covariance matrix
      Sigma0 <- diag(c(rep(1000, 3), 
                       rep(tau_add, ncol(add.part)),
                       rep(tau_inbred, ncol(inbred.part)), 
                       rep(tau_mat, ncol(mat.part)),
                       rep(tau_epi_sym, ncol(epi_sym.part)), 
                       rep(tau_epi_asym, ncol(epi_asym.part))))
      # constraint <- matrix(c(c(1,1,1,1,0,0,0,1),
      #                        c(1,1,1,1,0,0,0,1),
      #                        c(1,1,1,1,0,0,0,1),
      #                        c(1,1,1,1,0,0,0,1),
      #                        c(0,0,0,0,1,0,1,0),
      #                        c(0,0,0,0,0,1,0,0),
      #                        c(0,0,0,0,1,0,1,0),
      #                        c(1,1,1,1,0,0,0,1)),
      #                      byrow=TRUE, nrow=8, ncol=8)*tau_add
      # Sigma0[4:11,4:11] <- constraint + diag(8)*0.01
      
      
      
      # Update beta.vec
      beta.vec <- update.beta(X.all, y, sigma.2, Sigma0)
      
      # Update sigma.2
      sigma.2 <- update.sigma.2(X.all, y, ga.alpha, ga.beta, beta.vec)
      
      # Updating tau_add
      a.index <- 4:(4 + ncol(add.part) - 1)
      tau_add <- update.tau(diag(ncol(add.part)), ga.alpha, ga.beta, beta.vec[a.index])
      
      # Updating tau_inbred
      i.index <- (4 + ncol(add.part)):(4 + ncol(add.part) + ncol(inbred.part) - 1)
      tau_inbred <- update.tau(diag(ncol(inbred.part)), ga.alpha, ga.beta, beta.vec[i.index])
      
      # Updating tau_mat
      m.index <- (4 + ncol(add.part) + ncol(inbred.part)):(4 + ncol(add.part) + ncol(inbred.part) + ncol(mat.part) - 1)
      tau_mat <- update.tau(diag(ncol(mat.part)), ga.alpha, ga.beta, beta.vec[m.index])
      
      # Updating tau_epi_sym
      e_sym.index <- (4 + ncol(add.part) + ncol(inbred.part) + ncol(mat.part)):(4 + ncol(add.part) + ncol(inbred.part) + ncol(mat.part) + ncol(epi_sym.part) - 1)
      tau_epi_sym <- update.tau(diag(ncol(epi_sym.part)), ga.alpha, ga.beta, beta.vec[e_sym.index])
      
      # Updating tau_epi_asym
      e_asym.index <- (4 + ncol(add.part) + ncol(inbred.part) + ncol(mat.part) + ncol(epi_sym.part)):(4 + ncol(add.part) + ncol(inbred.part) + ncol(mat.part) + ncol(epi_sym.part) + ncol(epi_asym.part) - 1)
      tau_epi_asym <- update.tau(diag(ncol(epi_asym.part)), ga.alpha, ga.beta, beta.vec[e_asym.index])
      
      if (i <= burn.in) {
        # Progresses burn-in progress bar
        burnin.counter <- burnin.counter + 1
        if (use.progress.bar) {
          setTxtProgressBar(burnin.pb, burnin.counter)
        }
      }
      if (i == burn.in) {
        cat("\n")
      }
      # Keep values after burn-in
      if (i > burn.in) {
        if (i == burn.in + 1) { 
          cat("Saved MCMC sampling:\n")
          if (use.progress.bar) {
            pb <- txtProgressBar(min=0, max=n.iter*multi.chain, style=3)
          }
        }
        if ((i-1-burn.in) %% thin == 0) {
          if (use.constraint) {
            p.mat[counter,] <- c(beta.vec[1:3], 
                                 M.add %*% beta.vec[a.index],
                                 M.inbred %*% beta.vec[i.index],
                                 M.mat %*% beta.vec[m.index],
                                 M.epi_sym %*% beta.vec[e_sym.index],
                                 M.epi_asym %*% beta.vec[e_asym.index],
                                 sigma.2, tau_add, tau_inbred, tau_mat, tau_epi_sym, tau_epi_asym)
          }
          else {
            p.mat[counter,] <- c(beta.vec, sigma.2, tau_add, tau_inbred, tau_mat, tau_epi_sym, tau_epi_asym)
          }
          counter <- counter + 1
          if(use.progress.bar) {
            # Progresses progress bar
            setTxtProgressBar(pb, counter)
          }
        }
      }
    }
    colnames(p.mat) <- c("mu", "female", "inbred penalty", 
                         add.names, dom.names, mat.names, paste("epi_sym", epi.names, sep=":"), paste("epi_asym", epi.names, sep=":"),
                         "sigma2", "tau2 add", "tau2 inbred", "tau2 mat", "tau2 epi_sym", "tau2 epi_asym")
    
    # If multiple chains, build up list
    if(multi.chain > 1){
      chain.list[[j]] <- coda::as.mcmc(p.mat)
    } 
  }
  
  # Return matrix or list of matrix
  if (multi.chain > 1) {
    return(list(mcmc=coda::as.mcmc.list(chain.list),
                strains=levels(mother.str)))
  }
  else {
    return(list(mcmc=coda::as.mcmc(p.mat),
                strains=levels(mother.str)))
  }
}
```
