## Supplementary material for "Bayesian decision theoretic design of two-founder experimental crosses given diallel data": diallel_sim.R

```
#' Simulate diallel data from the BayesDiallel model
#'
#' This function generates data from the BayesDiallel model. It does not currently allow for sex
#' interactions with strain-level effects, as is possible in the proper BayesDiallel package.
#'
#' @param M DEFAULT: 8. The number of strains in the diallel cross. Default of 8 is the number of founders in the Collaborative Cross.
#' @param mu DEFAULT: 10. The grand mean of the phenotype.
#' @param inbred.mu DEFAULT: -5. The overall inbred effect.
#' @param female.mu DEFAULT: -2. The overall female effect (male is reference).
#' @param add.size The proportion of the variance explained by the additive effects.
#' @param inbred.size The proportion of the variance explained by the inbred effects.
#' @param epi.sym.size The proportion of the variance explained by the symmetric epistatic effects.
#' @param epi.asym.size The proportion of the variance explained by the asymmetric epistatic effects.
#' @param maternal.size The proportion of the variance explained by the maternal effects.
#' @param n.each DEFAULT: 5. The number of individuals simulated per cell of the diallel. 
#' Total sample size of diallel cross will be n.each * 64
#' @param num.sim DEFAULT: 1. The number of diallel cross simulations to perform.
#' @param strains DEFAULT: c("A", "B", "C", "D", "E", "F", "G", "H"). The strain names used. Defaults to 
#' simple letter representation.
#' @export simulate.diallel
#' @examples simulate.diallel()
simulate.diallel <- function(M = 8,
                             mu = 10, 
                             inbred.mu = -5, 
                             female.mu = -2,
                             add.size, 
                             inbred.size, 
                             epi.sym.size, 
                             epi.asym.size, 
                             maternal.size = 0, 
                             add.effect = NULL, 
                             inbred.effect = NULL, 
                             epi.sym.effect = NULL, 
                             epi.asym.effect = NULL, 
                             maternal.effect = NULL,
                             n.each = 5, 
                             num.sim = 1, 
                             strains = LETTERS[1:M]){
  
  non.sample.var <- function(x) {
    var.x <- var(x)*((length(x) - 1)/length(x))
    return(var.x)
  }
  
  ## Simulate diallel effects
  # add
  if (is.null(add.effect)) { add.effect <- rnorm(n = M) }
  add.effect <- (add.effect - mean(add.effect))/sqrt(non.sample.var(add.effect))
  add.effect <- 0.5*add.effect*sqrt(add.size)
  names(add.effect) <- strains
  # inbred
  if (is.null(inbred.effect)) { inbred.effect <- rnorm(n=M) }
  inbred.effect <- (inbred.effect - mean(inbred.effect))/sqrt(non.sample.var(inbred.effect))
  inbred.effect <- inbred.effect*sqrt(inbred.size)
  names(inbred.effect) <- strains
  # symmetric epistatic
  if (is.null(epi.sym.effect)) { epi.sym.effect <- rnorm(n=choose(M, 2)) }
  epi.sym.effect <- (epi.sym.effect - mean(epi.sym.effect))/sqrt(non.sample.var(epi.sym.effect))
  epi.sym.effect <- epi.sym.effect*sqrt(epi.sym.size)
  names(epi.sym.effect) <- make.pairs(strains)
  # asymmetric epistatic
  if (is.null(epi.asym.effect)) { epi.asym.effect <- rnorm(n=choose(M, 2)) }
  epi.asym.effect <- (epi.asym.effect - mean(epi.asym.effect))/sqrt(non.sample.var(epi.asym.effect))
  epi.asym.effect <- epi.asym.effect*sqrt(epi.asym.size)
  names(epi.asym.effect) <- make.pairs(strains)
  # maternal
  if (is.null(maternal.effect)) { maternal.effect <- rnorm(n=M) }
  maternal.effect <- (maternal.effect - mean(maternal.effect))/sqrt(non.sample.var(maternal.effect))
  maternal.effect <- 0.5*maternal.effect*sqrt(maternal.size)
  names(maternal.effect) <- strains
  
  ########################### Making component design matrices
  init.matrix <- diag(M); rownames(init.matrix) <- strains
  
  all.individuals <- make.all.pairs.matrix(strains)[sort(rep(1:length(strains)^2, times=n.each)),]; colnames(all.individuals) <- c("dam.id", "sire.id")
  dam.matrix <- init.matrix[all.individuals[, "dam.id"],]
  sire.matrix <- init.matrix[all.individuals[, "sire.id"],]
  
  ###### Additive
  add.matrix <- dam.matrix + sire.matrix; colnames(add.matrix) <- strains
  
  ###### Maternal
  maternal.matrix <- dam.matrix - sire.matrix; colnames(maternal.matrix) <- strains
  
  ###### Inbred
  inbred.matrix <- add.matrix - 1
  inbred.matrix[inbred.matrix == -1] <- 0
  inbred.fixed.effect <- rowSums(inbred.matrix); colnames(inbred.matrix) <- strains
  
  epi.init.matrix <- diag(choose(M, 2)); rownames(epi.init.matrix) <- make.pairs(strains)
  inbred.portion.epi.init.matrix <- matrix(0, nrow=M, ncol=choose(M, 2)); rownames(inbred.portion.epi.init.matrix) <- paste(strains, strains, sep=".")
  epi.init.matrix <- rbind(inbred.portion.epi.init.matrix, epi.init.matrix)
  
  ###### Epistatic
  epi.sym.matrix <- epi.init.matrix[apply(all.individuals, 1, function(x) paste(sort(x), collapse=".")),]; colnames(epi.sym.matrix) <- make.pairs(strains)

  asym.flip <- !apply(all.individuals, 1, function(x) paste(x, collapse=".")) == apply(all.individuals, 1, function(x) paste(sort(x), collapse="."))
  epi.asym.matrix <- epi.sym.matrix
  epi.asym.matrix[asym.flip,] <- -1*epi.sym.matrix[asym.flip,]
  
  ###### Sex
  is.female <- rbinom(n = n.each*M^2, size = 1, prob = 0.5)
  
  ###### Simulation
  noise.var <- 1 - add.size - inbred.size - epi.sym.size - epi.asym.size - maternal.size
  y.pred <- mu + is.female*female.mu + add.matrix %*% add.effect + inbred.fixed.effect*inbred.mu + inbred.matrix %*% inbred.effect + maternal.matrix %*% maternal.effect + epi.sym.matrix %*% epi.sym.effect + epi.asym.matrix %*% epi.asym.effect
  
  sample.scaled.resid <- function(n, 
                                  noise.var) {
    resid <- rnorm(n)
    resid <- (resid - mean(resid))/sqrt(non.sample.var(resid))
    resid <- resid*sqrt(noise.var)
    return(resid)
  }
  
  y <- sapply(1:num.sim, function(x) y.pred + sample.scaled.resid(n = length(y.pred), noise.var = noise.var))
  colnames(y) <- paste0("y.sim.", 1:num.sim)
  
  diallel.data <- data.frame(y, is.female, dam.id = all.individuals[,"dam.id"], sire.id = all.individuals[,"sire.id"])
  
  didact.input <- coda::as.mcmc(matrix(c(mu, female.mu, inbred.mu, 
                                         add.effect, 
                                         inbred.effect, 
                                         maternal.effect, 
                                         epi.sym.effect,
                                         epi.asym.effect,
                                         1 - add.size - inbred.size - maternal.size - epi.sym.size - epi.asym.size,
                                         add.size, inbred.size, maternal.size, epi.sym.size, epi.asym.size), nrow=1))
  colnames(didact.input) <- c("mu", "female", "inbred penalty", 
                              paste("add",names(add.effect), sep=":"), 
                              paste("inbred", names(inbred.effect), sep=":"),
                              paste("mat", names(maternal.effect), sep=":"),
                              paste("epi_sym", unlist(lapply(strsplit(x=names(epi.sym.effect), split=".", fixed=TRUE), function(x) paste(rev(x), collapse=";"))), sep=":"),
                              paste("epi_asym", unlist(lapply(strsplit(x=names(epi.asym.effect), split=".", fixed=TRUE), function(x) paste(rev(x), collapse=";"))), sep=":"),
                              "sigma2", "tau_add", "tau_inbred", "tau_mat", "tau_epi_sym", 'tau_epi_asym')
  results <- list(diallel.data = diallel.data,
                  effects = list(add = add.effect,
                               inbred = inbred.effect,
                               maternal = maternal.effect,
                               epi_sym = epi.sym.effect,
                               epi_asym = epi.asym.effect,
                               mu = mu,
                               inbred.mu = inbred.mu,
                               female.mu = female.mu),
                  didact.input = list(mcmc = didact.input,
                                    strains = names(add.effect)))
  return(results)
}

make.pairs <- function(strains, 
                       this.sep = "."){
  pairs <- rep(NA, choose(length(strains), 2))
  counter <- 1
  for (i in 1:(length(strains) - 1)) {
    for (j in (i+1):length(strains)) {
      pairs[counter] <- paste(strains[i], strains[j], sep = this.sep)
      counter <- counter + 1
    }
  }
  return(pairs)
}

make.all.pairs.matrix <- function(strains){
  pair.matrix <- matrix(NA, nrow = length(strains)^2, ncol=2)
  current.row <- 1
  for(i in 1:(length(strains))){
    for(j in 1:(length(strains))){
      pair.matrix[current.row,] <- c(strains[i], strains[j])
      current.row <- current.row + 1
    }
  }
  return(pair.matrix)
}
```
