## Supplementary material for "Bayesian decision theoretic design of two-founder experimental crosses given diallel data": plot_functions.R

```
#' Caterpillar plot of the highest posterior density (HPD) intervals for parameters from BayesDiallel model.
#' 
#' This function produces a caterpillar plot for samples from BayesDiallel posterior distributions.
#' 
#' @param gibbs.object Posterior samples from the BayesDiallel model fit with diallel.gibbs().
#' @param override.title DEFAULT: NULL. Allows title to be manually specified.
#' @param override.xlab DEFAULT: NULL. Allows x-axis label to be manually specified.
#' @param include.effect.types DEFAULT: c("mu", "female", "add", "mat", "inbred", "epi_sym", "epi_asym", "var"). 
#' The effect types to be included in the plot.
#' @param col DEFAULT: c("black", "black", "#A6CEE3", "#B2DF8A", "#FDBF6F", "#CAB2D6", "#D2B48C", "black").
#' Colors corresponding to the effect types.
#' @param inbred.penalty.col DEFAULT: "#FB9A99". The color for the overall inbred penalty.
#' @param rev.strain.output DEFAULT: TRUE. Reverses the ordering of strain effects.
#' @param manual.limits DEFAULT: NULL. Allows the x-axis limits to be manually specified.
#' @param override.col DEFAULT: NULL. Allows the HPD interval colors to be manually specified.
#' @param include.grid DEFAULT: TRUE. Adds gray horizontal guide lines for strain-level effects.
#' @param include.prefix DEFAULT: TRUE. If TRUE, effect type included on y-axis label as a prefix.
#' @param zero.lty DEFAULT: 2. The line type of the vertical line at 0.
#' @param zero.color DEFAULT: "gray". The color of the vertical line at 0.
#' @param effect.label.cex DEFAULT: 0.5. Specifies the size of the effect labels.
#' @export caterpillar.plot
#' @examples caterpillar.plot()
caterpillar.plot <- function(gibbs.object, 
                             override.title = NULL,
                             override.xlab = NULL,
                             include.effect.types = c("mu", "female", "add", "mat", "inbred", "epi_sym", "epi_asym", "var"),
                             col = c("black", "black", "#A6CEE3", "#B2DF8A", "#FDBF6F", "#CAB2D6", "#D2B48C", "black"),
                             inbred.penalty.col = "#FB9A99",
                             rev.strain.output = TRUE,
                             manual.limits = NULL,
                             override.col = NULL,
                             include.grid = TRUE,
                             include.prefix = TRUE,
                             zero.lty = 2,
                             zero.color = "gray",
                             effect.label.cex = 0.5){
  mcmc.object <- gibbs.object$mcmc
  
  # Processing the plotted variables
  reorder.names <- NULL
  include.col <- rep(FALSE, ncol(mcmc.object))
  use.color <- NULL
  for (i in 1:length(include.effect.types)) {
    if (include.effect.types[i] == "mu") {
      temp.include.col <- grepl(x=colnames(mcmc.object), pattern="^mu$", perl=TRUE)
    }
    else if (include.effect.types[i] == "var") {
      temp.include.col <- grepl(x=colnames(mcmc.object), pattern="tau|sigma")
    }
    else {
      temp.include.col <- grepl(x=colnames(mcmc.object), pattern=include.effect.types[i]) & !grepl(x=colnames(mcmc.object), pattern="tau|sigma")
    }
    temp.include.col.int <- which(temp.include.col)
    if (rev.strain.output) {
      temp.include.col.int <- rev(temp.include.col.int)
    }
    if (include.effect.types[i] == "inbred") {
      use.color <- c(use.color, c(rep(col[i], sum(temp.include.col) - 1), inbred.penalty.col))
    }
    else {
      use.color <- c(use.color, rep(col[i], sum(temp.include.col)))
    }
    reorder.names <- c(reorder.names, colnames(mcmc.object)[temp.include.col.int])
  }
  
  if (!is.null(override.col)) {
    use.color <- override.col
  }
  
  mcmc.object <- mcmc.object[,reorder.names]
  num.var <- ncol(mcmc.object)
  ci95.data <- coda::HPDinterval(mcmc.object, prob=0.95)
  ci50.data <- coda::HPDinterval(mcmc.object, prob=0.50)
  par.names <- rownames(ci95.data)
  means.data <- as.vector(apply(mcmc.object, 2, function(x) mean(x)))
  median.data <- as.vector(apply(mcmc.object, 2, function(x) median(x)))
  
  if (!is.null(override.title)) {
    title <- override.title
  }
  else {
    titlename <- strsplit(deparse(substitute(gibbs.object)), ".", fixed = TRUE)[[1]][1]
    title <- paste(titlename, "parameters")
  }
  
  # Window limits
  if (!is.null(manual.limits)) {
    this.x.lim <- manual.limits
  }
  else {
    this.x.lim <- c(min(ci95.data, na.rm = TRUE), max(ci95.data, na.rm = TRUE))
  }
  
  this.xlab <- ifelse(is.null(override.xlab), "HPD intervals of strain effects and model parameters", override.xlab)
  plot(ci95.data[1, 1:2], 
       c(1,1), 
       panel.first=abline(h = 1, lty = 3, col = ifelse(include.grid, "gray88", "white")), 
       type = "l", 
       ylim = c(0, num.var + 1), 
       main = title, 
       xlim = this.x.lim, xlab = this.xlab, 
       ylab = "", yaxt = "n", lty = 1, lwd = 1, col = use.color[1], frame.plot=FALSE)
  if (length(ci95.data[1,]) > 2) {
    for (i in seq(3, length(ci95.data[1,]) - 1, by = 2)) {
      lines(ci95.data[1, c(i,i + 1)], c(1, 1), lty = 1, lwd = 1, col = use.color[1], lend = 2)
    }
  }
  lines(ci50.data[1, 1:2], c(1, 1), lty = 1, lwd = 3, col = use.color[1], lend = 2)
  if (length(ci50.data[1,]) > 2){
    for (i in seq(3, length(ci50.data[1,]) - 1, by = 2)) {
      lines(ci50.data[1, c(i,i+1)], c(1,1), lty = 1, lwd = 3, col = use.color[1], lend = 2)
    }
  }
  for (j in 2:num.var) {
    if (include.grid) { abline(h = j, lty = 3, col = "gray88") }
    for (k in seq(1, length(ci95.data[j,]) - 1, by = 2)) {
      lines(ci95.data[j, c(k,k+1)], c(j,j), lty = 1, lwd = 1, col = use.color[j], lend = 2)
    }
    for (k in seq(1, length(ci50.data[j,]) - 1, by = 2)) {
      lines(ci50.data[j, c(k,k+1)], c(j,j), lty = 1, lwd = 3, col = use.color[j], lend = 2)
    }
  }
  points(median.data, 1:num.var, pch = "l", col = "white")
  points(means.data, 1:num.var, pch = "l", col = use.color)
  abline(v = 0, lty = zero.lty, col = zero.color)
  ## Remove effect types
  if (!include.prefix) { par.names <- gsub(pattern = "add:|mat:|inbred:|epi_sym:|epi_asym:",
                                           replacement = "",
                                           x = par.names) }
  axis(2, c(1:num.var), par.names, c(1:num.var), 
       line = -1.5, 
       las = 2, 
       tck = -0.005, 
       cex.axis = effect.label.cex)
}

#' Heatmap of mean phenotype for diallel cross.
#' 
#' This function produces a heatmap of the mean phenotypes per diallel cell, providing a visualization of
#' raw diallel data.
#' 
#' @param mother.str.var Name of variable encoding the mother/dam strain identity.
#' @param father.str.var Name of variable encoding the father/sire strain identity.
#' @param phenotype Name of the phenotype variable. A quantitative phenotype is expected.
#' @param data The data.frame that contains the phenotype and parental strain identities.
#' @param phenotype.title DEFAULT: NULL. The title for the phenotype. If NULL, it is left blank.
#' @param do.reorder DEFAULT: TRUE. If TRUE, reorders strains according to strain.reorder argument.
#' @param strains.reorder DEFAULT: c("AJ", "B6", "129", "NOD", "NZO", "CAST", "PWK", "WSB").
#' By default, reorders strains based on the standard order of the Collaborative Cross strains.
#' @param strain.names DEFAULT: c("AJ", "B6", "129", "NOD", "NZO", "CAST", "PWK", "WSB").
#' By default, renames strains to the slightly abbreviated Collaborative Cross labels.
#' @param strain.colors DEFAULT: c("#F0F000", "#808080", "#F08080", "#1010F0", "#00A0F0", "#00A000", "#F00000", "#9000E0").
#' By default, the standard Collaborative Cross colors are specified.
#' @param strain.cex DEFAULT: 1. Specifies the size of strain labels.
#' @export diallel.phenotype.map
#' @examples diallel.phenotype.map
diallel.phenotype.map <- function(mother.str.var, 
                                  father.str.var, 
                                  phenotype, 
                                  data,
                                  phenotype.title = NULL,
                                  do.reorder = TRUE,
                                  strains.reorder = c("AJ", "B6", "129", "NOD", "NZO", "CAST", "PWK", "WSB"),
                                  strain.names = c("AJ", "B6", "129", "NOD", "NZO", "CAST", "PWK", "WSB"),
                                  strain.colors = c("#F0F000", "#808080", "#F08080", "#1010F0", 
                                                    "#00A0F0", "#00A000", "#F00000", "#9000E0"),
                                  strain.cex = 1){
  if (do.reorder) {
    data[,mother.str.var] <- factor(data[,mother.str.var], levels=strains.reorder)
    data[,father.str.var] <- factor(data[,father.str.var], levels=strains.reorder)
  }

  num.strains <- length(unique(c(data[, mother.str.var], data[, father.str.var])))
  data.mat <- matrix(NA, nrow=num.strains, ncol=num.strains)
  match.vec <- expand.grid(levels(data[, mother.str.var]), levels(data[, mother.str.var]))
  match.vec$paste <- paste(match.vec$Var1, match.vec$Var2, sep = "x")
  match.mat <- matrix(as.character(match.vec$paste), nrow=num.strains, ncol=num.strains)
  colnames(match.mat) <- rownames(match.mat) <- strain.names
  colnames(data.mat) <- rownames(data.mat) <- strain.names
  my_palette <- colorRampPalette(c("white", "black"))(n = 299)
  
  data$cross <- paste(data[,mother.str.var], data[,father.str.var], sep="x")
  
  phenotype.summary <- aggregate(formula(paste(phenotype, "cross", sep="~")), data=data, FUN=mean, na.rm=TRUE)
  for (j in 1:num.strains) {
    for (k in 1:num.strains) {
      this.cross <- as.character(match.mat[j, k])
      try(data.mat[j, k] <- as.numeric(phenotype.summary[phenotype.summary$cross == this.cross, phenotype]),
          silent = TRUE)
    }
  }
  if (is.null(phenotype.title)) { phenotype.title <- phenotype }
  heatmap(t(data.mat), 
          scale = "none", 
          Colv = NA, 
          Rowv = NA,
          revC = TRUE, 
          symm = TRUE, 
          ColSideColors = strain.colors, 
          margins = c(6, 6),
          RowSideColors = strain.colors, 
          xlab = expression(bold("Paternal")), 
          ylab = expression(bold("Maternal")), 
          col = my_palette, 
          main = paste("Observed mean", phenotype.title),
          cexRow = strain.cex,
          cexCol = strain.cex)
}

#' Phenotype ramp plot for diallel cross.
#' 
#' This function produces a phenotype ramp to match the heatmap from diallel.phenotype.map().
#' 
#' @param mother.str.var Name of variable encoding the mother/dam strain identity.
#' @param father.str.var Name of variable encoding the father/sire strain identity.
#' @param phenotype Name of the phenotype variable. A quantitative phenotype is expected.
#' @param data The data.frame that contains the phenotype and parental strain identities.
#' @param include.decimal DEFAULT: FALSE. If FALSE, the scale labels are rounded to whole numbers.
#' @export diallel.phenotype.scale
#' @examples diallel.phenotype.scale()
diallel.phenotype.scale <- function(mother.str.var, 
                                    father.str.var, 
                                    phenotype, 
                                    data,
                                    include.decimal = FALSE){
  my_palette <- colorRampPalette(c("white", "black"))(n = 299)
  
  data$cross <- paste(data[,mother.str.var], data[,father.str.var], sep = "x")
  phenotype.summary <- aggregate(formula(paste(phenotype, "cross", sep = "~")), 
                                 data = data, 
                                 FUN = mean, 
                                 na.rm = TRUE)
  
  plot(c(1:length(my_palette)), rep(1, length(my_palette)), 
       pch = "|", col = my_palette, cex = 3, ylab = "", xlab = "",
       yaxt = "n", frame.plot = FALSE, ylim = c(0.75, 1.25), xaxt = "n",
       xlim=c(0, 350))
  if (!include.decimal) {
    labs <- signif(as.numeric(pretty(range(phenotype.summary[,phenotype], na.rm = TRUE))), digits = 2)
  }
  else {
    labs <- round(as.numeric(pretty(range(phenotype.summary[,phenotype], na.rm = TRUE))), digits = 1)
  }
  axis(side=1, 
       at = seq(from = 1, 
                to = length(my_palette), 
                length.out = length(labs)),
       labels = labs)
}


#' DIDACT grid plot of posterior utility.
#' 
#' This function takes posterior samples of utilities from evaluate.experiments() and produces a grid plot
#' of the posterior utilities for specified cross type.
#' 
#' @param results Samples of posterior utility produced by evaluate.experiments().
#' @param utility.type DEFAULT: "power". The posterior utility to be plotted. Currently the options are "power"
#' and "contrasts". Power is more appropriate for Mendelian-like phenotypes. Contrasts make less assumptions.
#' @param cross.type DEFAULT: "f2". Current options include "f2", "bc", "rbc1", and "rbc2".
#' @param pheno.name DEFAULT: "". Included in the information panel.
#' @param col.range DEFAULT: c("white", "black"). If specified, will create a color spectrum scale between the
#' two colors included.
#' @param col.spectrum DEFAULT: "blue2red". Use pre-specified spectrum. Options include "blue2red", "gray", 
#' "green2red", and "blue2green".
#' @param path DEFAULT: NULL. If a path is specified, a pdf will be created.
#' @param height DEFAULT: 12. The height of the pdf in inches.
#' @param width DEFAULT: 12. The width of the pdf in inches.
#' @param strains.relabel DEFAULT: NULL. Option to re-label the strains.
#' @param include.off.x DEFAULT: TRUE. If TRUE, "X" is plotted on the diagonal and lower diagonal of the grids.
#' If FALSE, square is left empty.
#' @param include.biparent.labels DEFAULT: TRUE. If TRUE, arrows are included to signify backcross parent.
#' @param include.density DEFAULT: TRUE. If TRUE, histogram of posterior density is included in square.
#' @param include.var.pie DEFAULT: FALSE. If TRUE, pie chart of phenotypic variance is included in square.
#' @param include.bar.plots DEFAULT: TRUE. If TRUE, box plots of posterior genotype class phenotypes are 
#' included in square.
#' @param density.col DEFAULT: "white". The color of the histogram if included in square.
#' @param border.col DEFAULT: "black". The color of the border lines of the various figures.
#' @param median.line.col DEFAULT: "black". The color of the vertical line that marks the median utility.
#' @param absolute.density.scale DEFAULT: TRUE. If TRUE, the densities are scaled to the same height.
#' @param include.info.plot DEFAULT: TRUE. If TRUE, info plot is included in one of the diagonal squares.
#' @param include.rank DEFAULT: FALSE. If TRUE, the rank of mean posterior utility is included on the cross 
#' square.
#' @param rank.col DEFAULT: "red". The color of the rank index included on the square.
#' @param label.cex DEFAULT: 1.1. Specifies the size of the strain labels.
#' @param label.padj DEFAULT: -0.3. Specifies how much the strain label is shifted from the square.
#' @export diallelPlotter
#' @examples diallelPlotter()
diallelPlotter <- function(results, 
                           utility.type = c("power", "contrasts"),
                           cross.type = c("f2", "bc", "rbc1", "rbc2"), 
                           pheno.name = "", 
                           col.range = c("white", "black"),
                           col.spectrum = c("blue2red", "gray", "green2red", "blue2green"),
                           path = NULL,
                           height = 12,
                           width = 12,
                           strains.relabel = NULL,
                           include.off.x = TRUE,
                           include.biparent.labels = TRUE,
                           include.density = TRUE,
                           include.var.pie = FALSE,
                           include.bar.plots = TRUE,
                           density.col = "white",
                           border.col = "black",
                           median.line.col = "black",
                           absolute.density.scale = TRUE,
                           include.info.plot = TRUE,
                           include.rank = FALSE,
                           rank = NULL,
                           rank.col = "red",
                           label.cex = 1.1,
                           label.padj = -0.3,
                           ...){
  cross.type <- cross.type[1]
  utility.type <- utility.type[1]
  
  if (utility.type == "power") {
    utility.list <- results$power
  }
  else {
    utility.list <- results$var
  }
  rank.list <- calc.ranks(utility.list = utility.list, 
                          cross.type = cross.type)

  pheno.list <- results$pheno
  var.list <- results$var
  qtl.num <- results$qtl.num
  n <- results$n
  col.spectrum <- col.spectrum[1]
  strains <- results$strains
  num.strains <- length(strains)
  if (utility.type == "power") {
    x.high <- qtl.num
  }
  else if (utility.type == "contrasts") {
    #x.high <- max(sapply(1:length(results$var), function(y) max(sapply(1:length(results$var[[y]]), function(x) max(results$var[[y]][[x]])))))
    x.high <- 100
  }

  absolute.max <- NULL
  if (absolute.density.scale) {
    for (i in grep(x=names(utility.list), pattern=cross.type, value=TRUE)) {
      #x.high <- ifelse(utility.type == "power", qtl.num, 100)
      absolute.max <- max(absolute.max,
                          1.05*max(unlist(lapply(utility.list[[i]], function(x) max(hist(x,
                                                                                         plot = FALSE, 
                                                                                         breaks = seq(0, x.high, length.out=20))$density)))))
    }
  }
  
  ## Setting color spectrum
  spectrum <- make.spectrum(col.range=col.range, col.spectrum=col.spectrum, n=1000)
  
  # Placing labels
  label.indices <- 1:num.strains
  if (!is.null(path)) {
    pdf(paste0(path,"/Diallel_", pheno.name, "_", cross.type, "_", qtl.num, "qtl.pdf"),
        width = 12, height = 12)    
  }
  par(mfrow = c(num.strains, num.strains), 
      cex = 0.5, 
      oma = c(1, 1, 1, 0), 
      mar = c(1, 3, 3, 1))
  ## CHOOSE SUBSET OF DATA
  for (i in 1:num.strains) {
    for (j in 1:num.strains) {
      if (cross.type == "f2") {
        if (i == j) {
          if (i == 1) {
            if (include.info.plot) {
              infoPlotter(trait = pheno.name, 
                          experiment = cross.type,
                          n = n,
                          spectrum = spectrum, 
                          utility.type = utility.type,
                          x.high = x.high)
            }
            else {
              emptyPlotter(include.off.x = include.off.x)
            }
          }
          else if (i == 2) {
            if (include.var.pie) {
              legendPlotter()
            }
            else {
              emptyPlotter(include.off.x = include.off.x)
            }
          }
          else {
            emptyPlotter(include.off.x = include.off.x)
          }
        }
        else if (i > j) {
          emptyPlotter(include.off.x = include.off.x)
        }      	
        else {
          oneParamPlotter(cross.utility = utility.list$f2[[paste(strains[i], strains[j], sep = "x")]],
                          cross.type="f2", 
                          qtl.perc=median(var.list$f2[[paste(strains[i], strains[j],  sep = "x")]]),
                          qtl.num = qtl.num, 
                          x.high = x.high,
                          hom1.vec = pheno.list$f2$hom1[[paste(strains[i], strains[j],  sep = "x")]],
                          hom2.vec = pheno.list$f2$hom2[[paste(strains[i], strains[j],  sep = "x")]],
                          het.vec = pheno.list$f2$het[[paste(strains[i], strains[j],  sep = "x")]],
                          spectrum = spectrum,
                          include.density = include.density,
                          include.var.pie = include.var.pie,
                          include.bar.plots = include.bar.plots,
                          absolute.max = absolute.max,
                          density.col = density.col,
                          border.col = border.col,
                          median.line.col = median.line.col,
                          include.rank = include.rank,
                          rank = rank.list[["f2"]][[paste(strains[i], strains[j],  sep = "x")]],
                          rank.col = rank.col,
                          ...)
        }
      }	
      else if (cross.type == "bc") {
        if (i == j) {
          if (i == 1) {
            if (include.info.plot) {
              infoPlotter(trait = pheno.name, 
                          experiment = cross.type,
                          n = n,
                          spectrum = spectrum, 
                          utility.type = utility.type,
                          x.high = x.high)
            }
            else {
              emptyPlotter(include.off.x = include.off.x)
            }
          }
          else if (i == 2) {
            if (include.var.pie) {
              legendPlotter()
            }
            else {
              emptyPlotter(include.off.x = include.off.x)
            }
          }
          else {
            emptyPlotter(include.off.x = include.off.x)
          }
        }
        else {
          if (i < j) { ## Upper diag plots
            oneParamPlotter(utility.list$bc1[[paste(strains[i], strains[j],  sep = "x")]],
                            cross.type = "bc", 
                            qtl.perc = median(var.list$bc1[[paste(strains[i], strains[j],  sep = "x")]]),
                            hom1.vec = pheno.list$bc1$hom[[paste(strains[i], strains[j],  sep = "x")]],
                            het.vec = pheno.list$bc1$het[[paste(strains[i], strains[j],  sep = "x")]],
                            qtl.num = qtl.num, 
                            x.high = x.high,
                            back.allele = "A",
                            spectrum = spectrum,
                            absolute.max = absolute.max,
                            include.density = include.density,
                            include.var.pie = include.var.pie,
                            include.bar.plots = include.bar.plots,
                            density.col = density.col,
                            border.col = border.col,
                            median.line.col = median.line.col,
                            include.rank = include.rank,
                            rank = rank.list[["bc1"]][[paste(strains[i], strains[j],  sep = "x")]],
                            rank.col = rank.col,
                            ...)          				
          }
          else { ## Lower diag plots
            oneParamPlotter(utility.list$bc2[[paste(strains[j], strains[i],  sep = "x")]],
                            cross.type = "bc", 
                            qtl.perc=median(var.list$bc2[[paste(strains[j], strains[i],  sep = "x")]]),
                            hom1.vec=pheno.list$bc2$hom[[paste(strains[j], strains[i],  sep = "x")]],
                            het.vec=pheno.list$bc2$het[[paste(strains[j], strains[i],  sep = "x")]],
                            qtl.num = qtl.num, 
                            x.high = x.high,
                            back.allele = "A",
                            spectrum = spectrum,
                            absolute.max = absolute.max,
                            include.density = include.density,
                            include.var.pie = include.var.pie,
                            include.bar.plots = include.bar.plots,
                            density.col = density.col,
                            border.col = border.col,
                            median.line.col = median.line.col,
                            include.rank = include.rank,
                            rank = rank.list[["bc2"]][[paste(strains[j], strains[i],  sep = "x")]],
                            rank.col = rank.col,
                            ...)   				
          }
        }
      }
      else if (cross.type == "rbc1") {
        if (i == j){
          if (i == 1) {
            if (include.info.plot) {
              infoPlotter(trait = pheno.name, 
                          experiment = cross.type,
                          n = n,
                          spectrum = spectrum, 
                          utility.type = utility.type,
                          x.high = x.high)
            }
            else {
              emptyRBC1Plotter()
            }
          }
          else if (i == 2) {
            if (include.var.pie) {
              legendPlotter()
            }
            else {
              emptyRBC1Plotter()
            }
          }
          
          else {
            emptyRBC1Plotter()
          }  
        }
        else {
          if (i < j) { ## Upper diagonal
            oneParamPlotter(utility.list$rbc1_1[[paste(strains[i], strains[j], sep = "x")]],
                            cross.type = "bc", 
                            qtl.perc = median(var.list$rbc1_1[[paste(strains[i], strains[j], sep = "x")]]),
                            hom1.vec = pheno.list$rbc1_1$hom[[paste(strains[i], strains[j], sep = "x")]],
                            het.vec = pheno.list$rbc1_1$het[[paste(strains[i], strains[j], sep = "x")]],
                            qtl.num = qtl.num, 
                            x.high = x.high,
                            back.allele = "A",
                            spectrum = spectrum,
                            absolute.max = absolute.max,
                            include.density = include.density,
                            include.var.pie = include.var.pie,
                            include.bar.plots = include.bar.plots,
                            density.col = density.col,
                            border.col = border.col,
                            median.line.col = median.line.col,
                            include.rank = include.rank,
                            rank = rank.list[["rbc1_1"]][[paste(strains[i], strains[j],  sep = "x")]],
                            rank.col = rank.col,
                            ...)            			
          }
          else { ## Lower diagonal
            oneParamPlotter(utility.list$rbc2_2[[paste(strains[j], strains[i], sep = "x")]],
                            cross.type = "bc", 
                            qtl.perc = median(var.list$rbc2_2[[paste(strains[j], strains[i], sep = "x")]]),
                            hom1.vec = pheno.list$rbc2_2$hom[[paste(strains[j], strains[i], sep = "x")]],
                            het.vec = pheno.list$rbc2_2$het[[paste(strains[j], strains[i], sep = "x")]],
                            qtl.num = qtl.num, 
                            x.high = x.high,
                            back.allele = "A",
                            spectrum = spectrum,
                            absolute.max = absolute.max,
                            include.density = include.density,
                            include.var.pie = include.var.pie,
                            include.bar.plots = include.bar.plots,
                            density.col = density.col,
                            border.col = border.col,
                            median.line.col = median.line.col,
                            include.rank = include.rank,
                            rank = rank.list[["rbc2_2"]][[paste(strains[j], strains[i],  sep = "x")]],
                            rank.col = rank.col,
                            ...)   				
          }
        }
      }
      else if (cross.type == "rbc2") {
        if (i == j) {
          if (i == 1) {
            if (include.info.plot) {
              infoPlotter(trait = pheno.name, 
                          experiment = cross.type, 
                          n = n, 
                          spectrum = spectrum, 
                          utility.type = utility.type,
                          x.high = x.high)
            }
            else {
              emptyRBC2Plotter()
            }
          }
          else if (i == 2) {
            if (include.var.pie) {
              legendPlotter()
            }
            else{
              emptyRBC2Plotter()
            }  
          }
          else {
            emptyRBC2Plotter()
          }
        }
        else {
          if (i < j) {
            oneParamPlotter(utility.list$rbc1_2[[paste(strains[i], strains[j], sep = "x")]],
                            cross.type = "bc", 
                            qtl.perc = median(var.list$rbc1_2[[paste(strains[i], strains[j], sep = "x")]]),
                            hom1.vec = pheno.list$rbc1_2$hom[[paste(strains[i], strains[j], sep = "x")]],
                            het.vec = pheno.list$rbc1_2$het[[paste(strains[i], strains[j], sep = "x")]],
                            qtl.num = qtl.num,
                            x.high = x.high,
                            back.allele = "A",
                            spectrum = spectrum,
                            absolute.max = absolute.max,
                            include.density = include.density,
                            include.var.pie = include.var.pie,
                            include.bar.plots = include.bar.plots,
                            density.col = density.col,
                            border.col = border.col,
                            median.line.col = median.line.col,
                            include.rank = include.rank,
                            rank = rank.list[["rbc1_2"]][[paste(strains[i], strains[j],  sep = "x")]],
                            rank.col = rank.col,
                            ...)              		
          }
          else {
            oneParamPlotter(utility.list$rbc2_1[[paste(strains[j], strains[i], sep = "x")]],
                            cross.type = "bc", 
                            qtl.perc = median(var.list$rbc2_1[[paste(strains[j], strains[i], sep = "x")]]),
                            hom1.vec = pheno.list$rbc2_1$hom[[paste(strains[j], strains[i], sep = "x")]],
                            het.vec = pheno.list$rbc2_1$het[[paste(strains[j], strains[i], sep = "x")]],
                            qtl.num = qtl.num, 
                            x.high = x.high,
                            back.allele = "A",
                            spectrum = spectrum,
                            absolute.max = absolute.max,
                            include.density = include.density,
                            include.var.pie = include.var.pie,
                            include.bar.plots = include.bar.plots,
                            density.col = density.col,
                            border.col = border.col,
                            median.line.col = median.line.col,
                            include.rank = include.rank,
                            rank = rank.list[["rbc2_1"]][[paste(strains[j], strains[i],  sep = "x")]],
                            rank.col = rank.col,
                            ...)   				
          }
        }
      }
      if (j %in% label.indices & i == 1) {
        if (include.biparent.labels) {
          mtext(paste(ifelse(is.null(strains.relabel), strains[j], strains.relabel[j]), "(B)"), 
                side = 3, cex = label.cex, padj = label.padj)
        }
        else {
          mtext(ifelse(is.null(strains.relabel), strains[j], strains.relabel[j]), 
                side = 3, cex = label.cex, padj = label.padj)
        }
      }
      if (i %in% label.indices & j == 1) {
        if (include.biparent.labels) {
          mtext(paste(ifelse(is.null(strains.relabel), strains[i], strains.relabel[i]), "(A)"), 
                side = 2, cex = label.cex, padj = label.padj)
        }
        else {
          mtext(ifelse(is.null(strains.relabel), strains[i], strains.relabel[i]), 
                side = 2, cex = label.cex, padj = label.padj)
        }
      }
    }
  }
  if (!is.null(path)) {
    dev.off()
  }
}


################## Component plots of Moonrise plot
make.spectrum <- function(col.range, 
                          col.spectrum, 
                          n = 1000) {
  if (is.null(col.range)) {
    if (col.spectrum == "gray") {
      spectrum <- gray(level=n:1/n)
    }
    else {
      spectrum <- do.call(what=eval(parse(text=paste0("colorRamps::", col.spectrum))), args=list(n=n))
    }
  }
  else {
    spectrum <- colorRampPalette(col.range)
    spectrum <- spectrum(n)
  }
  return(spectrum)
}

emptyPlotter <- function(include.off.x = FALSE, ...){
  if (!include.off.x) {
    this.col <- "white"
  }
  else {
    this.col <- "black"
  }
  plot(x = 1, y = 1, cex = 50, pch = 4, xlab = "", ylab = "", xlim = c(0, 1), ylim = c(0, 1),
       frame.plot = FALSE, xaxt = "n", yaxt = "n", col = "white")
  lines(x = c(0, 1), y = c(0, 1), col = this.col)
  lines(x = c(0, 1), y = c(1, 0), col = this.col)
}

emptyF2Plotter <- function(background){
  plot(x = 1, y = 1, cex = 1, pch = 4, xlab = "", ylab = "", xlim = c(-5, 5), ylim = c(-5,5),
       frame.plot = FALSE, xaxt = "n", yaxt = "n", col = "white")
  text(0, 3, "F2", cex = 4)
  
  text(0, -2, labels = paste(background, " (A)", sep = ""), cex = 2)
}

emptyBCPlotter <- function(background){
  plot(x=1,y=1, cex = 1, pch ="", xlab = "", ylab = "", xlim = c(-10, 10), ylim = c(-10,10),
       frame.plot = FALSE, xaxt = "n", yaxt = "n", col = "white")
  arrows(5, 0, -5, 0, length = 0.1)
  text(0, -3, labels = paste("Background: ", background, " (A)", sep = ""), cex = 1.5)
}

emptyRBC1Plotter <- function(){
  plot(x=1,y=1, cex = 1, pch ="", xlab = "", ylab = "", xlim = c(-10, 10), ylim = c(-10,10),
       frame.plot = FALSE, xaxt = "n", yaxt = "n", col = "white")

  arrows(3, 3, -3 , 3, length = 0.1)
  text(0, -1, "Maternal Strain", cex = 1.5)
}

emptyRBC2Plotter <- function(){
  plot(x = 1, y = 1, cex = 1, pch ="", xlab = "", ylab = "", xlim = c(-10, 10), ylim = c(-10,10),
       frame.plot = FALSE, xaxt = "n", yaxt = "n", col = "white")
  
  arrows(3, 3, -3 , 3, length = 0.1)
  text(0, -1, "Paternal Strain", cex = 1.5)
}

legendPlotter <- function(){
  plot(NA, xlim = c(0,1), ylim = c(0,1), xlab = "", ylab = "", frame = FALSE, xaxt = "n", yaxt = "n")
  
  legend("center", legend = c("QTL", "Noise"), 
         fill = c("white", "gray50"), border = "black", bty = "n", 
         title = "Variance attributable to", cex = 1.3, pt.cex = 1.3) 
}

infoPlotter <- function(trait,
                        experiment,
                        n, 
                        spectrum, 
                        x.high = 1,
                        utility.type){
  plot(NA, xlim = c(0,1), ylim = c(0,1), xlab = "", ylab = "", frame = FALSE, xaxt = "n", yaxt = "n")
  
  if (experiment == "f2") { this.experiment <- "F2" }
  if (experiment == "bc") { this.experiment <- "BC" }
  if (experiment == "rbc1") { this.experiment <- "RBC (A maternal)" }
  if (experiment == "rbc2") { this.experiment <- "RBC (A paternal)" }
  
  text(x=0.1, y=0.9, labels=paste("Cross type:", this.experiment), adj=0, cex=1.2)
  text(x=0.1, y=0.8, labels=paste("Trait:", trait), adj=0, cex=1.2)
  text(x=0.1, y=0.7, labels=paste("Utility:", utility.type), adj=0, cex=1.2)
  if (utility.type == "power") {
    text(x=0.1, y=0.6, labels=paste("QTL number:", x.high), adj=0, cex=1.2)
    text(x=0.1, y=0.5, labels=paste("Number of mice:", n), adj=0, cex=1.2)
  }
  
  barplot(height=rep(0.1, length(spectrum)), width=1/length(spectrum), density=1000,
          angle=90, col=spectrum, 
          border=FALSE, space=FALSE, axes=FALSE, add=TRUE)
  axis(1, at=c(0,1), labels=c(0, x.high), tick=TRUE, cex.axis=1.2)
  text(x=0.5, 0.2, labels="Posterior mean utility", cex=1.3)
}

#' Gradient scale for posterior utility from DIDACT grid plot.
#' 
#' This function produces a gradient scale that can be used with a DIDACT grid plot.
#' 
#' @param spectrum DEFAULT: make.spectrum(c("white", "black"), n = 100). The color spectrum to be used in the gradient.
#' @param mode DEFAULT: "horizontal". Specifies whether the gradient is horizontally or vertically oriented.
#' @param utility.type DEFAULT: "power". Specifies whether the utility function is power or contrasts.
#' @export make.gradient.scale
#' @examples make.gradient.scale()
make.gradient.scale <- function(spectrum = make.spectrum(c("white", "black"), n = 100),
                                mode = c("horizontal", "vertical"),
                                utility.type = c("power", "contrasts")) {
  mode <- mode[1]
  utility.type <- utility.type[1]
  
  if (mode == "horizontal") {
    barplot(height = rep(0.1, length(spectrum)), 
            width = 1/length(spectrum), 
            density = 1000, 
            angle = 90, 
            col = spectrum,
            border = FALSE, 
            space = FALSE, 
            axes = FALSE)
    if (utility.type == "power") {
      axis(1, at = c(0, 1))
      mtext(side = 1, text = "Posterior power", line = 3, cex = 1.5)
    }
    else {
      axis(1, at = c(0, 1), labels = c("low", "high"))
      mtext(side = 1, text = "Posterior contrast", line = 3, cex = 1.5)
    }
  }
  else {
    barplot(height = rep(0.1, length(spectrum)), 
            width = 1/length(spectrum), 
            density = 1000, 
            angle = 0, 
            col = spectrum,
            border = FALSE, 
            space = FALSE, 
            axes = FALSE,
            horiz = TRUE)
    if (utility.type == "power") {
      axis(4, las = 1, at = c(0, 1))
      mtext(side = 1, text = "Posterior power", cex = 1.5, line = 2)
    }
    else {
      axis(4, las = 1, at = c(0, 1), labels = c("low", "high"))
      mtext(side = 1, text = "Posterior contrast", cex = 1.5, line = 2)
    }
  }
}

f2boxPlotter <- function(hom1.vec, 
                         hom2.vec, 
                         het.vec, 
                         y.max, 
                         x.max,
                         border.col){
  max.box.y <- max(hom1.vec, hom2.vec, het.vec, na.rm=TRUE)
  min.box.y <- min(hom1.vec, hom2.vec, het.vec, na.rm=TRUE)
  box.y.range <- max.box.y - min.box.y
  # Scaling and rescaling
  hom1.vec.sc <- (hom1.vec - min.box.y)*(y.max*(3/5)/(max.box.y-min.box.y)) + y.max*(5/4)
  hom2.vec.sc <- (hom2.vec - min.box.y)*(y.max*(3/5)/(max.box.y-min.box.y)) + y.max*(5/4)
  het.vec.sc <- (het.vec - min.box.y)*(y.max*(3/5)/(max.box.y-min.box.y)) + y.max*(5/4)
  mid.x <- (2/5)*(3/8)*x.max
  shift.x <- (3/4)*mid.x
  boxplot(hom1.vec.sc, 
          het.vec.sc, 
          hom2.vec.sc,
          width=rep(1, 3), 
          boxwex=(1/15)*x.max,
          border=border.col,
          outline=FALSE, 
          col="white",
          add=TRUE, 
          at=c(mid.x-shift.x, mid.x, mid.x+shift.x), names=NA, yaxt="n", xaxt="n")
  text(x=mid.x-shift.x, y=y.max*(5/4), labels="A/A", cex=0.8, col=border.col)
  text(x=mid.x, y=y.max*(5/4), labels="A/B", cex=0.8, col=border.col)
  text(x=mid.x+shift.x, y=y.max*(5/4), labels="B/B", cex=0.8, col=border.col)
  text(x=mid.x+(1/2)*shift.x, y=2*y.max-(1/15)*y.max, labels="Phenotypes", cex=1.1, col=border.col)
}

bcboxPlotter <- function(hom.vec, 
                         het.vec, 
                         y.max, 
                         x.max, 
                         back.allele="A",
                         border.col){
  max.box.y <- max(hom.vec, het.vec, na.rm=TRUE)
  min.box.y <- min(hom.vec, het.vec, na.rm=TRUE)
  box.y.range <- max.box.y - min.box.y
  # Scaling and rescaling
  if (back.allele == "A") {
    hom1.vec.sc <- (hom.vec - min.box.y)*(y.max*(4/7)/(max.box.y-min.box.y)) + y.max*(4/3)
    het.vec.sc <- (het.vec - min.box.y)*(y.max*(4/7)/(max.box.y-min.box.y)) + y.max*(4/3)
    hom2.vec.sc <- rep(NA, length(hom.vec))
  }
  else {
    hom1.vec.sc <- rep(NA, length(hom.vec))
    het.vec.sc <- (het.vec - min.box.y)*(y.max*(4/7)/(max.box.y-min.box.y)) + y.max*(4/3)
    hom2.vec.sc <- (hom.vec - min.box.y)*(y.max*(4/7)/(max.box.y-min.box.y)) + y.max*(4/3)
  }

  mid.x <- (2/5)*(3/8)*x.max
  shift.x <- (3/4)*mid.x
  boxplot(hom1.vec.sc, 
          het.vec.sc,
          hom2.vec.sc,
          width=rep(1, 3), 
          boxwex=(1/15)*x.max, 
          outline=FALSE, 
          col="white",
          add=TRUE, 
          border=border.col,
          at=c(mid.x-shift.x, mid.x, mid.x+shift.x), names=NA, yaxt="n", xaxt="n")
  text(x=mid.x-shift.x, y=y.max*(5/4), labels="A/A", cex=0.8, col=border.col)
  text(x=mid.x, y=y.max*(5/4), labels="A/B", cex=0.8, col=border.col)
  text(x=mid.x+shift.x, y=y.max*(5/4), labels="B/B", cex=0.8, col=border.col)
  text(x=mid.x+(1/2)*shift.x, y=2*y.max-(1/15)*y.max, labels="Phenotypes", cex=1.1, col=border.col)
}

oneParamPlotter <- function(cross.utility, 
                            cross.type, 
                            qtl.perc, 
                            qtl.num=1,
                            x.high,
                            cross.label1="", 
                            cross.label2="",
                            hom1.vec, 
                            hom2.vec=NULL, 
                            het.vec, 
                            back.allele=NULL,
                            include.x.axis=FALSE,
                            spectrum,
                            absolute.max=NULL,
                            include.density,
                            include.var.pie,
                            include.bar.plots,
                            include.rank,
                            rank=NULL,
                            rank.col="red",
                            density.col="gray",
                            border.col="black",
                            median.line.col="black",
                            ...){
  post.mean <- mean(cross.utility)
  post.median <- median(cross.utility)
  bgcolor <- spectrum[round((post.mean/x.high)*length(spectrum))]
  n <- length(cross.utility)
  if (is.null(absolute.max)) {
    max.y <- max(hist(cross.utility, plot=FALSE, breaks=seq(0, x.high, length.out=20))$density)
  }
  else {
    max.y <- absolute.max
  }
  
  min.y <- 0
  
  ## Adjusting plot window for widgets
  if (include.var.pie | include.bar.plots) {
    y.max <- 1*(max.y - min.y) + max.y
  }
  else {
    y.max <- max.y
  }

  plot(1, 1, xlim = c(0, x.high), 
       ylim = c(0, y.max), 
       lwd = 3, xlab = "", main = "", frame.plot = FALSE, ylab = "", yaxt = "n", xaxt = "n", col = "white")
  rect(par("usr")[1], par("usr")[3], par("usr")[2], par("usr")[4], col = bgcolor)
  y.range <- max.y - min.y
  qtl <- round(qtl.perc/qtl.num, 1)
  total.qtl <- round(qtl.perc, 1)
  total <- round(100 - total.qtl, 1)

  if (include.var.pie) {
    if (qtl.num == 1) {
      mapplots::add.pie(z = c(qtl/100, total/100), 
                        labels = NA, 
                        x = 0.5*x.high, 
                        y = 0.75*y.max, 
                        radius = (3/10)*(9/10)*y.range, 
                        col = (c("white", "gray50")), 
                        cex = 1.3, 
                        label.dist = 1.2, 
                        border = border.col)
    }
    else {
      excess.qtl <- round(qtl*(qtl.num-1), 1)
      mapplots::add.pie(z = c(qtl/100, excess.qtl/100, total/100), 
                        labels = NA, 
                        x = 0.5*x.high, 
                        y = 0.75*y.max, 
                        radius = (3/10)*(9/10)*y.range, 
                        col = (c("white", "white", "gray50")), 
                        cex = 1.3, 
                        label.dist = 1.2, 
                        border = border.col)  
    }
    legend(x = 0.6*x.high,
           y = y.max+0.04*y.max,
           legend = c(qtl, total), 
           fill = c("white", "gray50"), 
           title = "% Variance", 
           border = border.col, 
           bty = "n", 
           text.col = border.col, 
           cex = 1.1)
  }
  if (include.bar.plots) {
    if (cross.type == "f2") {
      f2boxPlotter(hom1.vec = hom1.vec, 
                   hom2.vec = hom2.vec, 
                   het.vec = het.vec, 
                   y.max = max.y, 
                   x.max = x.high,
                   border.col = border.col)
    }
    else if (cross.type == "bc") {
      bcboxPlotter(hom.vec = hom1.vec, 
                   het.vec = het.vec, 
                   y.max = max.y, 
                   x.max = x.high, 
                   back.allele = back.allele,
                   border.col = border.col)
    }
  }
  if (include.density) {
    hist(cross.utility, 
         col = density.col, 
         breaks=seq(0, x.high, length.out = 20), 
         ylim = c(0, y.max), 
         xlim = c(0, x.high), 
         add = TRUE, 
         freq = FALSE, 
         border = border.col)
    lines(x = c(post.median, post.median), 
          y = c(0, max.y), 
          lty = 5, 
          lwd = 2, 
          col = median.line.col)
  }
  if (cross.label1 != "" & cross.label2 != "") {
    text(1, median(dens$y), paste(cross.label1, 'x', cross.label2, sep=""))
  }
  if (include.x.axis) {
    axis(1, at=0:x.high, labels=TRUE, tick=TRUE, cex.axis=1.2)
  }
  if (include.rank) {
    #legend("topright", legend=rank, bty="n", text.col=rank.col, cex=2.5, text.font=1)
    TeachingDemos_shadowtext(x=x.high*0.8, y=y.max*0.8, labels=rank, col=rank.col, cex=5, r=0.15)
  }
}

TeachingDemos_shadowtext <- function(x, y=NULL, labels, col='white', bg='black',
                                     theta= seq(pi/4, 2*pi, length.out=8), r=0.1, ...) {
  xy <- xy.coords(x,y)
  xo <- r*strwidth('A')
  yo <- r*strheight('A')
  
  for (i in theta) {
    text(xy$x + cos(i)*xo, xy$y + sin(i)*yo, labels, col=bg, ...)
  }
  text(xy$x, xy$y, labels, col=col, ...) 
}

#' Info plot that can be included in the diagonal of the DIDACT grid plot.
#' 
#' This function produces the information plot that is included in the DIDACT grid plot. This includes
#' informtion on the trait, cross type, number of QTL, sample size (for power utility), and utility ramp.
#' 
#' @param trait A string that is included as the trait or phenotype.
#' @param experiment DEFAULT: "f2". A string that is included as the cross type. Expected values include "f2" and "bc".
#' @param utility.type DEFAULT: "power". A string that is included as the utility type. Expected values are "power" and "contrasts".
#' @param n An integer that is included as the sample size for power calculation.
#' @param col.range DEFAULT: c("white", "black"). If specified, will create a color spectrum scale between the
#' two colors included.
#' @param col.spectrum DEFAULT: "blue2red". Use pre-specified spectrum. Options include "blue2red", "gray", 
#' "green2red", and "blue2green".
#' @param x.high DEFAULT: 1. The high value for the utility ramp. For power, 1 should be the maximum value.
#' @export make.big.info.plot
#' @examples make.big.info.plot()
make.big.info.plot <- function(trait,
                               experiment = c("f2", "bc"),
                               utility.type = c("power", "contrasts"),
                               n,
                               col.range = c("white", "black"),
                               col.spectrum = c("blue2red", "gray", "green2red", "blue2green"),
                               x.high = 1){
  
  experiment <- experiment[1]
  utility.type <- utility.type[1]
  
  ## Setting color spectrum
  spectrum <- make.spectrum(col.range = col.range, 
                            col.spectrum = col.spectrum, 
                            n = 1000)
  
  infoPlotter(trait = trait,
              experiment = experiment,
              n = n,
              spectrum = spectrum,
              utility.type = utility.type,
              x.high = x.high)
}

#' Single DIDACT square plot of posterior utility.
#' 
#' This function takes posterior samples of utilities from evaluate.experiments() and produces a single 
#' square plot of the posterior utilities for specified cross.
#'
#' @param cross The specific cross of strains to plot. Expects a string in the format of "AxB".
#' @param cross.type The type of cross to evaluate. Expects "f2", "bc", "rbc1", and "rbc2".
#' @param didact.object Samples of posterior utility produced by evaluate.experiments().
#' @param utility.type DEFAULT: "power". The posterior utility to be plotted. Currently the options are "power"
#' and "contrasts". Power is more appropriate for Mendelian-like phenotypes. Contrasts make less assumptions.
#' @param col.range DEFAULT: c("white", "black"). If specified, will create a color spectrum scale between the
#' two colors included.
#' @param col.spectrum DEFAULT: "blue2red". Use pre-specified spectrum. Options include "blue2red", "gray", 
#' "green2red", and "blue2green".
#' @param include.var.pie DEFAULT: TRUE. If TRUE, pie chart of phenotypic variance is included in square.
#' @param include.bar.plots DEFAULT: TRUE. If TRUE, box plots of posterior genotype class phenotypes are 
#' included in square.
#' @param include.density DEFAULT: TRUE. If TRUE, histogram of posterior density is included in square.
#' @param back.allele DEFAULT: "A". Specify which founder is backcrossed in a backcross.
#' @export make.single.cross.plot
#' @examples make.single.cross.plot()
make.single.cross.plot <- function(cross,
                                   cross.type,
                                   didact.object,
                                   utility.type = c("power", "contrasts"),
                                   col.range = c("white", "black"),
                                   col.spectrum = c("blue2red", "gray", "green2red", "blue2green"),
                                   include.var.pie = TRUE,
                                   include.bar.plots = TRUE,
                                   include.density = TRUE,
                                   back.allele = "A",
                                   ...){
  col.spectrum <- col.spectrum[1]
  utility.type <- utility.type[1]
  ## Setting color spectrum
  spectrum <- make.spectrum(col.range = col.range, col.spectrum = col.spectrum, n = 1000)

  if (utility.type == "power") {
    this.cross.utility <- didact.object$power[[cross.type]][[cross]]
  }
  else if (utility.type == "contrasts") {
    this.cross.utility <- didact.object$var[[cross.type]][[cross]]
  }
  
  this.qtl.perc <- median(didact.object$var[[cross.type]][[cross]])
  this.het.vec <- didact.object$pheno[[cross.type]][["het"]][[cross]]
  if (cross.type == "f2") {
    this.hom1.vec <- didact.object$pheno[[cross.type]][["hom1"]][[cross]]
    this.hom2.vec <- didact.object$pheno[[cross.type]][["hom2"]][[cross]]
  }
  else if (cross.type == "bc") {
    this.hom1.vec <- didact.object$pheno[[cross.type]][["hom"]][[cross]]
    this.hom2.vec <- NULL
  }
  qtl.num <- didact.object$qtl.num
  if (utility.type == "power") {
    x.high <- qtl.num
  }
  else if (utility.type == "contrasts") {
    #x.high <- max(sapply(1:length(didact.object$var), function(y) max(sapply(1:length(didact.object$var[[y]]), function(x) max(didact.object$var[[y]][[x]])))))
    x.high <- 100
  }
  oneParamPlotter(cross.utility = this.cross.utility,
                  cross.type = cross.type,
                  qtl.perc = this.qtl.perc,
                  qtl.num = qtl.num,
                  x.high = x.high,
                  hom1.vec = this.hom1.vec,
                  hom2.vec = this.hom2.vec,
                  het.vec = this.het.vec,
                  spectrum = spectrum,
                  include.var.pie = include.var.pie,
                  include.bar.plots = include.bar.plots,
                  include.density = include.density,
                  include.rank = FALSE,
                  back.allele = back.allele,
                  ...)
  if (utility.type == "power") {
    axis(side = 1, at = c(0, 1), labels = c("low", "high"))
  }
  else {
    axis(side = 1, at = c(0, 100), labels = c("low", "high"))
  }
  mtext(side = 1, text = "Posterior utility", line = 2.5)
}

calc.ranks <- function(utility.list,
                       cross.type=c("f2", "bc", "rbc1", "rbc2")) {
  rank.list <- list()
  
  rank.list$f2 <- as.list(rank(-unlist(lapply(utility.list$f2, function(x) mean(x)))))

  rank.vec <- rank(-unlist(lapply(c(utility.list$bc1, utility.list$bc2), function(x) mean(x))))
  rank.list$bc1 <- as.list(rank.vec[1:28])
  rank.list$bc2 <- as.list(rank.vec[29:56])

  rank.vec <- rank(-unlist(lapply(c(utility.list$rbc1_1, utility.list$rbc2_2), function(x) mean(x))))
  rank.list$rbc1_1 <- as.list(rank.vec[1:28])
  rank.list$rbc2_2 <- as.list(rank.vec[29:56])
  
  rank.vec <- rank(-unlist(lapply(c(utility.list$rbc1_2, utility.list$rbc2_1), function(x) mean(x))))
  rank.list$rbc1_2 <- as.list(rank.vec[1:28])
  rank.list$rbc2_1 <- as.list(rank.vec[29:56])
  
  return(rank.list)
}
```
