## Supplementary material for "Bayesian decision theoretic design of two-founder experimental crosses given diallel data": README.md

DIDACT
======

**D**iallel **I**nformed **D**ecision theoretic **A**pproach for **C**rosses **T**ool

DIDACT extends the [BayesDiallel model](http://valdarlab.unc.edu/software/bayesdiallel/BayesDiallel.html), a hierarchical Bayesian model of a diallel experiment, to posterior samples of specified decision theoretic utility functions that is valuable in selecting and designing follow-up experiments.

**Example:** Quantitative trait loci (QTL) mapping is not possible with a diallel experiment because it consists of F0s and F1s. No recombination events have remixed the founder haplotypes, allowing an effect on some phenotype to be attributed to a segregating portion of the haplotype. However, there is still genetic information on the strain/line level. DIDACT provides an approach for attenuating the signal and uncertainty estimated from the strain-level diallel data through to follow-up mapping experiments.

Example data is included in the package. The following code serves as a simple vignette for using the package for now.
```r
library(devtools)
install_github("gkeele/DIDACT")
library(DIDACT)

## advia.dat is a diallel data set pre-loaded in DIDACT

chgb.par <- diallel.gibbs(phenotype=advia.dat$cHGB, sex=as.numeric(advia.dat$is.female=="F"),
			  mother.str=advia.dat[,1], father.str=advia.dat[,2], n.iter=1000, burn.in=10000,
			  use.constraint=TRUE)
caterpillar.plot(chgb.par)
chgb.par.exp = evaluate.experiments(chgb.par, n=500, qtl.num=1)
diallelPlotter(results= chgb.par.exp, cross.type="f2", include.bar.plots = FALSE, include.density = FALSE, include.rank = TRUE, include.info.plot = FALSE, utility.type = "contrasts")
diallelPlotter(results= chgb.par.exp, cross.type="bc", include.bar.plots = FALSE, include.density = FALSE, include.rank = TRUE, include.info.plot = FALSE, utility.type = "contrasts")
diallelPlotter(results= chgb.par.exp, cross.type="rbc1", include.bar.plots = FALSE, include.density = FALSE, include.rank = TRUE, include.info.plot = FALSE, utility.type = "contrasts")
diallelPlotter(results= chgb.par.exp, cross.type="rbc2", include.bar.plots = FALSE, include.density = FALSE, include.rank = TRUE, include.info.plot = FALSE, utility.type = "contrasts")
```
